# The coast-wide collapse in marine survival of west coast Chinook and steelhead: slow-moving catastrophe or deeper failure?

**DOI:** 10.1101/476408

**Authors:** David W. Welch, Aswea D. Porter, Erin L. Rechisky

## Abstract

Accelerating decreases in survival are evident for northern Hemisphere salmon populations. We collated smolt survival and smolt-to-adult (marine) survival data for all regions of the Pacific coast of North America excluding California to examine the forces shaping salmon returns. A total of 3,055 years of annual survival estimates were available for Chinook (*Oncorhynchus tshawytscha*) and steelhead (*O. mykiss*). This dataset provides a fundamentally different perspective on west coast salmon conservation problems from the previously accepted view. We found that marine survival collapsed over the past half century by a factor of at least 4-5 fold to similar low levels (~1%) for most regions of the west coast. The size of the decline is too large to be compensated by freshwater habitat remediation or cessation of harvest, and too large-scale to be attributable to specific anthropogenic impacts such as dams in the Columbia River or salmon farming in British Columbia. Within the Columbia River, both smolt survivals during downstream migration in freshwater and adult return rates (SARs) of Snake River populations, often singled out as exemplars of poor survival, appear unexceptional and are in fact higher than estimates reported from other regions of the west coast lacking dams. Formal Columbia River rebuilding targets of 2-6% SARs may therefore be unachievable if regions with nearly pristine freshwater conditions also fail to achieve these targets. Finally, we present case studies demonstrating that the historical response to evidence that the salmon problems are primarily ocean-related was to re-emphasize freshwater actions and to stop work on ocean issues. With ocean temperatures forecast to increase far further, the failure of management to identify the drivers of salmon collapse and respond appropriately suggest that the future of most west coast salmon populations is bleak.

## Introduction

The total abundance of salmon in the North Pacific has now reached record levels [2-4]; however, a dramatic contrast in the winners and losers is obscured by this milestone. Most of the increased abundance is in the lowest valued species (pink *Oncorhynchus gorbuscha* and chum *O*. *keta* salmon) in far northern regions, at least partly due to major efforts at ocean ranching of these two species [4]. In contrast while essentially all west coast North American Chinook (*O. tshawytscha*) populations (including Alaska) are now performing poorly with dramatically reduced productivity [6]. The situation is similar for most southern populations of steelhead (*O. mykiss*) [7], coho (*O. kisutch*) [8, 9], and sockeye (*O. nerka*) [10-13]. These poorly performing species are of higher economic value and the preferred focus of First Nations, sport, and many commercial fisheries.

The historical geographic pattern of declines in salmon abundance (greatest problems in the south, least to the north) were originally assumed to reflect a freshwater anthropogenic cause because of the greater degree of terrestrial (i.e., freshwater) habitat modification in the more populous southern regions of the west coast [14, 15]. The growing appreciation of ocean climate change [16-18] has brought a greater awareness of the role of the ocean in influencing salmon survival. As Ryding and Skalski [19] noted almost two decades ago, “*It is becoming increasingly clear that understanding the relationship between the marine environment and salmon survival is central to better management of our salmonid resources*” (p. 2374).

Unfortunately, our scientific understanding of the events occurring in the marine phase remains severely limited, so there has been little change in management strategy beyond the essential first step of reducing harvest rates in the face of falling marine survival. The recent recognition of the decline in Chinook returns across essentially all of Alaska [20-22] and the Canadian portion of the Yukon River [23], where anthropogenic freshwater habitat impacts are generally negligible, is another example of how simple explanations looking at freshwater habitat changes are potentially flawed. If freshwater habitat disruption across this vast swathe of relatively pristine territory is severe enough to seriously impact salmon productivity, then there is little hope that freshwater habitat in more southern regions can be “fixed” to support a newly productive environment for salmon.

The same widespread problem of declining survival is also evident for other diadromous species. Both eulachon [24] and lamprey [25] have undergone sharp unexplained declines along the Pacific west coast of North America. In the Atlantic Ocean, both Atlantic salmon [26] and eels [27, 28] are similarly in sharp decline. In the case of eels, eulachon, and lamprey, the authors attribute the problem to likely marine-related factors, not freshwater. This point is particularly persuasive for eulachon because of the very short freshwater phase [24].

In this paper, we collate Chinook and steelhead time series for the west coast of North America (excluding California) to look at patterns in survival: (1) freshwater survival of smolts during the downstream migration phase and (2) smolt-to-adult return rates (SARs). The SAR is the three-fold product of freshwater smolt survival during downstream migration multiplied by the marine survival experienced over 2-3 years in the ocean multiplied by freshwater survival during the final upstream migration by the returning adults to the final enumeration point. (Depending upon the specific dataset, adult abundance may be enumerated prior to actual arrival at the spawning grounds; see Methods). In particular, given the very poor perceived returns of salmon to the Snake River, many of our analyses compare regional survival to that of Snake River stocks. We use the term SAR and marine survival interchangeably because, as we will demonstrate, the majority of the SAR is determined in the ocean.

For the downstream (freshwater) smolt survival analysis, 46 Chinook and 44 steelhead time series were collated, comprising 531 annual estimates of survival (see Methods). For the SAR comparison, 101 Chinook time series and 50 steelhead time series were available (Fig. 1) which equate to 1,729 Chinook and 795 annual steelhead SAR estimates. Altogether these datasets total 3,055 years of salmon monitoring— clearly, an enormous effort that likely sums to multiple billions of dollars. As the breakdown by regime periods will demonstrate, the tremendous increase in resources devoted to survival monitoring as salmon returns have dwindled over time has perhaps provided less actual insight into mechanisms (as opposed to numbers) than might be hoped, a theme we return to in the Discussion.

**Figure 1.**
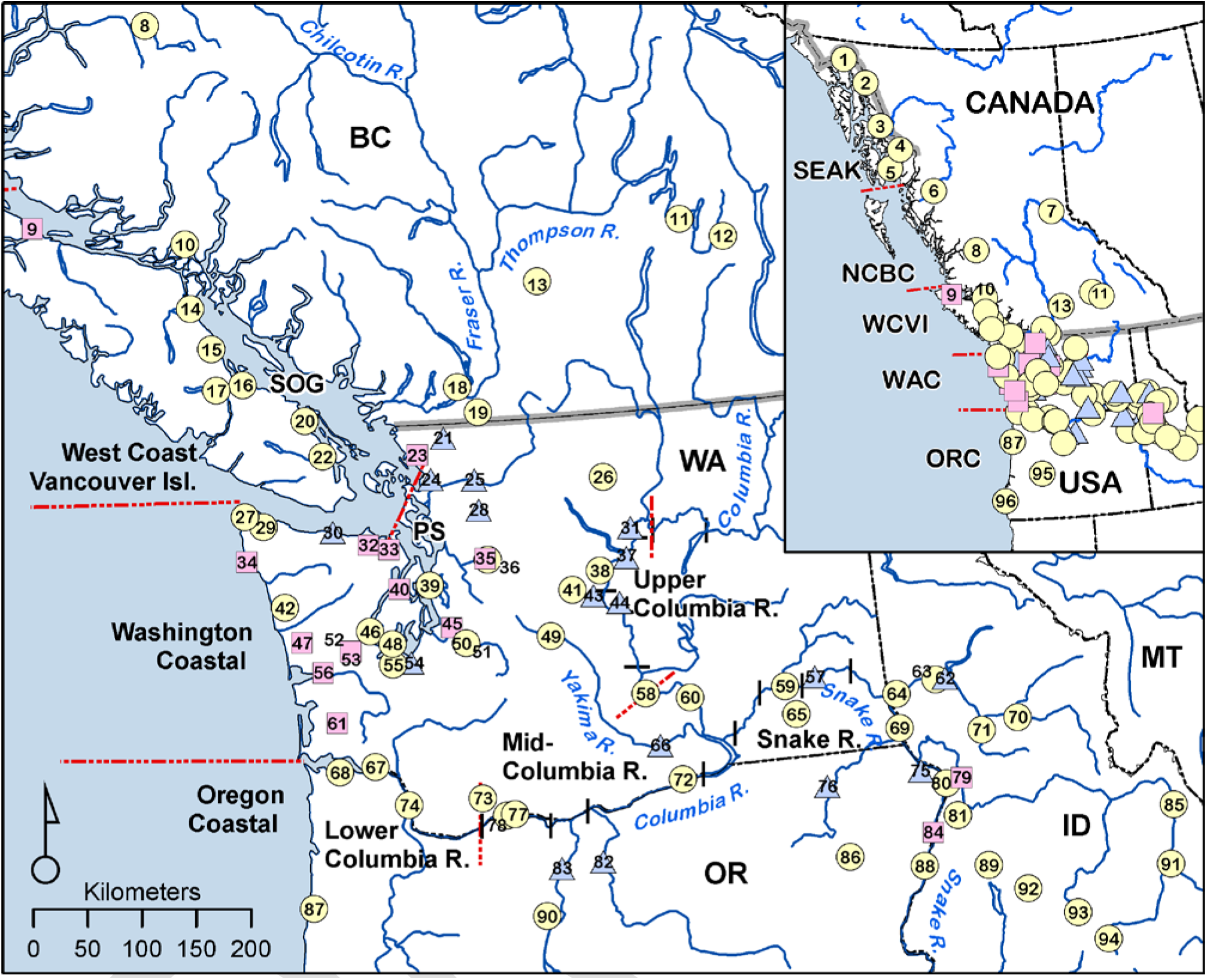
Map of salmon survival time series used in the analyses. Numbers inside symbols are keyed to the populations in Supplementary Table S1; yellow circles indicate Chinook populations, pink squares indicate steelhead, and blue triangles indicate a location with data for both species. Acronyms: SEAK (SE Alaska/Northern British Columbia Transboundary Rivers); NCBC (North-Central British Columbia); WCVI (West Coast Vancouver Island); WAC (Washington Coastal); ORC (Oregon Coastal); SOG (Strait of Georgia); PS (Puget Sound).

The passive integrated transponder (PIT) tag SAR estimates for Chinook and steelhead are specific to the Columbia River Basin and are reported by the Fish Passage Center, most recently by [5]. Estimates reported in an earlier paper by Raymond [1, 29] which predate PIT tag estimates for Columbia River basin Chinook and steelhead were also included. The primary data source for the coded wire tag (CWT) based SAR time series for Chinook used in this analysis is the official survival estimates submitted by various State and Federal government agencies to the Pacific Salmon Commission under the terms of the US-Canada Salmon Treaty. These data include SAR estimates from OR, WA, BC, and AK. For Washington State steelhead outside the Columbia River basin, SARs were collected and reported by Kendall et al [7] for Puget Sound, as well as a number of locations along the coast of Juan de Fuca Strait and the outer (western) WA coast. In BC, SARs are only available for one steelhead population (Keogh River). We are unaware of additional steelhead SAR data for Alaska or coastal Oregon rivers. Individual time series ranged between 2-39 years for Chinook, and 2-37 year for steelhead. (Datasets comprised of only a single year of data were excluded).

What are acceptable levels of salmon survival? For much of the west coast outside of the Columbia River basin, formal recovery targets (SARs) have not been specified, although it is clear in all regions that historical levels of productivity would be greatly preferred to current return rates. (And, to foretell an underlying theme to the paper, current SAR levels may in fact be much preferred to what climate change has in store for salmon in the future). In the extensively dammed Columbia River basin, the Northwest Power and Conservation Council’s Fish and Wildlife Program (NPCC) set rebuilding targets for SARs at 2%-6% ([5], p. 4), roughly the survival observed in the 1960s prior to the completion of the 8-dam Federal Columbia River Power System (FCRPS) [29, 30]. The sharp decrease in salmon returns to the Columbia River (and most particularly the Snake River) after the completion of the final Snake River dam in the mid-1970s was widely assumed to be due to the construction of the dams, and great effort has therefore been devoted to improving in-river smolt survival since that time. For this reason, we have chosen to contrast survival in other geographic regions to that of the Snake River as an objective standard of “poor” survival.

The NPCC SAR objectives did not specify the points in the life cycle where Chinook smolt and adult numbers should be determined. However, one extensive analysis for Snake River spring/summer Chinook was based on SARs calculated as adult and jack returns to the uppermost dam encountered in the migration path [31]: “*Median SARs must exceed 4% to achieve complete certainty of meeting the 48-year recovery standard, while … A median of greater than 6% is needed to meet the 24-year survival standard with certainty*” (p. 41). With most current Columbia River basin SARs on the order of ca. 1%, migratory-phase life-cycle survival would have to increase 200%-600% (two-to six-fold) to meet these targets. It is unclear whether this level of rebuilding is actually possible for reasons that we discuss later.

Unfortunately, as Chinook and steelhead stocks continue to dwindle, progress on addressing and incorporating ocean impacts on salmon dynamics has been slow, perhaps due to a combined lack of understanding about how to address marine survival issues and to pessimism about whether improved understanding of the marine phase can advance conservation. Therefore, lastly we review two case studies which show that even when the overriding role of marine survival is identified, there is still a strong predilection to preferentially identify freshwater factors to study and manipulate. This has resulted in both the failure to directly address the marine survival problem and a rather uncritical approach that too readily identifies widely accepted freshwater stressors as being responsible for the problems evident in specific populations. In our view, a large part of the difficulty lies in some of the fundamental underlying assumptions that the fisheries community makes about the nature of the core problem. Because these assumptions are part of our training and professional ethos, they are difficult to recognize or question. Nevertheless, given the widespread geographic range and magnitude of the collapse in survival that is now evident, we view it as urgent that assumptions about causative agents be carefully assessed.

## Results

### Freshwater (downstream) smolt survival

To separate and assess what are typical freshwater survival levels for smolts migrating downstream, we collated the published studies for west coast North American rivers excluding California (See Methods for a more detailed summary and Table S1 for reported estimates). We used these data to make regional comparisons of smolt survival and survival scaled by distance travelled during the downstream freshwater migration to the sea.

Within the Columbia River basin, survival estimates for a range of stocks and river reaches are available, although the majority are for survival through the hydrosystem (dammed segment of the river). For yearling Chinook, smolt survival estimates varied considerably between grouping categories (Fig 2; center column, top row); however, when survival is scaled by distance travelled (bottom row), two patterns become apparent. First, regardless of release location or origin (Snake, Upper, or Mid-Columbia), all yearling Chinook from the Columbia River basin have remarkably similar median survival rates of 88% per 100 km of migration distance. Second, survival rates in the dammed and undammed sections of the river (the hydrosystem and LRE) are largely similar.

**Fig 2.**
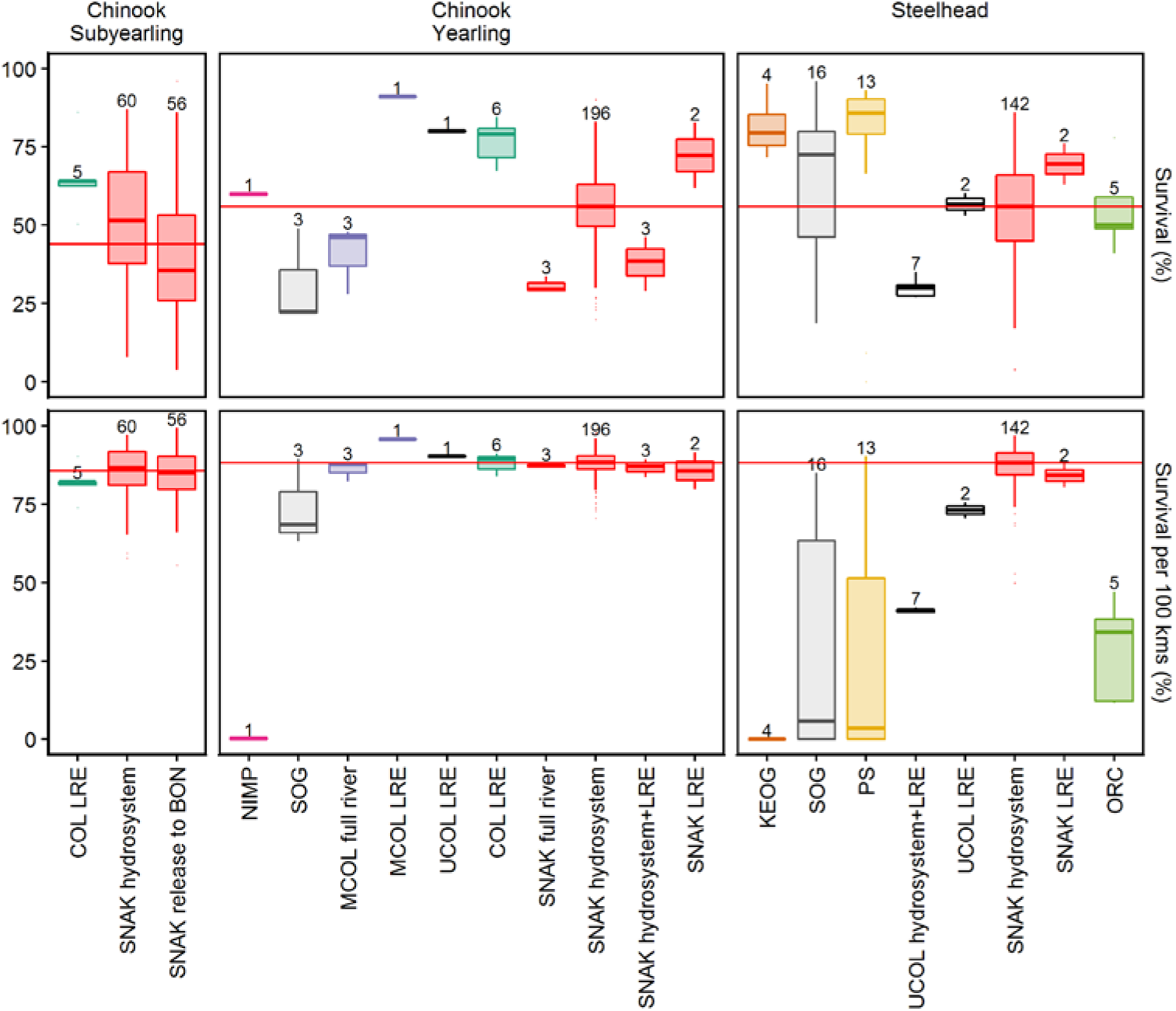
Freshwater smolt survival for west coast North American rivers. A total of N=531 annual survival estimates are included. Top row: smolt survival from release to river mouth (and intermediate locations in the case of the Columbia). Bottom row: survival per 100 km of migration distance. The red horizontal line shows the median value for all Snake River data in a given panel (red coloured bars). Data are shown as a box and whisker plot with associated sample size listed above the appropriate boxes. Abbreviations: LRE, Lower Columbia River and estuary (i.e., the river reach from just below the lowest (Bonneville) dam to the river mouth); Release to BON measures Snake River survival from hatchery release through the Snake River above Lower Granite Dam and down through the 8-dam hydrosystem to the last dam (Bonneville). Full river measures survival from release to the mouth of the Columbia River. Data sources and annual survival estimates are reported in Supplementary Table S1.

For other populations outside of the Columbia River basin which have published estimates (n=3), survival rates per 100 km varied. Survival rate of the Nimpkish River (B.C.) population was particularly low: estimated survival to the river mouth was 60% but the migration distance was only 8.5 km, resulting in only 0.25% survival per 100 km. Coldwater River (Fraser River/SOG) yearling Chinook survival rate was 63% and 68% per 100 km. Survival of hatchery-reared Chilko River Chinook (a Fraser River/SOG population) was the only population similar to the Columbia River basin; survival was 49% during their 640 km downstream migration in the Fraser River basin, resulting in a survival rate of 89% per 100 km.

A similar result is evident for Snake River steelhead which had nearly identical median survival rates per 100 km of migration distance (87%) as yearling Chinook irrespective of the section of the Columbia River basin that survival was measured over. Upper Columbia River steelhead tagged and released at Bonneville Dam in the lower Columbia River had survival rates of 70-75% per 100 km in the lower river and estuary, however, steelhead tagged and released at Rock Island Dam (UCOL) had consistently lower median survival rates, only ~41% per 100 km.

Survival rates per 100 km in the other regions for which we have steelhead data (Keogh River, Strait of Georgia, Puget Sound, and Oregon Coast) were generally lower than either the upper Columbia or Snake River. Keogh River steelhead had particularly low rates; the release site was located only 300 from the river mouth and survival ranged between 72-95%, resulting in an estimated survival rate per 100 km close to zero. Puget Sound and Oregon Coast populations had relatively short migrations to the ocean (0.3-102 km) and highly variable survival rates; these results suggest intense losses concentrated in the lowest reaches of these rivers. The only exception was hatchery-reared Skagit River steelhead which had a survival rate of 90% per 100 km.

There are no subyearling Chinook survival data available outside of the Columbia River basin, but within the basin, subyearling Chinook had similar median survival rates to yearling Chinook and steelhead in the hydrosystem and in the LRE (~85% per 100 km).

## Chinook SARs

### Coast-wide trends in adult survival (SARs)

Adult survival data for Chinook salmon are available for a varying range of years. The most extensive data sets are for the upper Columbia (both subyearling and yearling Chinook) and Snake rivers (yearlings), which extend back to the 1960s (Table S1).

Data were available for other regions beginning in the 1970s and for all regions by 2001 for yearlings, and 1987 for subyearlings.

In essentially all regions where time series extend back to the 1970s or earlier, survival to adult return has substantially decreased with time (Fig 3). A large drop in SARs for yearling Snake River Chinook is evident from the 1960s to approximately the mid-1970s, the time period when Snake River dams were completed [2,28]. Although the timing varies with region, the collapse in survival is also evident in other regions with long time series for both yearling (Upper Columbia River and—notably—Alaskan yearling stocks from SE Alaska), and subyearling Chinook (west coast Vancouver Island, the Strait of Georgia, and Puget Sound). Raymond [1, 29] (and many subsequent authors) ascribed the cause of the drop in survival to dam construction; however, declining SARs are also evident in other regions not affected by the construction of the FCRPS.

**Fig 3.**
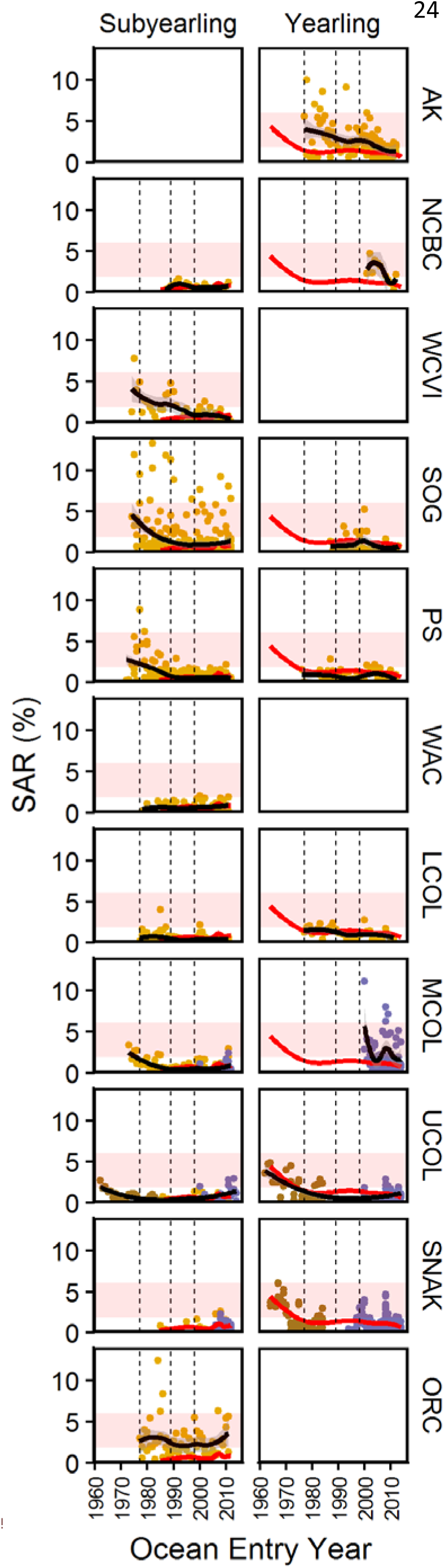
Time series of smolt to adult survival (SAR) data for west coast Chinook stocks (excluding California). Left column: subyearlings; Right column: yearlings. Regions are oriented from north (top) to south. Gold dots are SAR measurements based on CWT tags (PSC database), brown dots are SARs reported by Raymond [1], and violet dots are SARs based on PIT tags [2]. A loess curve of survival and associated 95% confidence interval (shaded region) using all available data for each panel is shown as a black line (the smoothing parameter was set to α=0.75); the loess curves for Snake River subyearling and yearling survival are overplotted in red to facilitate comparison with other regions. Blank panels indicate regions where the life history type does not occur (for example, Fall (subyearling) Chinook do not occur in Alaska, while Spring (yearling) Chinook do not occur in the low elevation streams on the west coast of Vancouver Island or Oregon coast). The major regime shift years of 1977, 1989, and 1998 are indicated by vertical lines. In this and subsequent figures the pale red band delineates the official Columbia River SAR rebuilding targets of 2-6%.

From the time of the major ocean regime shift in 1977 forwards, no substantial recovery in SARs is evident in any region. As more monitoring programs were initiated in the 1980s, SARs for all these regions were either declining or essentially fluctuating around a low mean value closely approximating the Snake River SARs (red lines) in all regions apart from the Oregon Coast; here, SARs were also roughly flat over time but at a persistently higher mean level relative to the Snake.

### Regional survival differences

When compared by region (Fig 4), median Snake River yearling (Spring) Chinook SAR (1%) is higher than the regional median SARs for Puget Sound (0.55%) and the Strait of Georgia (0.53%), and is virtually identical to median survival for the Upper (0.96%) and Lower (1.08%) Columbia River populations. Regional yearling SARs are higher than the Snake River values only for three geographic areas: the mid-Columbia River region (1.49%), Northern & Central BC (2.31%), and Alaska (1.88%). Within a few of these geographic regions, striking population-specific differences are also evident which we consider later.

**Fig 4.**
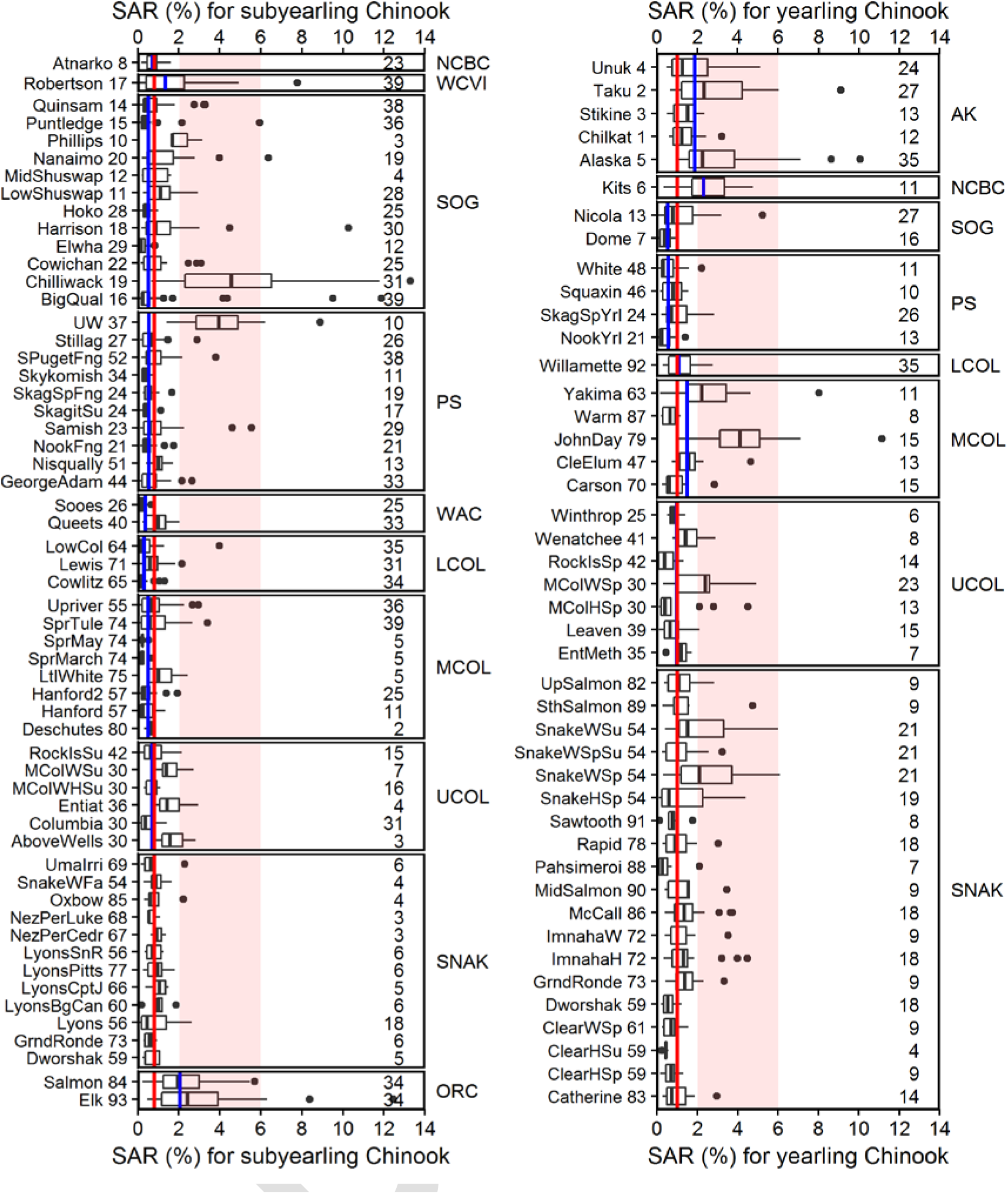
Box and whisker plot of SARs by population (all available years). The black horizontal line within each bar is the median of the SAR data available for each population. Median survival across all available data for each region is shown as a blue line; median Snake River survival for all populations combined is shown as a red line and overplotted on all panels for comparison. The number of years of data is shown to the right. To save space, abbreviated population names are used here along with the map code from Figure 1; full names for the populations are listed in Supplementary Table S2.

For subyearlings (Fall Chinook), Snake River median SARs (0.81%) are similar to or higher than median survival in other regions of the west coast apart from coastal Oregon (ORC: 2.07%) and the west coast of Vancouver Island (WCVI: 1.34%; Fig 4). As the time series plot (Fig 3) makes clear, the higher median survival evident for WCVI (Robertson Creek) Chinook relative to the Snake River may not actually be due to persistently better SARs, but rather to the longer time series of data for Robertson Creek that extends back to the period of particularly high SARs in the 1970s. Data for this time period are lacking for Snake River subyearling Chinook; we consider this issue further below.

In addition to the high median SARs for Oregon Coast and WCVI Fall Chinook, two specific subyearling hatchery populations from farther north (University of Washington Accelerated Fall Chinook in Puget Sound (3.96%), and Chilliwack Fall Chinook from the Strait of Georgia (lower Fraser River; 4.56%)) are also of note because of the strikingly large survival difference (~4X) of these stocks relative to the majority of populations within each region. The higher median SAR for yearling Chinook from the Mid-Columbia region (1.49%) is similarly due to two wild populations (Yakima: 2.21% and John Day: 4.12%) and one hatchery population (Cle Elum: 1.57%) having higher SARs while two other hatchery populations have lower SARs (Carson 0.62% and Warm Springs 0.66%) than both Snake River and Lower Columbia River median SARs (SNAK=1; LCOL=1.08%).

Strikingly, although there are some exceptional populations, no region outside the Columbia River now achieves the Columbia River basin’s official SAR recovery targets of 2%-6%. The Alaskan stocks attained these target survival levels in the early 1980s, but since that time Alaskan SARs have fallen below the Columbia River basin rebuilding targets, and in the most recent years have essentially reached the current survival rates of Columbia basin stocks (Fig 2).

## Relative survival (scaled by Snake River)

The regional-scale aggregation of SAR data provides a useful overview of survival between regions. However, important population-specific differences are potentially obscured because small numerical differences may in fact reflect large differential impacts on survival when SARs are low. For example, when regional SARs are only 1%, a population-specific SAR of 0.5% actually represents a population whose survival rate is only half that of the other populations. In addition, regional comparisons may be distorted because of trends in survival over time, and differing lengths to the various time series.

The potential influence of these factors can be reduced by normalizing the SAR estimates. In Fig 5, we divided each annual SAR estimate by the median of all Snake River SAR data available for the same year. This approach removes the potential confounding caused by temporal trends in SAR when time series with different lengths are compared. When SAR data for all available years are normalized in this way, SARs for Snake River yearling Chinook are higher than Puget Sound and Strait of Georgia and virtually indistinguishable from those for the Lower Columbia River (Willamette R) and the Upper Columbia River. Only normalized SARs for mid-Columbia, North & Central BC, and SE Alaskan populations of Spring Chinook are higher than the Snake River populations.

**Fig 5.**
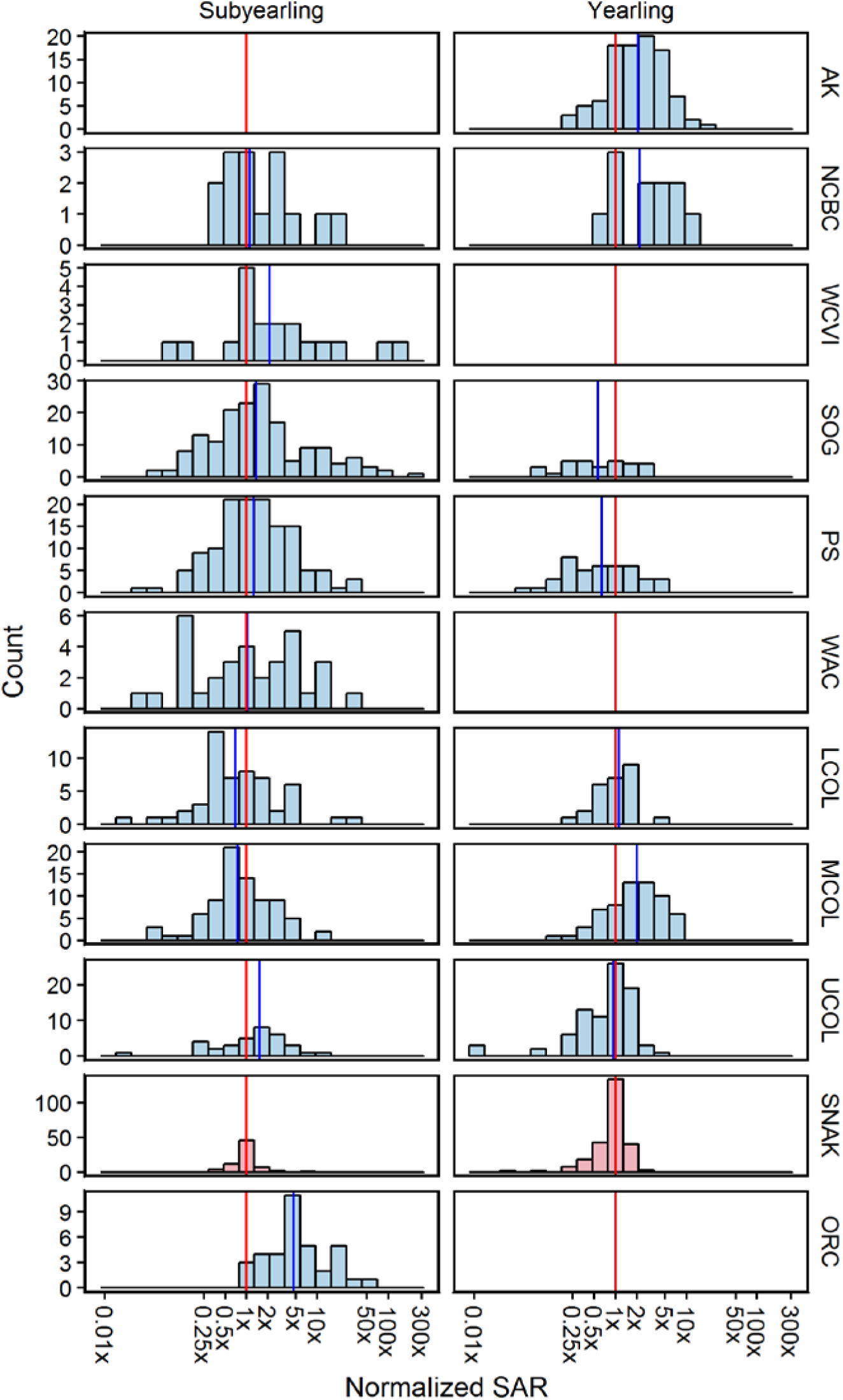
Normalized SARs calculated by dividing individual SAR estimates for each stock and each year by the median Snake River SAR for the same year. Vertical lines show the median SAR for the Snake River (red) and other regions (blue). Note the logarithmic scale on the x-axis. As in the prior plots, Columbia & Snake River SAR estimates based on PIT tags do not incorporate harvest or above-dam survival.

The situation is similar for subyearling Chinook when normalized SARs are compared: Snake River subyearling SARs are either lower (Upper Columbia; Strait of Georgia, Puget Sound), higher (Mid Columbia; Lower Columbia), or closely equivalent (Washington Coast, North-Central BC) to SARs observed for all other regions with data. The only pronounced differences are the nearly 5-fold higher survival of the two Oregon coast stocks and the roughly 2-fold higher SAR for the Robertson Creek population (west coast Vancouver Island).

## Survival by regime period

Significant changes in ocean productivity are known to impact salmon populations on time scales ranging from decades [16, 18, 32-34] to centuries [35-37]. An alternative approach to comparing survival normalized by year is to break the survival data into recognized ocean regime periods [16-18, 32, 33, 38, 39] and then compare the normalized SARs. We defined four periods based on the year of ocean entry by smolts: 1977 and earlier, 1978-89, 1990-98, and 1999 or later. The results (Fig 6) essentially mirror prior analyses, with the ratio of median Alaskan yearling Chinook survival relative to the Snake River falling from ~19X the Snake River value in the pre-1977 period to ~3X the Snake River value in the next two regime periods and then down to ~2X the Snake River value after the 1990 regime shift. Only the Alaskan, north-central BC, and Mid-Columbia populations remain ~2X higher than the Snake River populations’ SARs post-1998, but well below their earlier levels of productivity. (In fact the time series of Alaskan and north-central BC SAR data (Fig 3) show that in the most recent years SARs have fallen to Snake River SAR levels). Upper and Lower Columbia, Puget Sound, and Strait of Georgia populations all have similar or lower survival. An analogous pattern is evident for subyearling Chinook, except here it is only the Oregon Coastal populations that have persistently higher survival. The progressive collapse in survival across regimes is notable for each region.

**Fig 6.**
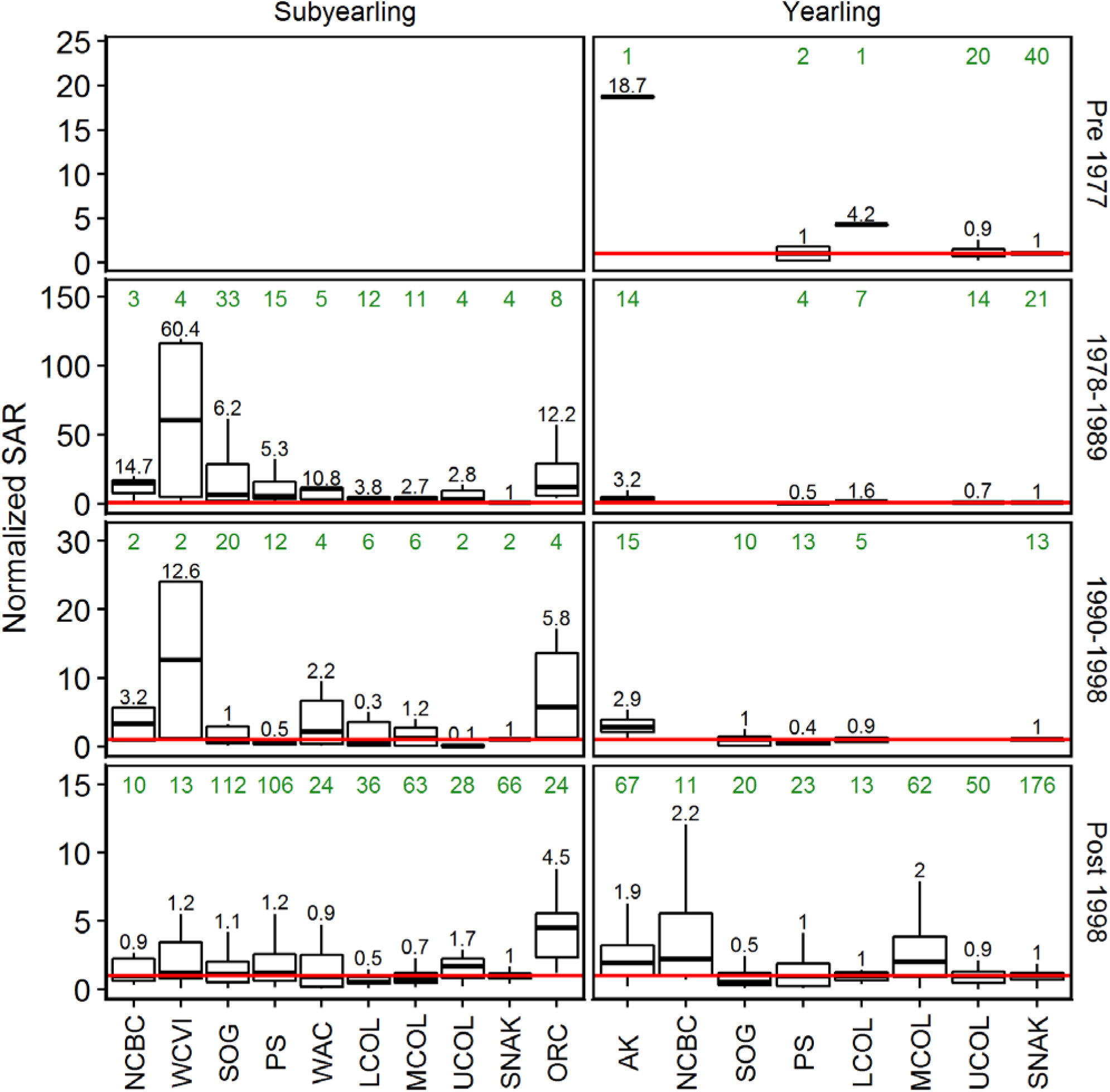
Comparison of normalized Chinook SARs by regime periods: pre-1977, 1978-1989, 1990-1998, and post 1998. Boxes and whiskers have the conventional interpretation; the horizontal red line shows the Snake R median SAR value for each regime to facilitate comparison (1.0 by definition). Sample sizes are shown above each group (green font) and the ratio of median SARs relative to the Snake River is shown immediately above the upper whiskers (black font).

## Steelhead SARs

### Coast-wide survival

Data on steelhead survival (SAR) are more geographically limited than for Chinook (Fig. 1 & Table S2), but share many of the same features (Fig 7). For simplicity, we have included the Keogh R time series from the extreme NE tip of Vancouver Island in the Strait of Georgia/Juan de Fuca Strait (SOG) region, although the population enters Queen Charlotte Strait, not the Strait of Georgia proper.

**Fig 7.**
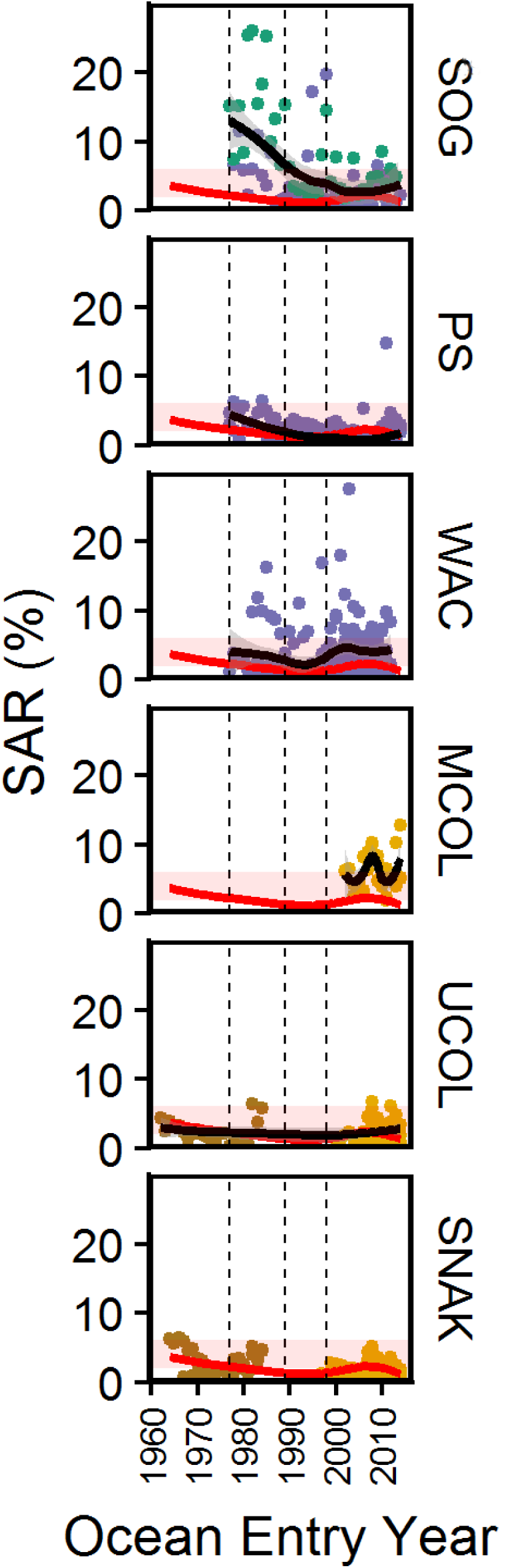
Steelhead SARS plotted against ocean entry year. Regions are oriented from north (left) to south (right); the Keogh R (KEOG) is situated on the NE tip of Vancouver Island (BC). Gold dots are SAR measurements based on PIT tags, brown dots are SARs reported by Raymond [1], and violet dots are SARs based on CWT tags. A loess curve of survival and associated 95% confidence interval (shaded region) using all available data for each panel is shown as a black line (the smoothing parameter was set to α=0.75); the Snake River loess curve is shown in red and over plotted on all other panels to facilitate comparison. The major regime shift years of 1977, 1989, and 1998 are indicated by vertical lines.

Prior to the 1977 regime shift, data are only available for the Upper Columbia and Snake Rivers (Fig 7). Similar to Snake River yearling Chinook, steelhead SARs in both the Upper Columbia and Snake Rivers declined in the period prior to the mid-1970s (when both FCRPS dam construction was completed and a major marine regime shift occurred). SAR data becomes available for Washington Coast, Puget Sound, and Strait of Georgia regions in the period after the 1977 regime shift. The very high SARs of Keogh R steelhead (northern SOG region) in the early years of the historical record, which exceeded 20% in some years, compresses the SAR differences with other regions (indicated by the LOESS curves), making the differences somewhat difficult to see.

However, plotting the data in this way demonstrate that under former climatic conditions, very high SARs were achieved in some regions.

Following ocean entry year 1990, further decline is evident in Washington Coast and Strait of Georgia steelhead SARs around ocean entry year 1990 (the time of the subsequent ocean regime shift) as well as a continuing decline in Puget Sound SARs to levels below that of Snake River steelhead. Although SAR data are not available for B.C. steelhead stocks other than the Keogh River (northern Vancouver Island), the pattern of adult returns to other southern B.C. rivers closely matches returns to Keogh River, supporting the view that the Keogh SAR pattern applies more broadly; see [40]. Both Washington outer coast (WAC) and mid-Columbia SARs are substantially higher than those the Snake River (as is Keogh), while Puget Sound SARs drop to lower values after 1990. Upper Columbia River steelhead SARs are closely similar to Snake River values.

### Regional survival differences

A few specific steelhead populations are notable for having anomalously high survival (all three mid-Columbia River and three of eight Washington Coast populations; Fig 8). Median SARs for the Snake River region (1.7%) are comparable to the Upper Columbia (1.9%) and the Washington Coast regions (2.3%), but more than double that of Puget Sound steelhead populations (0.8%). Only the median SARs for the mid-Columbia River region (5.5%) and the Strait of Georgia region (Keogh and Snow Creek; 3.3%) are appreciably higher than Snake River survival.

**Fig 8.**
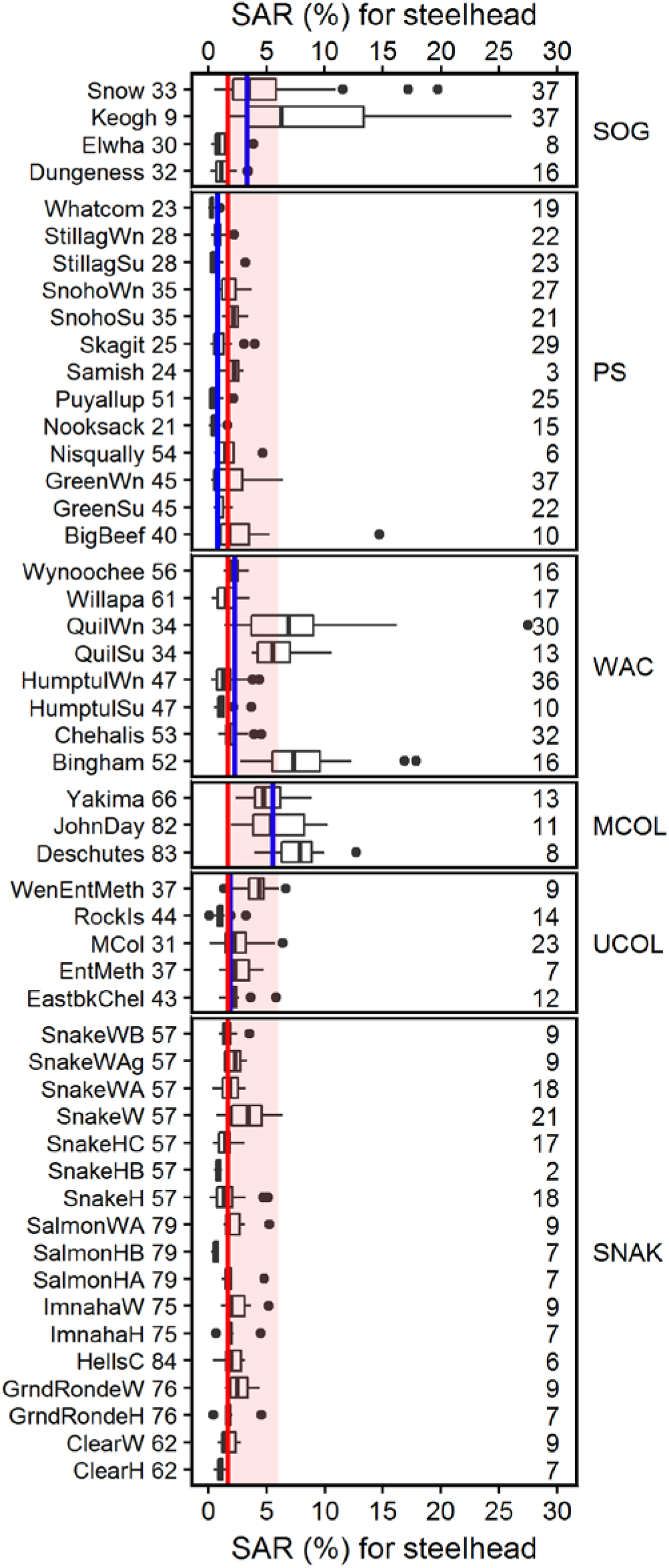
Box and whisker plot of steelhead SARs by population (all available years). Population names are listed in Supplementary Table S1. The black horizontal line within each bar is the median of the SAR data available for that population. Median survival across all available data for each geographic region is shown as a blue line; median Snake River survival for all populations combined is shown as a red line and overplotted on all panels for comparison. The number of years of data is shown to the right.

### Relative survival (scaled by Snake River)

When annual SAR estimates for individual steelhead stocks are normalized by the Snake River median SAR values in each year, a similar relationship emerges (Fig 9). Median steelhead SARs are either indistinguishable from the Snake River (Upper Columbia River), slightly higher (Washington Coast), or substantially lower (Puget Sound). Only the Mid-Columbia River and Strait of Georgia have substantively higher SARs than the Snake River when compared on a year-for-year basis.

**Fig 9.**
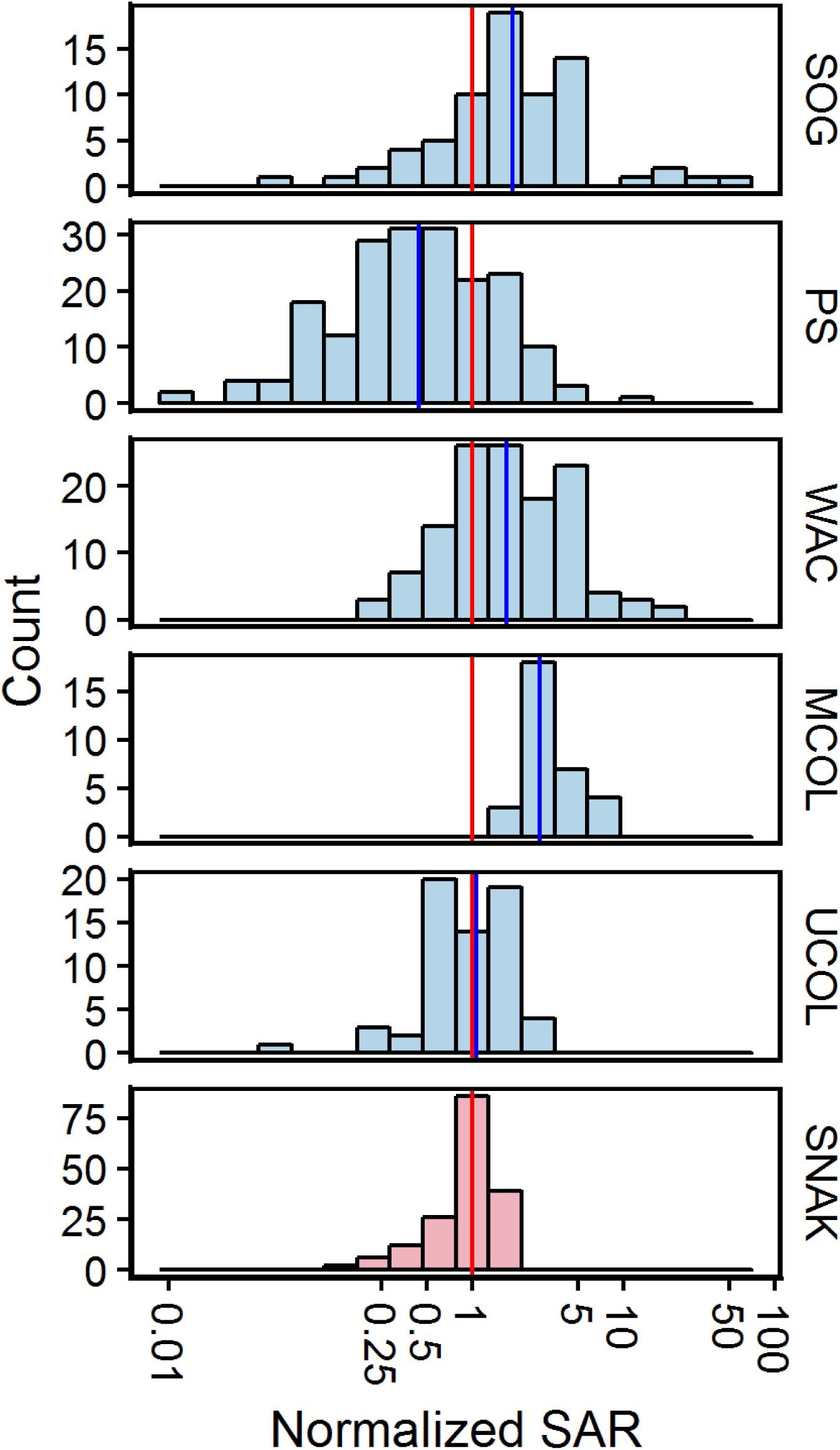
Normalized steelhead SARs obtained by dividing each individual SAR estimate (i.e., for each stock and each year) by the median SAR calculated across all available Snake River SARs for that year. The median Snake River SAR is overplotted in red. Note the logarithmic scale on the x-axis.

### Survival by regime period

This pattern becomes particularly clear when normalized steelhead SARs are examined by regime periods (Fig 10). The large drop in Strait of Georgia SARs in the post-1998 regime period is particularly notable, (in absolute terms, the drop in survival corresponds to a change in median SARs from 8.4% in the 1978-88 period to 2.6% in the post-1998 period—a three-fold decline). The second aspect to the steelhead data is the similarity of the other regions. Excluding the Mid-Columbia River, where only data for the post-1998 period are available, most other regions have median SARs roughly similar to the Snake River across all regime periods; only the mid-Columbia and SOG stand out as having consistently higher median SARs, while Puget Sound drops from higher median SARs than the Snake River to substantially lower SARs (less than half) in the post-1998 period.

**Fig 10.**
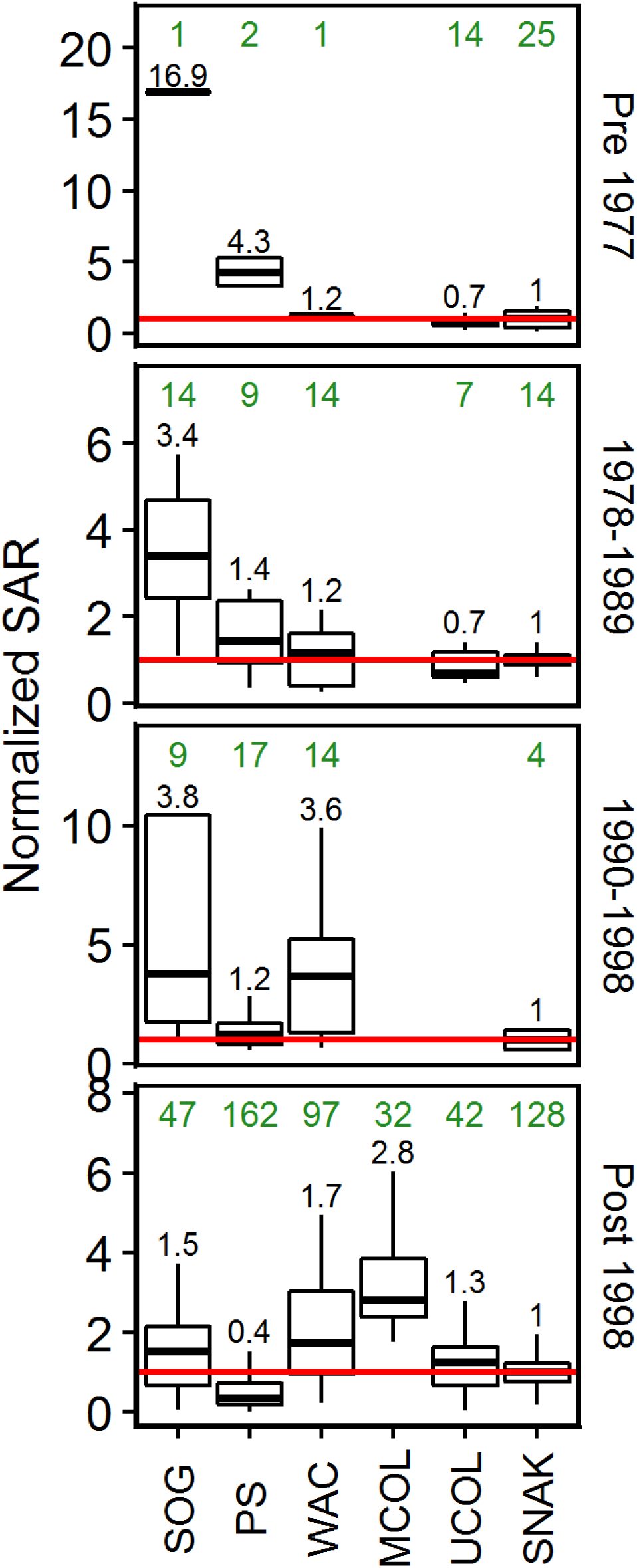
Comparison of normalized steelhead SARs by regime periods: pre-1977, 1978-1989, 1990-1998, and post 1998. Boxes and whiskers have the conventional interpretation; the horizontal red line shows the Snake R median SAR value for each regime to facilitate comparison (1.0 by definition). Sample sizes are shown above each group (green font) and the ratio of median SARs relative to the Snake River is shown immediately above the upper whiskers (black font).

## Discussion

Our analysis shows that over time Chinook and steelhead SARs have declined to reach approximately the same low level for almost all measured populations across the entire west coast of North America—with a few important exceptions that we discuss later. Although we do not have direct measurements of survival for Chinook stocks located west of SE Alaska or steelhead for regions north of Vancouver Island, the decrease in the number of adult Chinook returning to the rest of Alaska [20, 21] shows the broad region over which the conservation crisis now extends. We first address juvenile survival during seaward migration as a possible cause of the decline in adult abundance and then demonstrate the importance of the marine habitat.

### The freshwater contribution to SARs

Freshwater survival of smolts during downstream migration to the sea has been assessed for a number of river systems only over the past 15 years following the advent of miniaturized acoustic transmitters and the expansion of the PIT tag system within the Columbia River basin. The published studies collated in Table S1 report varying freshwater survival levels lying mostly within the 25-75% range for yearling Chinook. When scaled by migration distance, median survival rates of Columbia River basin yearling Chinook populations are either similar to or better than available populations from outside of the basin per 100 km of migration distance. Snake River steelhead have median survival and median survival rates very similar to Snake River yearling Chinook, and survival rates per 100 km are much better than those of all steelhead populations located outside the Snake River (Fig 2).

Within the Columbia River basin survival scaled by distance travelled is nearly constant for yearling Chinook irrespective of the source population and migration segment examined. For steelhead, downstream survival rates are lower in the Upper Columbia than the Snake River, but are still higher than values reported for outside of the Columbia River basin.

Both observations are at odds with conventional wisdom. Given the enormous focus over the past half century on improving smolt survival within the Columbia River hydrosystem, our interpretation is that these efforts were successful because survival rates are now higher than in undammed river systems. This result extends our earlier finding that Columbia River smolt survival was slightly higher than the adjacent undammed Fraser River [41], particularly for steelhead where estimates are now available for a substantial number of river systems. Thus, significant further improvement is unlikely because the Snake River now boasts the highest measured freshwater survival rates in the Pacific Northwest.

If survival rates were in fact low in the Columbia River basin, improvements in freshwater survival could potentially increase the SAR. For example, Chinook smolt survival in California ranged from 3-16% for a 516 km migration in the Sacramento River [42] to an astonishingly low 0-2% through the lower 92 km of the San Joaquin River Delta [43]. Such low survival provides substantial scope for potential improvement. This difference is important because the large drop in coast-wide SARs excluding California to around 1% and relatively high freshwater survival isolate the main conservation problem as being in the ocean.

Our results also indicate that the river mouth is a perilous location for smolts, something also noted in California [42], because survival rates scaled by distance are extremely low in rivers where post-release distance to the mouth is short, e.g., Keogh River and Big Beef Creek steelhead. Losses (presumably to predators) must be concentrated near the river mouth to result in this pattern, and continued losses from predation may well occur after ocean entry because smolts are still concentrated and the migration timing is predictable, conditions which cause predator aggregation in other situations [44, 45].

It is important to outline why past declines in freshwater survival cannot have been the driver of the observed drop in SARs—put simply, currently measured freshwater survival levels are too high. The longer SAR time series indicate at least a 4-5 fold decline over time. However, for freshwater survival to be the cause of this decline in SARs, current values of freshwater survival cannot be more than 20-25%. That they are substantially higher for many populations (Fig 2) means that it is mathematically impossible for freshwater survival to have fallen far enough to explain the decline in SARs. For example, even if downstream survival through the dams was originally 80% prior to the 1970s and then fell to 40% this would “only” produce a two-fold decline in SARs, e.g., from 6% to 3%, so the scope for primarily freshwater regulation of SARs is limited.

### The importance of marine habitat

Occam’s Razor dictates that any coherent theory to explain the large and geographically widespread drop in survival to similar low levels should be applicable to all populations. We are unable to identify a consistent mechanism of action because of the current limits to our understanding of the ocean phase, but some explanations (various forms of anthropogenic freshwater habitat disruption) are clearly less likely as explanations of poor salmon survival than others (climate-related changes in the ocean).

Salmon, as well as other anadromous fish such as lamprey and eulachon, migrate widely across a complex landscape composed of many successive freshwater and marine habitats; even something as simple as the number of distinct habitats each salmon population occupies over the duration of the marine phase is unknown. The number of returning adults is therefore successively affected by changes in survival in a complex sequence of freshwater and marine habitats, most of which are poorly understood, as the product SAR=S_1_• S_2_• S_3_•…•Sn. If survival drops to 1/10th of its original value in any one of these habitats, the SAR will also decline equivalently unless density-dependent factors occurring at some later point in the life history buffer the impact on adult returns.

Despite this, conventional conservation thinking for Pacific salmon primarily focuses on freshwater habitat issues. The rationale for this focus can be traced back to two separate events first occurring in the 1970s. The first was the passage of the U.S. Endangered Species Act (ESA) in 1973, with its strong focus on protecting and preserving habitat as the paramount priority for conservation [46]. Canada’s Species at Risk Act was enacted in 2003, and was partially modeled on the US ESA. The Canadian legislation provided a remarkably broad definition of habitat, which essentially prohibited: “damaging or destroying the residence of one or more individuals”, with residence defined as “…a dwelling-place such as a den, nest or other similar area or place, that is occupied or habitually occupied by one or more individuals during all or part of their life cycles” ([47]; p. 227). Unfortunately, “habitat” in both countries is ill-defined for migratory animals such as salmon which occupy many different habitats as they complete their life cycle. The larger question, not discussed in either country’s legislation, is this: to what degree can (or should) habitat related declines in some part of the ocean phase be compensated for by remedial action in some other part of the life history? That is, excluding the direct impacts to habitat which are obvious candidates for correction, can (and should) ocean impacts be remediated by intervening in other points in the life history?

The second event, unappreciated at the time, was a major shift in ocean climate in 1977 which had impacts on a wide range of marine fish stocks as well as salmon across the entire west coast of North America [38, 39]. The timing of this regime shift also nearly coincided with the completion of the final Snake River dam forming the Federal Columbia River Power System (FCRPS) in 1975. Not surprisingly given the understanding of salmon dynamics in that era, the ensuing decline in adult salmon returns a few years later was ascribed purely to poor smolt survival through the dams; however, as we have demonstrated, a similar drop in survival is seen in many other regions after 1977 and in purely marine species as well.

The decline in marine survival began earliest in the south and then progressively expanded farther north along the coast with time (Figs 3 & 7). Almost none of the rivers outside the Columbia have dams, so the argument that the poor performance of Snake River stocks is primarily due to the completion of the FCRPS is inconsistent with the broader data. (We are not dismissing the argument that extensive past modifications to the FCRPS have improved freshwater survival. Rather, we are making the point that further improvements in freshwater survival will have small or negligible impact on increasing adult returns and that the very large ocean impacts may in fact distort our understanding of how adult returns are related to freshwater modifications). As we will discuss, many other “single factor” reasons for poor salmon survival along the west coast also suffer from the same logical flaw that survival now seems to be poor everywhere.

### Overfishing alone can’t explain the decline

Wasser et al [48] cite this blanket statement: “Anadromous salmonids (*Oncorhynchus* sp.), which hatch in fresh water, migrate to the ocean, and then return to their natal waterways to breed, are threatened primarily by habitat loss from dams and overfishing (SOS 2011)” (Lines 98-101 of the SI). The sentiment underpinning this statement is widespread and reflects a fundamental problem with simply making a casual association between the assumed cause (freshwater habitat loss) and the effect (declining salmon stocks). We view the reality as considerably more nuanced: Fall (ocean-type) Chinook harvest levels of 50%-70% that were formerly sustainable are no longer sustainable because marine survival dropped 4-5 fold over the past few decades. The drop in marine survival is too large (75-80%) to be compensated by even the complete cessation of harvest. The magnitude of the gap is widely unappreciated, and the relatively small percentage difference between the harvest rate (50-70%) and SARs (75-80%) is misleading.

To fully compensate and maintain adult escapements, the initially sustainable harvests of the 1970s would have to be as large as the drop in marine survival has been. Algebraically,

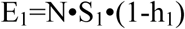

and

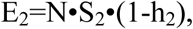

Here E_t_ is the escapement at time t, N is the number of smolts beginning migration to the sea, S_t_ is the SAR, and h_t_ is the harvest fraction, where t=1,2 is the start and end of the time series.

For escapement, E_t_, to remain constant in the two time periods implies that

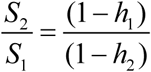

or

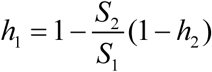

The maximum compensation management can make for declining marine survival occurs when all fisheries are curtailed completely (h_2_=0). In this case, ceasing or reducing harvest can only fully compensate if the initial rate of sustainable harvest is 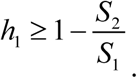. The key feature of this equation is that it is the ratio of the current to the initial period marine survival that determines how large the initial sustainable harvest rate must have been to allow full compensation by harvest rate reduction. If marine survival drops by almost an order of magnitude, as it has in at least some regions, sustainability can only be maintained if the initial sustainable harvest rate was at least 90%.

Taking the Columbia River basin as a less extreme example, marine survival has dropped from ~6% to 1%, so the initial harvest rate would have to be h_1_≥83% to allow full compensation for changing environmental conditions. Historical harvest rates reported by the PSC [49] suggest that Chinook harvest rates were on the order of 50%-60% for many subyearling stocks, implying that complete harvest rate compensation for declining marine survival would only be possible for survival ratios of 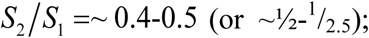; far less decrease in survival than has actually occurred.

The same major decline in survival can be seen in British Columbia after the 1977 regime shift, the period when the first real measurements of SARs for other west coast regions started. Perhaps the best measurements documenting the magnitude of the drop in British Columbia SARs was reported by Bilton et al [50]. In the early 1970s, SARs for Strait of Georgia coho of 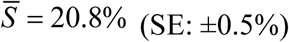 (SE: ±0.5%) and S_median_=17.2% were obtained in extensive experimental hatchery releases (six replicates of each of three size classes of smolts in each of three months (April, May, and June)). The magnitude of these survival levels (ca. one in five smolts surviving to return as adults) justified Canada’s decision to fund the Salmon Enhancement Program (SEP), a major investment in hatcheries. Yet less than two decades after the start of SEP in 1977, average coho SARs for the nearby Big Qualicum hatchery had dropped from 28.6% (1973-77 ocean entry years) to 5.6% (1990-99) and then to 1.5% (2000-2012) (data from [8, 51]). As a result, average survival rates dropped from 1 in 3.5 smolts in the 1970s to 1 in 67 smolts—a decrease to 1/20th of the initial value. (See [8] for a detailed description of the decline over time in Strait of Georgia coho SARs).

To place the magnitude of this change in perspective, by the 2000s coho SARs in the Strait of Georgia were the equivalent to surviving through a sequence of n= log(S_2000s_)/log(S_1970s_) =3.4 successive survival periods, with each time period equivalent to the entire survival process experienced by cohorts in 1973-77 (a time when intensive sport and commercial fisheries were operating, unlike recent years). Whatever the change in the environment was, it was the equivalent to the coho now remaining at sea for 60 months (five yr) instead of 1.5 yr while experiencing the overall mortality rates characteristic of the 1970s. As coho harvest rates are near zero in recent years, it is essentially all natural mortality processes that are currently operative.

Statements about the major role of particular factors in driving salmon declines (dams in the Columbia River or salmon farming in British Columbia, which developed in the 1990s) must therefore be assessed critically because salmon from other regions lacking these specific factors also return from the ocean with very poor marine survival. Thus, dams or salmon aquaculture may contribute as habitat issues to overall losses, but the essential policy debate is (1) whether modifying their operation will materially contribute to improving salmon returns, and (2) whether proposed courses of action are actually credible and cost-effective given the primary influence of ocean conditions.

### The role of dams

A wide range of west coast rivers lacking dams have similar or worse reported survival than the Snake River, both in terms of downstream smolt survival and adult return rates. We interpret this as evidence for a fundamental flaw in our biological understanding of the conservation factors controlling salmon productivity.

### Direct Mortality

Conventional thinking holds that if average marine survival was 4-6% in regions without dams, then the four-to six-fold lower survival of Columbia River Chinook populations (currently ca. 1%) would be clear evidence that the Columbia River dams were the cause of poor survival. The conclusion would then be that removing or modifying dams lying in the migration path of Snake River basin populations should increase SARs four-to six-fold, thereby achieving rebuilding targets. Yet the same conclusion, which has implicitly guided much conservation thinking, clearly cannot be used in reverse—presumably no one would argue that constructing eight dams in the Fraser River would double salmon returns, raising median Chinook survival in the years since 2000 from a mere 0.53% in the Fraser River to the Snake River’s current 1%. (Median SAR for all other Strait of Georgia yearling Chinook populations is also 0.53%; none have dams in the migration path).

A similar conclusion is evident when the level of survival through the FCRPS is assessed. Spring Chinook smolt survival through the 8-dam FCRPS ranges from 50-60% (Tables A.1 and A.2 of [5]), so even eliminating all sources of freshwater mortality during hydrosystem migration—direct impacts of the dams on survival, predation, and possible losses from disease—could only increase SARs by a factor of 0.5^-1^-0.6^-1^, or 1.7-2%. These levels are well below official rebuilding targets. Further, because a significant fraction of the downstream loss is due to predation by birds [52] and fish [53], unless all predatory wildlife species are eliminated even an increase to 1.7-2% SARs is unrealistic.

### Indirect (Delayed) Mortality

The mathematical inability of even perfect hydrosystem survival to achieve minimum rebuilding targets likely underlies the theory that delayed mortality caused by multiple dam passage contributes to poor ocean survival [5, 54-64]. Three of five Spring Chinook populations (Fig. 4) and all three steelhead populations (Fig. 8) from the mid-Columbia region not migrating through the Snake River dams have substantially higher SARs than Snake River populations, supporting this view; however, when a broader range of populations is considered the delayed mortality theory is not supported. For example, most mid-Columbia stocks of subyearling Chinook and two of five mid-Columbia yearling Chinook have similar or lower SARs relative to Snake River populations (Fig. 4). A similar pattern of anomalously high SARs is also seen for two Washington Coast steelhead populations and one (each) Strait of Georgia and Puget Sound Fall Chinook populations despite the majority of nearby populations having SARs consistent with the Snake River median (Figs. 4 & 8). Thus it is unlikely that greater dam passage causes delayed mortality in the estuary or ocean both because something unrelated to dam passage also causes a few populations to have substantially higher survival by the time the adults return from the sea in river systems lacking dams and because many populations lacking dams in the migration path now have similarly low levels of survival.

### Misplaced efforts: Case studies where the marine environment was implicated, but fresh water research was initiated

The data analyzed in this paper demonstrate both a long term coast-wide decline in survival for Chinook and steelhead and that the cause of the low SARs must predominantly be located during the marine phase of the life history. Although managers have moved to reduce Chinook harvest to partially compensate for the drop, relatively little has been done to determine the cause of the decrease in marine survival because much of the focus remains on remediating freshwater habitat.

Festinger [65] first defined the term “*cognitive dissonance*”. In brief, it can be described as the inability to recognize the true problem, despite the evidence. More formally, in psychology the term has come to mean the process by which an individual manages inconsistent thoughts, beliefs, or attitudes, especially as relating to behavioral decisions and attitude change, by modifying aspects of their cognitive process to achieve internal consistency; for example, discounting or diminishing data inconsistent with the individual’s pre-existing beliefs.

The history of west coast salmon management suggests that cognitive dissonance concerning the marine survival problem is widespread and the reason declining salmon stocks are redressed by addressing primarily freshwater habitat issues. (Interested readers should also consult Janis [66] (especially Chapter 8) for an excellent summary of the sociological factors leading to “*groupthink*” and the poor decision making processes that result). We now review three case studies to illustrate how cognitive dissonance seems to be at play in determining past operational responses to falling marine survival: (i) Rivers-Smith Inlet sockeye (Central B.C.); (ii) Columbia River Chinook and steelhead; and (iii) Upper Fraser steelhead.

### Rivers and Smith Inlet Sockeye (B.C.)

The Rivers-Smith Inlet sockeye complex formed the second largest sockeye fishery in British Columbia for much of the last century (the Fraser River being the largest). Adult harvest levels averaged around 1M sockeye for six decades (1910-1970), and escapement (measured from the late 1940s forward) was stable at ca. 400,000 adults [67]. The Rivers and Smith Inlet populations are located in adjacent watersheds in the remote central coast of BC where there is little anthropogenic impact.

Following 1970, the productivity of both the Rivers and Smith Inlet sockeye populations suddenly collapsed [67-72]. Because escapement remained stable until the 1970s [67], recruitment overfishing did not occur during this period. Probably because of the isolated location and the lack of any other nearby significant salmon fisheries, prompt management decisions to reduce harvest to near zero were promptly taken and were maintained. However, despite harvest being curtailed, the stocks did not recover as standard fisheries theory would predict, although escapements remained stable. Following the next ocean regime shift in 1989, escapement levels fell to record lows, from >1 million spawning adults to ca. 9,500 adults by 1999—a collapse to 1/100^th^ of the original stock size in just over two decades. Because the fishery had already been curtailed, no further management action was possible to compensate for the second drop in survival. There was also evidence that additional nearby sockeye stocks were impacted similarly [72]. Thus, the stock collapsed despite prompt and full action by management.

A study of the management response [67] to the collapse detailed the reasons for rejecting a freshwater cause (including using data extending back over half a century to demonstrate that pre-smolt abundance in the lake was above the long-term mean). The authors noted that *“Poor marine survival is the most parsimonious explanation for the declining fry-to-adult survival in Owikeno Lake, particularly in light of coincident declines in sockeye salmon returns per spawner at Long Lake (a nearby pristine watershed) and declines in adult sockeye salmon abundance in other populations to the north of Rivers Inlet*.”

The key findings from a joint federal and provincial government technical committee reviewing the collapse are worth quoting verbatim [68, 70]:

> *“(1) The drastic declines in abundance appear to be due to an extended period of poor marine survival that cannot be explained by any one event, such as sea-entry during an unusual El Niño year. At least two recent years (1996 and 1997) show signs of near-zero marine survival, but the reasons for those low survival rates are not known at this time*.
>
> *(2) There is little evidence to suggest that logging or other human activity in either of the drainage basins has had more than small and localized impacts on sockeye spawning and rearing. The simultaneous declines in both basins – i.e., in Owikeno, where there has been extensive logging and in Long Lake, where there has been very little – is convincing evidence that the cause of the declines does not lie in freshwater habitat disturbance”.*

The Rivers-Smith Inlet study is to our knowledge unique in North America. Not only do the twin conclusions state that the problem lies in the ocean, they also state that freshwater habitat problems were not contributive—something that is generally not possible to rule out with certainty for most salmon populations.

The joint technical committee then recommended necessary research to clarify the cause of the collapse, and regulatory action that might be taken to improve the situation. Strikingly, despite the conclusions quoted above, marine survival is not cited in any of the research which the various review committees recommended pursuing [68-70]. Instead, the committees recommended three research foci:

> *“(1) determine absolute escapement levels to Owikeno Lake… in order to improve the credibility of stock assessment;*
>
> *(2) improve the understanding of habitat use… by sockeye juveniles in Owikeno Lake and smolts in the Wannock estuary; and*
>
> *(3) investigate the status of ocean-type and lake-spawning sockeye, which are less familiar and, although not specifically covered in this plan, may require future intervention”.* (The joint committee noted that there was some evidence for an unusual sockeye life history type that went directly to sea without rearing in the lake for a year as pre-smolts (the normal life history pattern) [70]; the other committee reports have similar language).

No mention is made of addressing the marine survival issue that was at the core of the collapse; the reference to improving the understanding of smolt habitat use in the “*Wannock estuary*” mentions that *“sockeye smolts do not appear to rear in these estuaries for much time”* [69]. The report further mentions that there are numerous estuaries within River and Smith Inlets, with varying sizes and importance to salmonids. It is unclear why the Wannock was identified as particularly worthy of investigation, but the report does note that “*approximately 25% of the Wannock estuary was dyked and filled in 1973 for a log dump facility*” (i.e., almost two decades earlier).

The recommendations under Habitat are even more striking:

> *“5. Existing conceptual plans for habitat restoration developed by DFO, the provincial Watershed Restoration Program, and other stakeholders should be evaluated for their potential long term benefits to sockeye, and the feasibility of proposed restoration projects should be thoroughly assessed*.
>
> *6. Habitat restoration projects could include the reconnection of spawning and early rearing habitats along the margins of floodplains and in side-channels that have been isolated by road construction or degraded by natural and logging-related activities.*
>
> *7. Any habitat restoration projects that are undertaken should be monitored to determine their benefits for sockeye.*
>
> *8. DFO and other agencies and stakeholders should continue to collaborate on developing habitat protection strategy during resource development planning processes (e.g., CCLCRMP, Forest Development Plans).*
>
> *9. The site-specific and cumulative impacts of logging on habitats used by sockeye should be more comprehensively evaluated”. (ref. [70]; the other committee reports have similar language).*

In other words, despite the reports identifying with high certainty that freshwater habitat issues were not contributory, the committees did not attempt to understand the marine drivers and instead advocated a series of actions in freshwater.

### Columbia River

Two nearly contemporaneous studies identified the importance of either estuary (lower river) or ocean processes in controlling the poor survival of Snake River salmon. First, Kareiva *et al.* [73] applied a matrix life cycle model to demonstrate that recovery of endangered salmon populations in the Columbia River could only be achieved by improving survival in the lower river/estuary or in the coastal ocean and that (similar to our own argument) even raising main stem survival to 100% would not prevent extinction. Second, Marmorek and Peters [74] in a review of the PATH (Plan for Analyzing and Testing Hypotheses) process, stated “*Importantly, we found that the different models’ estimate of the survival rate of in-river migrants through the hydropower system, a hotly debated value, was NOT an important determinant of overall life cycle survival. Rather, the key uncertainties that emerged from these sensitivity analyses were related to the cause of mortality in the estuary and ocean*”. (See also [31]).

Probably owing to the lack of any direct information on juvenile survival in the lower Columbia River and estuary regions, two initiatives were subsequently funded: (a) the development of the JSATS acoustic telemetry system [75], and (b) directed research using commercially available telemetry equipment to formally test the delayed mortality hypothesis in the lower river and coastal ocean [76]. Both approaches established that survival was high in the lower river below Bonneville Dam and lower (but still high) in the estuary/plume region [56, 77-81]. The studies by Rechisky et al extended these results further, showing that survival was even lower in the coastal ocean region extending from the Columbia River plume to the NW tip of Vancouver Island.

Despite these findings, further work to measure ocean survival and directly address the conclusions of Kareiva *et al.* [73] and Marmorek and Peters *[74]* was not carried out. After the ocean phase was identified as being the likely cause of poor returns and not the lower river, research shifted to focus exclusively on studying freshwater survival upstream at the hydropower dams. Although several publications subsequently identified the presence of smolts in side channels within the estuary and suggested the potential importance of estuarine wetlands for salmon conservation [82-86], we are unaware of any studies that have actually identified low survival in the estuary or established the period of residency—necessary requisites for improving SARs. In summary, ocean issues remain largely unaddressed by Columbia River basin salmon managers, and it is unclear whether research soley focussing on freshwater or lower river/estuary issues will compensate for poor ocean survival.

Overall, these studies demonstrate a consistent pattern: a strong proclivity to preferentially identify and work on freshwater habitat, even in cases where marine survival has been identified as either the sole or most serious detriment to population growth.

We are not arguing that freshwater monitoring should not be conducted; monitoring population trends, and particularly survival, is critical to making informed management decisions. However, monitoring alone is insufficient. As we noted in the Introduction, the survival data used in this paper amount to a total of more than 3,000 years of sampling effort. Recent work in BC documented a substantial decline in monitoring effort in north-central BC, and the authors argued that the situation must be improved if salmon conservation efforts are to be effective [87]. While some degree of monitoring is necessary, we note that the previously substantial monitoring effort was insufficient to develop a successful management response. Obviously, if agencies cannot respond effectively to the already available data indicating a widespread collapse in marine survival of salmon populations that has been formally submitted to the PSC on an annual basis, then it is unclear why simply increasing monitoring further will lead to a more effective response. Clearly, greater monitoring alone does not necessarily lead to improved conservation outcomes.

### Managing salmon research

We are troubled that the increase in monitoring evident as survival has dwindled over time was not matched by an equally intensive analysis to assess whether existing approaches to salmon management are correct. Salmon smolt survival could only be measured in most river systems after the relatively recent development of acoustic telemetry, and PIT tags in the Columbia River Basin. Excluding smolt survival data and focusing only on the adult survival (SAR) data, the number of years of available data for Chinook and steelhead demonstrates a massive increase in monitoring over the decades (pre-1975: 117 yrs; 1976-85: 318 yrs; 1986-95: 456 yrs; 1996-2005: 715 yrs; 2006-2014: 918 yrs). Yet, despite a nearly order of magnitude increase in monitoring outputs, the point that basic aspects of this data set are in fundamental disagreement with common assumptions about the cause of the “salmon problem” has gone unrecognized. In brief, a minor industry has developed in salmon monitoring, but the implications remained unappreciated.

We view it as critical that the roles of various proposed deleterious impacts on salmon returns be rigorously quantified, rather than simply identified as important without careful thought about other potential contributing factors. As both Lackey [88, 89] and Kareiva and Marvier [90] have noted, there is a widespread implicit assumption that ecosystems unaltered by human activity are inherently good, and that restoring anthropogenically altered freshwater ecosystems will help redress the problems (e.g., [91]).

Further, competing economic activities may be unfairly blamed for the ongoing collapse of several important salmon species and unrealistic expectations placed on what various recovery options may actually achieve. This is not simply restricted to dam removal in the Columbia River basin or banning open-net salmon aquaculture in British Columba, two current hot button issues, but extends to impacts of forestry, competing rights to groundwater, or development in general. Policy options for promoting Chinook recovery need to recognize that the wide geographic footprint of poor salmon survival likely implies that efforts focused on “fixing” possible contributing factors specific to some regions are unlikely to be effective. At the very least, these efforts should be held to a significant standard: (a) clearly demonstrating a real and substantive improvement is possible, and (b) demonstrating a clear benefit relative to the proposed costs.

### Refocusing on marine migration pathways

The pattern of variation in SARs along the west coast of North America suggests that a progressive worsening of marine survival with time occurred and was accompanied by a geographic expansion northward in the region of poor survival. However, several aspects of this explanation seem to be inconsistent with the roughly similar coast-wide SARs now observed.

Fall Chinook are believed to remain shelf-resident for their entire marine phase while Spring Chinook migrate north on the shelf before eventually moving off-shelf or into the Bering Sea/Aleutian Islands. Because both groups have poor SARs, this would imply that the area of poor marine survival might be restricted to the coastal shelf off Washington, British Columbia, and SE Alaska; however, the large-scale collapse in adult Spring Chinook returns includes the Yukon and Kuskokwim Rivers (draining into the Bering Sea) and the Kenai River (Cook Inlet, Gulf of Alaska) [20-23, 92, 93]. This suggests that either the area of poor marine survival is now simultaneously large, so that exposure times to regions of poor survival are similar, or that all stocks congregate at some point in the marine phase into a more geographically confined region where their survival is similarly affected.

We have no evidence for the latter possibility. Fall (subyearling) Chinook stocks only migrate as far north as SE Alaska [94, 95] after one or more years at sea (and at least some Strait of Georgia and Puget Sound Chinook remain resident in southern BC waters for their entire marine lifespan [96-100]). The marine movements of eulachon [24] and lamprey [25], which have also undergone dramatic declines in abundance, are less well-known but are likely similar to Fall Chinook. Thus, the conditions leading to poor marine survival must be geographically widespread because western Alaska Spring Chinook are not known to migrate to the shelf region off SE Alaska or BC.

A key prediction is that stocks with the lowest SARs should have greater exposure to poor ocean conditions in southern regions. The anomalously high SARs of some specific salmon populations (Fig 4) might provide the basis for an explicit test of this prediction. Although our understanding of population-specific differences in marine migration routes is currently very limited, especially for steelhead [101, 102], there is now some developing evidence for differential salmon survival in the sea; e.g., [100, 103-105]).

Assuming that the region of poor survival progressively expanded northward along the coast at the time of successive regime shifts, there are several testable hypotheses. For example, Strait of Georgia or Puget Sound Chinook populations may have lower survival than adjacent outer coast stocks (west coast Vancouver Island, coastal Washington) because they either remain resident for a longer time period in coastal marine waters with similar survival rates (greater exposure), or because survival rates per unit time are lower in Strait of Georgia waters (greater rates of loss). This could also potentially explain why SE Alaska and north-central BC Chinook stocks in recent years still have SARs ~2X Snake River stocks and ~4X Strait of Georgia stocks (Fig 6)—Strait of Georgia Chinook stocks remain resident in the Strait of Georgia for multiple months after ocean entry [106, 107], while Snake River yearling Chinook juveniles promptly migrate north along the outer shelf to Alaska [55, 56, 108].

In this context, the consistently low survival of the Dworshak Hatchery yearling Chinook relative to other Snake River Chinook stocks is noteworthy; mean survival from Lower Granite Dam to adult return over the 2000-2015 period was only 0.58% for the Dworshak Hatchery stock versus 1.28% for McCall Hatchery and 1.29% for Imnaha Hatchery fish (ref [5], Tables B.16, B.22, & B.24). The Dworshak SAR is thus less than ½ that of the other two populations. All Snake River populations migrate through the same set of dams, so one explanation for the particularly low survival of the Dworshak population could be a differential migration to an area of the North Pacific (or Bering Sea) whose relative survival prospects was only one-half that of other regions (Columbia River Chinook salmon are known to be seasonally present in the Bering Sea and to overwinter in the Gulf of Alaska [109]). Our tenuous understanding of where Chinook and steelhead migrate to in the ocean and how long they remain in various regions (let alone how these patterns differ between populations) clearly needs urgent improvement if these issues are to be resolved.

One important possibility for establishing the geographic differences in survival is if predators increasingly target returning adult salmon. There is now ample evidence for substantial increases in marine mammal abundance and presumably predation on returning adults [110-115]. Ohlberger et al [116] reviewed the decline in size and age-structure of Chinook across western North America. They noted that consistent with the adult predation hypothesis, the decline was most pronounced in the older age groups in some (but not all) regions of the eastern Pacific. Recent work has also demonstrated that in fish, large females may confer higher fitness on their offspring [117].

Competition for food may also conceivably play a role. The geographically widespread decline in salmon growth over time seen for multiple species by the mid-1990s, and which was potentially attributed to the growth of hatchery production of pink salmon [118] has apparently continued. Continued increase in pink salmon abundance has been shown to affect plankton populations [119] and reduce survival of at least one marine seabird (shearwaters) [120, 121] as well as some salmon species [4, 122]. Thus, geographically variable rates of competition with pink salmon or marine mammal predation at older ages could both contribute to determining differential rates of salmon survival.

Large differences in SARs point to important directions for future study. A very few stocks have SARs 3 to 4-fold higher than nearby stocks. At the extreme, the Chilliwack stock of Fall (subyearling) Chinook has a median SAR of ca. 4%, an order of magnitude greater than other nearby Strait of Georgia stocks. Oregon Coastal Fall Chinook also have SARs much higher than any Columbia River basin stocks. Understanding why only a few populations consistently have high SARs when returning from the ocean as adults could pay large dividends in understanding what differences in ocean experience result in those few populations remaining productive while many others have essentially collapsed. As Peterman and Dorner [13] remarked for sockeye, “*Further research should focus on mechanisms that operate at large, multiregional spatial scales, and (or) in marine areas where numerous correlated sockeye stocks overlap*”. The markedly higher SARs evident for Oregon coastal Chinook relative to most other populations (Figs 4 & 5) may provide important guidance in this context. Riddell et al [123] (p. 580) specifically note the unique marine distributions of southern Oregon Chinook stocks, which restricts them for their entire ocean phase to life in the California Current. Nicholas and Hankin [124] (Table 2) report that Fall Chinook from the Salmon and Elk rivers in Oregon are north migrating stocks and that Oregon coastal stocks show variation in ocean migration “*with some migrating north, some south, and one stock has a mixed north and south ocean migration*” [14]. Lending credence to the possibility that ocean migration pathways influence productivity, Nehlsen et al [14] reported that the few “south migrating” Oregon Fall Chinook stocks were all characterized as having “depressed” runs in 1988 (prior to the 1989 regime shift), whereas the “north migrating” runs all had no or increasing abundance trends.

It thus seems plausible that specific salmon populations have genetically determined migration behaviours that allow them to home to distinct feeding grounds within the North Pacific, some of which confer better survival [125]. Batten et al [126] identified at least 10 geographically distinct plankton communities evident in a single transect across the North Pacific that were temporally stable across years and demonstrated that geographically distinct seabird assemblages patterned similar to the plankton communities. An analysis of tufted puffin communities [127] found that different forage fish communities also were present in different sub-regions of the Aleutian Chain. Thus geographically stable and distinct biological communities exist within the North Pacific Ocean, including the pelagic offshore. Salmon populations homing to different feeding grounds (or a succession of different feeding grounds) could therefore have very different fates if these regions develop differently over time, for which there is at least some experimental evidence [99, 128, 129].

### Columbia River basin policy implications

A critical policy question for the Columbia River basin concerns whether recovery of listed fish stocks is limited by the hydropower system as currently operated, or by ocean conditions [130]. The available evidence indicates that smolt survival during downstream freshwater migration is not higher in rivers without hydropower dams (Fig 1 and Table S1) and that a number of much shorter coastal rivers have even lower smolt survival than is experienced through the Columbia River hydrosystem, particularly when survival is scaled by distance travelled.

Bisbal and McConnaha [130] suggest several ways in which aspects of the freshwater habitat might be manipulated to improve ocean survival. However, given that recovery targets are specified in terms of attained SARs, current evidence indicates that Snake River SARs are roughly equal to (or better) than those currently achieved in the nearby Salish Sea region, a region where dams are absent. It therefore seems unlikely that recovery can be achieved without an improvement in ocean survival. Unfortunately, current scientific knowledge is simply insufficient to understand how to promote this.

### The future of Pacific salmon

Salmon are cold water fish living in a rapidly warming world. There are no easy answers for maintaining Pacific salmon populations [131] and current problems are likely to get much worse. At least eight separate ice ages are recorded in the last 800,000 years of the ice core record alone [132] and there were likely more than 50 ice ages over the past 2.6M year extent of the Quaternary [133]. Climate change is thus the norm, not the exception. However, projected levels of future climate change are far outside anything experienced in either the last 150 years of industrialization or the previous 2.5M years of the Pleistocene Epoch. Recent marine heat waves along the Pacific Coast [134] are thought to have had significant negative effects on adult salmon returns [135]. The frequency, duration, and intensity of marine heat waves are all projected to increase dramatically in future [136], further exacerbating already serious problems for salmon.

Current CO_2_ emission policies are expected to limit warming by 2100 to approximately 3.0°C [137], or more than four times greater warming than the total warming experienced over the past 150 years of the instrumental record (~0.7°C). Even if all countries meet their commitments under the Paris Agreement, these emissions scenarios are predicted to see global mean temperatures stabilize at 1.5– 2.0°C above pre-industrial levels, or ca. 2-3 times the temperature increase so far—and an increase achieved in the next 80 years, not 150 years.

Warming rates 4-6 times those experienced in the recent past mean that further surprises in how much salmon survival drops are inevitable. Given the past slow and erratic response, the likelihood that the fisheries community will identify the correct drivers of the problem and then move to successfully address them is not high if current practice continues. So far, the response has been to re-double efforts on what we know how to study (freshwater) and to largely avoid what we currently have little ability to study (the marine phase). There are real economic, social, and biological costs to doing so, with many groups now identifying various single issue factors as the primary underlying problem that needs to be fixed (hydropower dams, salmon aquaculture, forestry, land use practices, water rights). These region-specific issues cannot possibly be the drivers of the continental-scale response that we document and to further delay not only increases the threat to salmon, but to those species that rely on them, such as southern resident killer whales. Wasser et al [48] state that “*Low availability of Chinook salmon appears to be an important stressor among these fish-eating whales as well as a significant cause of late pregnancy failure, including unobserved perinatal loss… Results point to the importance of promoting Chinook salmon recovery to enhance population growth of Southern Resident killer whales.*” There are many real consequences to ineffective policy responses—lost time, inability to boost salmon abundances (for both human and non-human salmon predators alike), and elevated costs for many other industries.

The history of North American marine research on Pacific salmon has been described elsewhere [138-140]. In the last decade, small-scale efforts to describe the marine life history of juvenile salmon have developed in specific regions of the continental shelf (no small feat in itself; e.g., [141-145]). However, life history observation is useful to infer possible mechanisms affecting overall biology, not to test and validate the mechanisms driving survival. This means that the rapid learning characteristic of physics or chemistry, where hypotheses are explicitly tested and important scientific advance occurs when theories are rejected (not merely advanced), is unlikely because it is difficult to refute observation-based mechanisms. A key issue is that if marine survival problems are widespread along the Pacific Coast, mechanisms specific to only some continental shelf regions or river watersheds cannot be the major driver unless the movements of all the different salmon populations expose them to these stressors. Because poor marine survival is demonstrably widespread, research and policy predicated on the assumption that the problems are geographically specific is unlikely to be successful.

Given the massive investment in restoration and monitoring activities for Pacific salmon, the development of correct conservation analyses and policy planning is critical. Over $1 Billion is now spent annually in the continental United States alone on freshwater habitat restoration [146, 147], and there is great pressure to remove or modify hydropower dams in the Columbia River basin to rebuild salmon runs to historical levels of abundance and productivity and more recently to help endangered orca populations [48]. Within the Columbia River, the total cost of recent conservation efforts reaches or exceeds ca. 25% of FCRPS annual revenues (including foregone clean power generation), or >$0.5 Billion per year [148]. Similarly, in British Columbia, where dams are not present in the migration paths, much effort has focused on removing salmon farms to help restore Fraser River salmon populations [149-151]. Clearly, it is important to understand the impact of various anthropogenic impacts on poor salmon returns, but it is also important that the real prospects for improvement as a result of these region-specific actions are carefully assessed.

In the novel “The Sun Also Rises”, the character Bill Gorton is asked how he went bankrupt. He replied, “*Two ways. Gradually, then suddenly.*” [152]. The same process appears to be playing out in the ways fisheries science has addressed the marine survival problem for salmon. In west coast salmon management, the first issue was incorrectly diagnosing the problem (poor and worsening ocean survival) as primarily a freshwater issue, and the second problem was failing to change behaviour quickly and maintaining a focus largely on freshwater issues (which may inflict significant costs on other economic activities). As with economic bankruptcy, failing to staunch losses and persisting with previously unsuccessful behaviour is a recipe for eventual catastrophic loss. Some positive response is certainly evident, in that harvest from Chinook and steelhead fisheries was substantially restricted (e.g., [49]). However, harvest rates of shelf-resident Fall Chinook were historically in the 50%-60% range. As we have demonstrated, even the complete elimination of all harvest can only compensate for a roughly two-fold decline in marine survival; for Spring Chinook and steelhead, which are much less impacted by saltwater harvest, maximum compensation is far less.

It is not unreasonable to anticipate a further ten-fold decline in the marine survival of salmon from climate change in the relatively near future. In fact, the survival time series used in this manuscript generally end prior to 2015. The datasets therefore do not include more recent years of even worse anticipated survival. An overall pattern of low smolt to adult returns of upper Columbia and Snake river Spring Chinook salmon and steelhead has been reported for 2015-16 and is considered likely to worsen given the apparently poor early ocean survival of juvenile salmon in 2017 and unprecedented ocean conditions occurring in 2018 in the Northern California Current and Gulf of Alaska [153-155].

With the option of reducing harvest rates now almost exhausted, large reductions in escapement can now be expected similar to what occurred in the Rivers Inlet case study. Without improved understanding of what is happening at sea, potentially inappropriate policy recommendations seem likely to continue. As we have shown in the case studies, each time salmon research reached the point where it became clear that the survival problem was at sea, the ensuing response was to re-focus effort on freshwater activities, leaving the marine survival issues unaddressed while often increasing potentially costly freshwater interventions. We view this as evidence of widespread cognitive dissonance [65] and significant groupthink [66] in salmon management. A useful first step towards breaking the current impasse would be to determine whether differences in early marine migration pathways and survival of geographically close populations cause the strongly disparate SARs that we document for some populations.

## Methods

### Smolt survival data

To assess freshwater survival levels for Chinook and steelhead smolts migrating downstream, we collated all published studies for west coast North American rivers (Table S1), excluding California, and then scaled survival by distance travelled. In a previous paper [41], we collated data for the Fraser-Thompson and Columbia-Snake rivers for comparison. Our current paper includes available survival estimates from coastal Oregon to northern Vancouver Island as well as one additional smolt survival study for the Fraser River (Chilko Chinook).

Smolt survival during downstream migration was available for several regions, but the data are most extensive in the Columbia River basin where PIT tag-based studies have been conducted for over two decades and since the more recent development of acoustic tags (Juvenile Salmon Acoustic Tracking System (JSATS) or VEMCO). In other river systems, survival during downstream migration was estimated using VEMCO acoustic telemetry; there are no published PIT tags survival estimates outside the Columbia River basin. A total of 531 estimates, representing 73 individual populations, runs or release sites, and time series between 1-23 years in length were used in the comparison. All survival estimates were calculated using the Cormack-Jolly-Seber model or its derivatives. The specific methods can be found in the sources listed in Table S1.

Within the Columbia River, smolt survival was estimated from various release points including dams, traps located in rivers, and hatcheries. Downstream census points were PIT arrays located at dams or acoustic receiver arrays in the lower Columbia River or estuary. In the Columbia River basin, and particularly the Snake River, a proportion of smolts are diverted into barges at the dams and then transported downstream to below the lowest (Bonneville) dam; these fish were not included in the hydrosystem estimates, but may have been included in lower river and estuary estimates. Most Columbia River basin smolt survival estimates (N=461) were calculated by NOAA or the Fish Passage Centre using PIT tag data. Twenty-eight were from JSATS or VEMCO acoustic tag studies. Smolt survival in the basin was measured over distances ranging between 144-909 km.

In the other regions, smolt survival was estimated from hatcheries or traps, to acoustic receiver arrays near the river mouths. In some cases, fish were transported either upstream or downstream of their tagging location, e.g., Chilko Chinook were reared at a lower Fraser River hatchery but released ~500 km upstream in the Chilko River. Migration distances to the sea after release were typically much shorter than in the Columbia or Fraser Rivers (see Table S1). Excluding the Fraser and Columbia River populations, average smolt migration distance was only 19 km for all other regions.

To better compare survival across basins we scaled survival measurements by the migration distance. We used distance because travel time was not reported for all of the studies. We excluded survival estimates from [41] that were based on populations where >75% of smolts had fork lengths not meeting current best practices on acceptable tag burdens [156-161] (<130 mm for VEMCO V7 and < 140 mm for V9). This resulted in the exclusion of three survival estimates from the Nicola and Spius River tributaries of the Fraser River because of high tag burden that were included in our earlier paper [41].

## Data sources for Chinook

Most survival rates of Pacific salmon are based on mark-recapture efforts, where juveniles are “marked” or implanted with a tags--either coded wire tags (CWT) or passive integrated transponder (PIT) tags, and recaptured in the fishery or upon return to the river. The basic tag technologies are well described elsewhere [162-166].

CWT technology dates back to the 1960s. A review is provided by [167] and the application of the methodology to coastal marine migrations of coho and Chinook is described by [95, 168] and to measuring harvest and survival by [21, 49, 169]. Because the tag is implanted in the nose cartilage of smolts, the fish must be dissected to recover the tag after capture, ensuring the death of that particular tagged animal and preventing further study of its movements. CWT technology provides the basis for the Pacific Salmon Commission’s Chinook survival database used for coast-wide management of Chinook salmon under the Pacific Salmon Treaty [49]. We used this database as the source of Chinook survival data for all regions outside the Columbia River basin and for a few stocks located in the Columbia River basin (Table S2). The data are contributed by the various governments (provincial, state, and federal agencies) responsible for conducting the individual monitoring programs.

In contrast, systematic survival data based on PIT tags first came into widespread use in the Columbia River Basin in 1997 (Table S2). PIT tags are long-lived but extremely short distance radio frequency tags that can successfully transmit their unique ID code only when within <0.5 m of a detector. Although there are some recent exceptions in small rivers, the short detection range essentially limits the use of PIT tags to the Columbia River dams, which channel sufficient tagged individuals close to the detectors to generate useful results. Tagging data are contributed to a central database (PTAGIS-Pit Tag Information System) by the various agencies (state, tribal and federal) and the SARs are estimated by the Fish Passage Center. All PIT tag-based SAR estimates reported in this paper are taken from the Fish Passage Center’s Comparative Salmon Survival (CSS) Study (McCann et al [5]) and are listed in our Table S2.

Earlier survival data for Snake River Chinook populations from the 1960s and 1970s is available from Raymond [1], who noted that “*From the positive relation found between rates of return of adults and survival rates of smolts, it was apparent that mortality of smolts migrating downriver through the dam complex was the main cause of the decline in Snake River salmon and steelhead runs*”, a view that has become commonplace amongst salmon biologists. We have included these data in our analysis because Raymond’s pioneering studies [1, 29, 30] are of unique importance owing to the documentation of the high SARs occurring in the 1960s and early years of the 1970s, a time period prior to the completion of the Snake River dams and the 1977 marine regime shift, and because they defined the focus for much subsequent work in the Columbia River basin to improve survival.

The two major tagging technologies available, PIT and CWT, are largely geographically discrete, with most recent survival data from the Columbia River based on PIT tag technology and most survival data for other regions based on CWT data. Although rarely discussed, the differences in the two technologies determine what aspects of migration-phase survival are estimated. These difference are discussed below, as is a brief description of Raymond’s methods. (Raymond’s [1] early survival analysis was based on direct estimation of the number of smolts migrating downstream past Snake River dams, and dividing this value into the number of adults returning several years later; see Raymond [1] for details; as such, comments on the extent of the migration path monitored also apply to this early study).

### CWT tags

The precise technical methods of counting the number of CWT-tagged adults returning back to each population are not documented in the Pacific Salmon Commission (PSC) database by the various agencies contributing survival data; however, an example of the mark-recapture approach used by ADF&G in the Transboundary Rivers of SE Alaska and Northern British Columbia for wild Chinook stocks can be found in [21]. Most agencies operate hatcheries or (in a few cases) rotary screw traps to estimate downstream smolt numbers for wild stocks. In general, CWT-based survival estimates are calculated for hatcheries by dividing the total number of maturing adults of various ages estimated to return back to the spawning grounds or hatchery or caught in the fishery by the number of smolts released in the year of ocean entry. CWT–based survival estimates (49 time series) are only available for Chinook, not for steelhead.

The PSC database provides several measures of marine survival. In this study, we used survival data calculated as the sum of adults returning at all ages or caught in the fisheries, uninflated for losses to natural mortality for Chinook remaining at sea for longer than two years because these values are most similar to the CSS PIT-tag based survival estimates [5]. Survival estimated using this procedure slightly underestimates true survival to ocean age two because some two year old Chinook destined to mature at older ages die from natural causes prior to maturing and are therefore not enumerated. However, in cases where SAR (survival over the migratory phase of the life history), is 1%, the instantaneous total mortality rate is M_Total_=4.6. Ricker [170] suggested that the loss due to natural mortality between age two and older ages was perhaps M=0.46 yr^-1^, or only 10% of M_Total_. More recent estimates of age-specific natural mortality for Chinook are even smaller: age 2, 40%; age 3, 30%; age 4, 20%; and age 5 and older 10%; ([49], p. 8). Consequently, not correcting for natural mortality losses occurring between age 2 and older ages is unlikely to introduce major errors into the SAR estimates, particularly as the majority of Chinook return at ocean age two, and especially so in recent years [116]. (The PSC database also includes survival estimates with age 3+ adults inflated to account for losses at sea; we chose not to use these estimates because the PIT-tag based survival estimates are not inflated for mortality at older ages, so for purposes of comparison uninflated values should be more comparable. We highlight it because of our concern (see Discussion) that fisheries biologists may be underestimating the magnitude of losses at older ages and thus incorrectly assuming that the primary survival issue is in the first year after ocean entry.

### PIT tags

PIT tag estimates of SARs are taken directly from Appendix B of McCann *et al* [5], who reports SAR in several ways. We selected the SARs covering the greatest extent of the migratory life-history (i.e., smolt releases and adult returns to the highest dam available in the Columbia River basin), and we generally used SAR estimates that included jacks when available. In the mid-Columbia region, SAR estimates with jacks were sometimes available only for a shorter migration segment; in these cases we selected the SAR data sets representing the longer migration segment but excluding jacks because this was most similar to the CWT survival estimates. The extensive PIT-tag based SAR estimates for the Columbia River basin total N=45 Chinook SAR time series and N=22 steelhead SAR time series [5].

Because returning adults must ascend fish ladders with PIT tag detectors, all PIT tagged adults surviving to return can be censused (ignoring tag shedding). Dividing adult returns by the estimated number of tagged smolts reaching the most upstream dam in the year of ocean entry provides an estimate of the SAR. However, the PIT-tag based SAR estimates for the Columbia River basin differ from CWT-based estimates in three main ways: (i) they exclude losses to harvest (lowering survival relative to what is estimated in the PSC database), (ii) they exclude losses occurring from smolt release to encountering the first dam in the migration path (raising survival), and (iii) they exclude losses occurring from the time the returning adults migrate past the last dam until they reach the spawning grounds (raising survival). We review these differences in the context of the two major life history groups (yearling and subyearling) for Chinook.

### Data sources for steelhead

All steelhead data analyzed in this paper were from Kendall et al [7] updated to incorporate more recent years’ data, including new information on the actual age-structure of the adult returns. Kendall et al [7] should be consulted for a full description of these data sets and the data are available directly from Dr Kendall (Dr Neala Kendall, pers. comm.).

### Chinook

#### Division by life history type

In general, Chinook salmon display two life history types: subyearling and yearling. These life history types are identified in our analysis because there are important ecological differences between them (see reviews by [123, 171], and references therein) which likely influence survival. Subyearling smolts migrate to the ocean within a few months of hatching in the spring, while yearlings migrate to sea after completing one or more full years of life in freshwater, and are thus significantly larger at ocean entry and (generally) spend one less year in the ocean. The subyearling/yearling smolt life history types also generally correspond with adult run timing (“Fall” or “Spring”), but this linkage is somewhat subjective primarily owing to hatchery rearing practices.

Spring (yearling) populations are largely found in high altitude headwater tributaries of large river systems penetrating well into the interior of the continent such as the Columbia and Fraser rivers, and are the only Chinook life history type reported for Alaskan rivers [172, 173]. In contrast, Fall (subyearling) populations are widely found in low gradient coastal streams or in the lower mainstem of major rivers but are absent from Alaska. Early work [174] suggested an ancient genetic divide with subyearling Chinook smolts primarily produced by adult runs returning to freshwater in the fall and spawning directly after reaching their natal streams, and yearling smolts produced mainly by adults that return in the spring and then hold in freshwater without feeding until spawning in the autumn.

#### Life history & harvest rates

Spring Chinook are thought to move offshore and become purely open ocean residents for much of the marine phase, and thus essentially immune to harvest by fisheries until their return. As a consequence, offshore (pelagic) harvest of Spring Chinook is likely negligible because a convention banning high seas fishing beyond the 200 mile EEZ of Pacific Rim countries was signed in 1992 (http://www.npafc.org/new/about_convention.html) and enforcement patrols consistently find few illegal driftnet vessels and only in the far western Pacific, well beyond the known ocean distribution of North American Chinook stocks [175, 176] (but possibly not steelhead). However, some incidental harvest of immature and maturing Chinook occurs in the groundfish fisheries of the Bering Sea, with current evidence suggesting that Pacific northwest populations form ca. ⅓ of Chinook bycatch in the Bering Sea/Aleutian Islands region [109]. Unfortunately, owing to a general inability to use collected Chinook fish scales to determine the duration of the freshwater period (and thus discriminate yearling from subyearling animals), it is unclear which life history type the Pacific northwest populations analyzed in [109] represent.

In contrast, Fall Chinook are known to remain as long-term residents of the continental shelf off the west coast of North America and are thus exposed to commercial and sport harvest in coastal marine waters over multiple years [171]. Survival of shelf-resident subyearling Fall Chinook populations can therefore be significantly reduced by coastal fisheries that can harvest these animals over several years of marine life.

In reality, this relatively simple picture is more complicated. Some hatcheries hold subyearling (Fall) Chinook for an additional year before releasing them as larger yearling smolts, and others release Spring run Chinook as subyearlings (e.g., Nooksack and Skagit-See Table S2). Thus some yearling production is of smolts that presumably remain shelf-resident for several years because their intrinsic genetic make-up dictates this behaviour despite their larger (and older) age at release. Sharma and Quinn [171] also document regional differences in migration distribution between lower Columbia River and upper Columbia-Snake River Spring yearling populations which they attribute to possibly greater interbreeding between Spring and Fall run individuals in the lower Columbia River. Clarke et al [177] similarly present evidence from breeding trials that the yearling/subyearling smolting pattern follows simple Mendelian genetic rules in crosses of Fall and Spring adults (with the added twist that the sex of the parent also influences the result)! More recent work by Prince et al [178] has potentially identified a single gene in both Chinook and steelhead that controls early (spring or summer) re-entry of Chinook and steelhead that then mature in freshwater prior to spawning in the autumn; whether and how this gene might also influence marine migration behaviour is unknown.

Very recently, Riddell et al [123] have reviewed the literature and made the argument that repeated parallel evolution of the yearling and subyearling life history types in Chinook may have occurred in different watersheds. If true, this makes Healey’s [173] earlier assumption that yearling (Spring) Chinook and subyearling (Fall) Chinook have strongly dichotomous ocean migration pathways untenable unless evolution of age at ocean entry is strongly linked to migratory behaviour in the ocean.

In this paper, we have thus opted to aggregate smolt returns by age at ocean entry (yearling, subyearling) for simplicity, but note that in future it would be very valuable to disentangle the role of age at release from genetically determined differences in marine migration pathways on survival. Unfortunately, a rigorous assessment of the genetic origins of each hatchery program would almost certainly require a genetic determination of whether each hatchery program was releasing Fall or Spring Chinook, and would need to take into account whether or not hybrid populations had been created; it is a fascinating research question whose answer is completely unclear at the current time to contemplate whether the offspring of an inadvertent hybridization between a Fall and a Spring Chinook parent would rear offshore or on the shelf and how it would get there!

The difference in likely marine rearing areas is important because in CWT-based estimates of survival [49], the commercial and sport harvest of the different age groups is added to the escapement to generate the reported SAR. In contrast, PIT tag-based survival estimates for the Columbia River basin do not incorporate losses due to harvest ([5]; see p. 95). Columbia River survival estimates using PIT tags will therefore underestimate survival relative to the PSC’s CWT-based survival estimates. For example, the PSC (Table 2.7) reports average annual stock-specific harvest rates of 29-62% for Strait of Georgia Fall (subyearling) Chinook stocks with harvest rates declining over time [49]. For some Spring (yearling) Chinook, harvest rates are much lower (at the extreme, Willamette Spring Hatchery Chinook are reported as having only a 11% mean harvest rate; see Table 2.10 of [49]).

In this report we do not attempt to directly correct for the effects of harvest or differences in the proportion of the migratory phase survival is measured over because our most important conclusions seem robust to these differences, but it is important to recognize that methodological differences exist and influence survival estimates. In a few situations, we found both CWT and PIT tag-based survival estimates for the same population and the same release year (Supplementary Info S3). Relative to the 1:1 relationship expected if both methodologies “perfectly” captured the same survival process, we find ratios of 1.5SAR_CWT_:1SAR_PIT_ for three subyearling Chinook populations, consistent with expectation as CWT-based SAR estimates incorporate harvest, while PIT tag based estimates do not. Unfortunately, we did not find data to directly compare yearling Chinook survival estimates but provide some indirect comparisons in Supplementary Info S3.

Summarizing, the PIT tag-based survival estimates for the Columbia River basin are biased high relative to total migratory phase survival because these estimates exclude losses in the initial and final phases of the migration period above the dams, and biased low because they exclude harvest (which varies in potential influence between large for Fall (subyearling) and low for Spring (yearling) stocks). Finally, some of the CWT-based survival estimates for wild stocks are also biased low to some degree because they exclude survival losses occurring in the initial and final phases of the migration upstream of the enumeration points for smolts and adults. However, at least for hatchery-reared populations, smolt numbers used in the denominator of the CWT survival estimate are estimated at the time of release from the hatchery, and therefore exclude the possibility of migratory losses occurring prior to census.

For these reasons it should be noted that the strongest comparisons are within individual survival time series (the coast-wide declining trends in survival) which are based on the most consistent methodologies, while comparison between populations will be less reliable because of differences in where each populations is censused to measure survival over the migration phase. However, the coast-wide convergence of survival in recent years to very low levels at a time when most sport and commercial harvest has been drastically reduced is strong evidence that a common factor is driving the collapse in survival. It is unlikely that a single consistent conversion factor between CWT and PIT tag-based SAR estimates can be derived, because survival losses incurred upstream of the initial and final census point for calculating SARs can vary substantially between rivers and between populations within a river system and only CWT-based methods can account for losses to harvest. Only hatchery releases can potentially reach this technical standard of measuring survival over the entire migratory phase of the life history, and only if adult enumeration takes place on the spawning grounds (or at the hatchery).

### Steelhead

The migration of steelhead is poorly understood, but it is thought that they may migrate directly offshore soon after the smolts reach saltwater [102, 179]. Virtually nothing is known of their marine migration, although the open ocean distribution extends as a band bounded by specific maximum and minimum sea temperatures across the North Pacific [180]. This suggests that (similar to Spring Chinook) maturing steelhead may return directly from the offshore to their natal river and be little exposed to commercial fisheries operating in continental shelf waters except those lying on the direct migration path from the offshore. No commercial fisheries target steelhead, so harvest is limited to freshwater sport fisheries and saltwater bycatch in other fisheries.

Although many steelhead rivers and hatcheries are located in B.C., adult returns have not been accurately enumerated which prevents direct estimation of survival. As a result, SAR data for British Columbia is restricted to the Keogh River (Fig 1), where a weir located within ca. 300 m of the ocean has monitored wild steelhead since 1977 [181]. Despite the lack of SAR data for other populations, it is known that the survival trends evident for the Keogh River are mirrored in adult returns for the province of B.C. as a whole, with some differences evident between geographic regions [40, 182, 183] in more recent regime periods. Importantly, it is broadly recognized that adult steelhead returns have been falling for decades (e.g., [40, 184]) and are now at record lows; for example, the Thompson and Chilcotin tributaries of the upper Fraser River now each have adult steelhead returns of less than 200 adults [185], despite being of roughly similar size and biogeoclimatic zone to the Snake River.

For Washington State outside the Columbia River basin, steelhead SARs were assembled and analyzed for Puget Sound (Washington State), as well as a number of locations along the coasts of Juan de Fuca Strait, and the outer (western) WA coast as well as Oregon; see [7] for detailed methods. SAR data for the Columbia and Snake rivers were taken from [5]. We are unaware of additional steelhead SAR data for Alaskan rivers.

### Comparison of relative survival

Several of our analyses are based on comparisons with the SARs of Snake River populations as these are widely considered to be poor owing to the many dams (8) in the migration path, and in particular the four Snake River dams. Because the various survival time series vary in length and sampling methodology, and because survival also declines episodically with time, we chose to make the survival analysis as simple and clear cut as possible.

As a result, in each year or regime period, we divided all available individual SAR estimates by the median SAR for all Snake River populations in the same time period. The normalized median SAR for the Snake River region equals one by definition and the frequency distribution of individual normalized estimates allows us to directly judge the similarity of the SAR values between regions in the selected time periods under examination.

### Treatment of SAR data

SAR data for salmon are log-normally distributed [186]; i.e., a time series of SAR data, S_t_, will have the form S_t_ = e^*μ*+σZ_*t*_^, where *μ* and σ are respectively the mean and standard deviation of log_e_(S), and Z_t_ is the standard normal variable Z~*N*(*μ*, σ). This is important because the log-normal distribution is skewed, exhibiting occasional rare high survivals which increases the expected value above the mean. As a result, the expected value of a log-normally distributed SAR time series is neither the simple mean 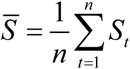 nor *μ*, but rather E(S_t_) = e^*μ*;+σ^2^/2^ (in fact, it is the median value of the log-normal distribution that is related to *μ*, as *S*_median_ = e^*μ*^). Calculating the average of the untransformed survival data, although often reported, does not have a simple statistical interpretation.

When comparing survival time series between regions, some important but subtle differences should therefore be kept in mind. We have opted to use the median (equivalent to the “geometric mean” if the data is truly log-normally distributed, 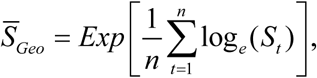 used in some literature), as well as the simple average 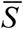 of the untransformed SAR data in a number of key comparisons. The simple average is what a number of prior studies report, and therefore what most policy makers and fisheries managers are likely comfortable interpreting. For example, the NWPPC has set a rebuilding target of 2%-6% for SARs and deemed 1% SARs (roughly the current average) to be inadequate, but did not define how SAR values should be calculated.

However, when the distribution of SARs are compared between two regions *i, j* then if the medians are found to be the similar, the implication is then that **μ**_*i*_ = **μ**_*j*_ and that the simple means of the log-transformed data are also equal; this does not, however, imply that the expected values E(S_t_) = e^*μ*+σ^2^/2^ are equal because this value also depends on the variance of the time series. For these reasons, we use both measures of central tendency

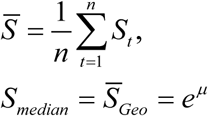

in our analysis, and not the expected mean values of the log-normal distribution E(S_t_) = e^*μ*+σ^2^/2^, owing to the more complex definition and lack of easy interpretation, which the (simple) mean and the median readily impart.

### The precision of survival estimates

The standard error on a binomial proportion reported by the CSS and PSC, survival, is
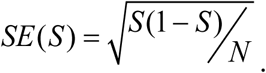. The precision of a survival estimate, φ(S), degrades as survival decreases, because

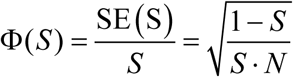

In the limit as survival approaches either 1 or zero,

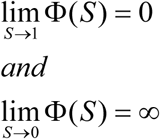

The relative uncertainty in a survival estimate with a given sample size increases without bound as survival decreases towards zero. With survival values now at 1% or less, the relative precision of a survival estimate now relative to several decades ago when survival was in the 5-6% range is

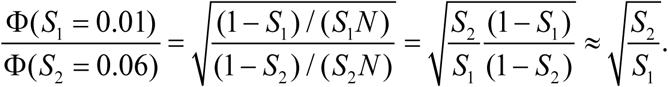

In this numeric example, where survival falls from 6% at the start of the record to 1% at the end, the uncertainty relative to the point estimate increases almost 2.5-fold 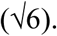. (Taking into account that both the number of outgoing smolts and the number of returning adults is not known without error, as is implicitly assumed in using the binomial probability distribution, the actual uncertainty will be even larger when these uncertainties are taken into account).

It is interesting to note that should survival fall from the current ca. 1% level to 0.1%--a ten-fold further decline—it would in fact be difficult to recognize this massive decline (a fall as large as the decline from 100% to 10% or 10% to 1% survival) because of the limited precision with which survival can be measured at such low levels. Thus for both purely mathematical reasons as well as the methodological differences between tagging approaches listed in the prior section, it is likely infeasible to obtain a perfect conversion ratio between survival estimates calculated using different methodologies (PIT vs CWT).

## Supporting information

## Acknowledgements

We particularly thank Drs Neala Kendall (WDFW) and Gayle Brown (DFO) for providing access to steelhead and Chinook SAR data, respectively, and for many discussions clarifying the interpretation of the data. The Keogh R steelhead survival project is managed by the Province of British Columbia and is currently primarily funded by the Habitat Conservation Trust Foundation (HCTF). We have also benefited from discussions with several of our colleagues, in particular Drs. XXX for early discussions on this topic. The vast (>3000 years!) of data that this paper relies upon obviously has involved many more individuals; we collectively thank them all for the effort required to generate the data used here. All data except the Keogh R steelhead SAR data are available without restriction at ??? (or from the authors).

Investigators interested in accessing Keogh SAR data should request these data from YYY. (Individuals XXX and YYY have been requested to indicate their approval to be named in the Acknowledgements, as per PLoS ONE stipulation. At time of submission, formal responses had not been received. Specific names will be incorporated during the review process).

## CRediT (Contributor Roles Taxonomy)

Conceptualization, DWW (Lead); Methodology, ADP & DWW; Software, ADP; Validation, ADP, ELR; Formal Analysis, ADP, DWW; Investigation, DWW; Data Curation, ADP; Writing – Original Draft Preparation, DWW; Writing – Review & Editing, DWW, ADP, ELR; Visualization, ADP; Supervision, DWW; Project Administration, DWW, ELR; Funding Acquisition, DWW, ELR.

## Abbreviations

BC: British Columbia, Canada
FCRPS: Federal Columbia River Power System
PSC: Pacific Salmon Commission
SAR: Smolt to Adult Return (Survival)

## References

1. Raymond HL. Effects of Hydroelectric Development and Fisheries Enhancement on Spring and Summer Chinook Salmon and Steelhead in the Columbia River Basin. North American Journal of Fisheries Management. 1988;8(1):1-24.

2. Schoen ER, Wipfli MS, Trammell EJ, Rinella DJ, Floyd AL, Grunblatt J, et al. Future of Pacific Salmon in the Face of Environmental Change: Lessons from One of the World's Remaining Productive Salmon Regions. Fisheries. 2017;42(10):538-53. doi: 10.1080/03632415.2017.1374251.

3. Irvine JR, Fukuwaka M, Kaga T, Park JH, Seong KB, Kang S, et al. Pacific Salmon Status and Abundance Trends. Vancouver, B.C.: NPAFC; 2009. p. 153 pp.

4. Ruggerone GT, Irvine JR. Numbers and Biomass of Natural- and Hatchery-Origin Pink Salmon, Chum Salmon, and Sockeye Salmon in the North Pacific Ocean, 1925–2015. Marine and Coastal Fisheries. 2018;10(2):152-68. doi: doi:10.1002/mcf2.10023.

5. McCann J, Chockley B, Cooper E, Hsu B, Schaller H, Haeseker S, et al. Comparative Survival Study of PIT-tagged Spring/Summer/Fall Chinook, Summer Steelhead, and Sockeye. 2017 Annual Report. Portland, Oregon2017.

6. Dorner B, Catalano MJ, Peterman RM. Spatial and temporal patterns of covariation in productivity of Chinook salmon populations of the Northeastern Pacific. Can J Fish Aquat Sci. 2017. doi: 10.1139/cjfas-2017-0197.

7. Kendall NW, Marston GW, Klungle MM. Declining patterns of Pacific Northwest steelhead trout (Oncorhynchus mykiss) adult abundance and smolt survival in the ocean. Can J Fish Aquat Sci. 2017;74:1275–90. doi: 10.1139/cjfas-2016-0486.

8. Zimmerman MS, Irvine JR, O’Neill M, Anderson JH, Greene CM, Weinheimer J, et al. Spatial and Temporal Patterns in Smolt Survival of Wild and Hatchery Coho Salmon in the Salish Sea. Marine and Coastal Fisheries. 2015;7(1):116-34. doi: 10.1080/19425120.2015.1012246.

9. Logerwell EA, Mantua N, Lawson PW, Francis RC, Agostini VN. Tracking environmental processes in the coastal zone for understanding and predicting Oregon coho (Oncorhynchus kisutch) marine survival. Fish Oceanogr. 2003;12:554-68.

10. Cohen BI. Commission of Inquiry into the Decline of Sockeye Salmon in the Fraser River (Canada). Final Report. Ottawa, Canada: Public Works and Government Services Canada; 2012.

11. COSEWIC. Assessment and Status Report on the Sockeye Salmon Oncorhynchus nerka, 24 Designatable Units in the Fraser River Drainage Basin, in Canada. Ottawa: Committee on the Status of Endangered Wildlife in Canada; (In Press) 2017. p. xli + 179 pp.

12. Rand PS, Goslin M, Gross MR, Irvine JR, Augerot X, McHugh PA, et al. Global Assessment of Extinction Risk to Populations of Sockeye Salmon *Oncorhynchus nerka*. PLoS ONE. 2012;7(4):e34065. doi: 10.1371/journal.pone.0034065.

13. Peterman Randall M, Dorner B. A widespread decrease in productivity of sockeye salmon (Oncorhynchus nerka) populations in western North America. Can J Fish Aquat Sci. 2012:1255-60. doi: 10.1139/f2012-063.

14. Nehlsen W, Williams JE, Lichatowich JA. Pacific Salmon at the Crossroads: Stocks at Risk from California, Oregon, Idaho, and Washington. Fisheries. 1991;16(2):4-21.

15. Allendorf FW, Bayles D, Bottom DL, Currens KP, Frissell CA, Hankin D, et al. Prioritizing Pacific Salmon Stocks for Conservation. Conservation Biology. 1997;11-1:140-52.

16. Mantua NJ, Hare SJ, Zhang Y, Wallace JM, Francis RC. A Pacific interdecadal climate oscillation with impacts on salmon production. Bulletin American Meteorological Society. 1997;78, No. 6, June:1069-79.

17. Hare SR, Mantua NJ, Francis RC. Inverse Production Regimes: Alaska and West Coast Pacific Salmon. Fisheries. 1999;24:6-14.

18. Mantua NJ, Hare SR. The Pacific Decadal Oscillation. J Oceanog. 2002;58(1):35-44. doi: 10.1023/A:1015820616384.

19. Ryding KE, Skalski JR. Multivariate regression relationships between ocean conditions and early marine survival of coho salmon (Oncorhynchus kisutch). Can J Fish Aquat Sci. 1999;56:2374-84.

20. Schindler D, Krueger C, Bisson P, Bradford M, Clark B, Conitz J, et al., editors. Arctic-Yukon-Kuskokwim Chinook Salmon Research Action Plan: Evidence of Decline of Chinook Salmon Populations and Recommendations for Future Research2013: Prepared for the AYK Sustainable Salmon Initiative (Anchorage, AK). v + 70 pp.

21. ADF&G Chinook Salmon Research Team. Chinook Salmon Stock Assessment and Research Plan, 2013. Anchorage, AK: Alaska Department of Fish & Game; 2013. Available from: http://www.adfg.alaska.gov/static/home/news/hottopics/pdfs/chinook_research_plan.pdf.

22. Cunningham CJ, Westley PAH, Adkison MD. Signals of large scale climate drivers, hatchery enhancement, and marine factors in Yukon River Chinook salmon survival revealed with a Bayesian life history model. Global Change Biology. 2018;0(0). doi: doi:10.1111/gcb.14315.

23. Bradford MJ, von Finster A, Milligan PA. Freshwater life history, habitat, and the production of Chinook Salmon from the upper Yukon basin. American Fisheries Society Symposium. 702009. p. 1-20.

24. Hay D, McCarter PB. Status of the eulachon Thaleichthys pacificus in Canada. CSAS Res Doc. 2000;2000/145:92.

25. Murauskas JG, Orlov AM, Siwicke KA. Relationships between the Abundance of Pacific Lamprey in the Columbia River and Their Common Hosts in the Marine Environment. Transactions of the American Fisheries Society. 2013;142(1):143-55.

26. Chaput G. Overview of the status of Atlantic salmon (Salmo salar) in the North Atlantic and trends in marine mortality. ICES J Mar Sci. 2012;69(9):1538-48.

27. Castonguay M, Hodson PV, Moriarty C, Drinkwater KF, Jessop BM. Is there a role of ocean environment in American and European eel decline? Fish Oceanogr. 1994;3(3):197-203. doi: 10.1111/j.1365-2419.1994.tb00097.x.

28. Dekker W, Casselman JM. The 2003 Québec Declaration of Concern About Eel Declines—11 Years Later: Are Eels Climbing Back up the Slippery Slope? Fisheries. 2014;39(12):613-4. doi: 10.1080/03632415.2014.979342.

29. Raymond HL. Effects of Dams and Impoundments on Migrations of Juvenile Chinook Salmon and Steelhead from the Snake River, 1966 to 1975. Transactions of the American Fisheries Society. 1979;108(6):505-29.

30. Raymond HL. Migration Rates of Yearling Chinook Salmon in Relation to Flows and Impoundments in the Columbia and Snake Rivers. Transactions of the American Fisheries Society. 1968;97(4):356-9. doi: 10.1577/1548-8659(1968)97[356:MROYCS]2.0.CO;2.

31. Marmorek DR, C.N. Peters and I. Parnell (eds.).. Path Final Report For Fiscal Year 1998. (Available from Bonneville Power Administration, Portland, Oregon http://www.efw.bpa.gov/Environment/PATH/reports/ISRP1999CD/PATH%20Reports/WOE_Report/). 1998:263 pp.

32. Mantua NJ. Shifting patterns in Pacific climate, West Coast salmon survival rates, and increased volatility in ecosystem services. Proceedings of the National Academy of Sciences. 2015;112(35):10823-4. doi: 10.1073/pnas.1513511112.

33. Sharma R, VÉLez-Espino LA, Wertheimer AC, Mantua N, Francis RC. Relating spatial and temporal scales of climate and ocean variability to survival of Pacific Northwest Chinook salmon (Oncorhynchus tshawytscha). Fish Oceanogr. 2013;22(1):14-31. doi: 10.1111/fog.12001.

34. Stachura MM, Mantua NJ, Scheuerell MD. Oceanographic influences on patterns in North Pacific salmon abundance. Can J Fish Aquat Sci. 2013;71(2):226-35. doi: 10.1139/cjfas-2013-0367.

35. Finney BP, Gregory-Eaves I, Sweetman J, Douglas MSV, Smol JP. Impacts of Climatic Change and Fishing on Pacific Salmon Abundance Over the Past 300 Years. Science. 2000;290(5492):795-9. doi: 10.1126/science.290.5492.795.

36. Rogers LA, Schindler DE, Lisi PJ, Holtgrieve GW, Leavitt PR, Bunting L, et al. Centennial-scale fluctuations and regional complexity characterize Pacific salmon population dynamics over the past five centuries. Proceedings of the National Academy of Sciences. 2013. doi: 10.1073/pnas.1212858110.

37. Finney BP, Gregory-Eaves I, Douglas M, Smol J. Fisheries productivity in the northeastern Pacific Ocean over the past 2,200 years. Nature. 2002;416:729-33.

38. Beamish RJ. Climate and exceptional fish production off the west coast of North America. Can J Fish Aquat Sci. 1993;50:2270-91.

39. Beamish RJ, Bouillon DR. Pacific salmon production trends in relation to climate. Can J Fish Aquat Sci. 1993;50:1002-16.

40. Welch DW, Ward BR, Smith BD, Eveson JP. Temporal and spatial responses of British Columbia steelhead (Oncorhynchus mykiss) populations to ocean climate shifts. Fish Oceanogr. 2000;9(1):17-32. doi: 10.1046/j.1365-2419.2000.00119.x

41. Welch DW, Rechisky EL, Melnychuk MC, Porter AD, Walters CJ, Clements S, et al. Survival of Migrating Salmon Smolts in Large Rivers With and Without Dams. PLoS Biology. 2008;6(10):2101-8. doi: 10.1371/journal.pbio.0060265.

42. Michel CJ, Ammann AJ, Lindley ST, Sandstrom PT, Chapman ED, Thomas MJ, et al. Chinook salmon outmigration survival in wet and dry years in California’s Sacramento River. Can J Fish Aquat Sci. 2015;72(11):1749-59. doi: 10.1139/cjfas-2014-0528.

43. Buchanan RA, Brandes PL, Skalski JR. Survival of Juvenile Fall-Run Chinook Salmon through the San Joaquin River Delta, California, 2010–2015. North American Journal of Fisheries Management. 2018 (In Press);0(0):1-17. doi: 10.1002/nafm.10063.

44. Lowe CG, Wetherbee BM, Meyer CG. Using acoustic telemetry monitoring techniques to quantify movement patterns and site fidelity of sharks and giant trevally around French Frigate Shoals and Midway Atoll. Atoll Research Bulletin. 2006;543:281–303.

45. Meyer CG, Clark TB, Papastamatiou YP, Whitney NM, Holland KN. Long-term movement patterns of tiger sharks Galeocerdo cuvier in Hawaii. Marine Ecology Progress Series. 2009;381:223-35.

46. Baur DC, Irvin WR, editors. Endangered Species Act: Law, Policy, and Perspectives. 2nd Edition (November 10, 2009) ed: American Bar Association; 2010.

47. Vanderzwaag DL, Hutchings JA. Canada's marine species at risk: science and law at the helm, but a sea of uncertainties. Ocean Development & International Law. 2005;36(3):219-59. doi: 10.1080/00908320591004333.

48. Wasser SK, Lundin JI, Ayres K, Seely E, Giles D, Balcomb K, et al. Population growth is limited by nutritional impacts on pregnancy success in endangered Southern Resident killer whales (Orcinus orca). PLOS ONE. 2017;12(6):e0179824. doi: 10.1371/journal.pone.0179824.

49. CTC (Chinook Technical Committee). 2014 Exploitation Rate Analysis and Model Calibration. Volume One. TCCHINOOK (15)-1 V. 1. Vancouver, B.C., Canada: Pacific Salmon Commission, Joint Chinook Technical Committee; 2014.

50. Bilton HT, Alderdice DF, Schnute JT. Influence of time and size at release of juvenile coho salmon (Oncorhynchus kisutch) on returns at maturity. Can J Fish Aquat Sci. 1982;39(3):426-47.

51. Irvine JR, O’Neill M, Godbout L, Schnute J. Effects of smolt release timing and size on the survival of hatchery-origin coho salmon in the Strait of Georgia. Prog Oceanogr. 2013;115(0):111-8. doi: http://dx.doi.org/10.1016/j.pocean.2013.05.014.

52. Hostetter NJ, Evans AF, Cramer BM, Collis K, Lyons DE, Roby DD. Quantifying Avian Predation on Fish Populations: Integrating Predator-Specific Deposition Probabilities in Tag Recovery Studies. Transactions of the American Fisheries Society. 2015;144(2):410-22. doi: 10.1080/00028487.2014.988882.

53. Ward DL, Petersen JH, Loch JJ. Index of Predation on Juvenile Salmonids by Northern Squawfish in the Lower and Middle Columbia River and in the Lower Snake River. Transactions of the American Fisheries Society. 1995;124(3):321-34.

54. Gosselin JL, Anderson JJ. Combining Migration History, River Conditions, and Fish Condition to Examine Cross-Life-Stage Effects on Marine Survival in Chinook Salmon. Transactions of the American Fisheries Society. 2017;146(3):408-21. doi: 10.1080/00028487.2017.1281166.

55. Rechisky EL, Welch DW, Porter AD, Hess JE, Narum SR. Testing for delayed mortality effects in the early marine life history of Columbia River yearling Chinook salmon. Mar Ecol Prog Series. 2014;496:159–80 doi: 10.3354/meps10692.

56. Rechisky EL, Welch DW, Porter AD, Jacobs-Scott MC, Winchell PM. Influence of multiple dam passage on survival of juvenile Chinook salmon in the Columbia River estuary and coastal ocean. Proc Nat Acad Sci USA. 2013. Epub Published online before print April 1, 2013. doi: 10.1073/pnas.1219910110.

57. Sandford BP, Zabel RW, Gilbreath LG, Smith SG. Exploring Latent Mortality of Juvenile Salmonids Related to Migration through the Columbia River Hydropower System. Transactions of the American Fisheries Society. 2012;141(2):343-52. doi: 10.1080/00028487.2012.664601.

58. Schaller H, Petrosky CE. Assessing Hydrosystem Influence on Delayed Mortality of Snake River Stream-Type Chinook Salmon. North American Journal of Fisheries Management. 2007;27:810–24.

59. Muir W, Marsh D, Sanford B, Smith S, Williams J. Post-Hydropower System Delayed Mortality of Transported Snake River Stream-Type Chinook Salmon: Unraveling the Mystery. Transactions of the American Fisheries Society. 2006;135:1523–34.

60. Budy P, Thiede GP, Bouwes N, Petrosky CE, Schaller H. Evidence linking delayed mortality of Snake River salmon to their earlier hydrosystem experience. North American Journal of Fisheries Management. 2002;22:35-51.

61. Zabel RW, Williams JG. Comments on “Contrasting patterns of productivity and survival rates for stream-type chinook salmon (Oncorhynchus tshawytscha) populations of the Snake and Columbia rivers” by Schaller et al (1999). Can J Fish Aquat Sci. 2000;57:1739-41.

62. Schaller HA, Petrosky CE, Langness OP. Contrasting patterns of productivity and survival rates for stream-type chinook salmon (Oncorhynchus tshawytscha) populations of the Snake and Columbia rivers. Can J Fish Aquat Sci. 1999;56(6):1031-45.

63. Schaller HA, Petrosky CE, Tinus ES. Evaluating river management during seaward migration to recover Columbia River stream-type Chinook salmon considering the variation in marine conditions. Can J Fish Aquat Sci. 2013;71(2):259-71. doi: 10.1139/cjfas-2013-0226.

64. Independent Scientific Advisory Board (ISAB). Latent Mortality Report. Review of Hypotheses and Causative Factors Contributing to Latent Mortality and their Likely Relevance to the “Below Bonneville” Component of the COMPASS Model. Portland, Oregon. : Northwest Power and Conservation Council., 2007 Document No. ISAB 2007-1.

65. Festinger L. A Theory of Cognitive Dissonance. Stanford, Calif.: Stanford University Press; 1957.

66. Janis IL. Victims of groupthink: a psychological study of foreign-policy decisions and fiascoes. Boston: Houghton Mifflin, 349 pp.; 1972.

67. McKinnell SM, Wood CC, Rutherford DT, Hyatt KD, Welch DW. The demise of Owikeno Lake sockeye salmon. North American Journal of Fisheries Management. 2001;21:774-91. doi: 10.1577/1548-8675(2001)0210774:TDOOLS2.0.CO;2.

68. Anonymous. Recommendations for a Recovery Plan for the Rivers Inlet and Smith Inlet Sockeye Salmon (Final Draft, V. 8). Fisheries & Oceans Canada. Report prepared by The Technical Coordinating Committee, Rivers and Smith Inlet Recovery Plan Working Group; 2001. p. 112 pp.

69. Rivers and Smith Inlets Salmon Ecosystem Planning Society. Rivers and Smith Inlets Salmon Ecosystem Recovery Plan. Pacific Salmon Endowment Fund Society; 2003.

70. Holtby BC. Recommendations for a Recovery Plan for the Rivers Inlet and Smith Inlet Sockeye Salmon. Prepared by: Technical Coordinating Committee Rivers Inlet and Smith Inlet Recovery Plan Working Group; 2000.

71. English KK, Glova GJ, Blakley AC. An Upstream Battle: Declines in 10 Pacific Salmon Stocks and Solutions for Their Survival. Vancouver: David Suzuki Foundation, 2008.

72. Rutherford D, Wood C. Assessment of Rivers and Smith Inlet Sockeye Salmon, with Commentary on Small Sockeye Salmon Stocks in Statistical Area 8. Canadian Stock Assessment Secretariat research document, 1480-4883 ; 2000/162; 2000. p. 57 p.

73. Kareiva PM, Marvier M, McClure MM. Recovery and management options for spring/summer Chinook salmon in the Columbia River. Science. 2000;290:977-9. doi: 10.1126/science.290.5493.977.

74. Marmorek D, Peters C. Finding a PATH toward scientific collaboration: insights from the Columbia River Basin. Conservation Ecology. 2001;5(2).

75. McMichael GA, Eppard MB, Carlson TJ, Carter JA, Ebberts BD, Brown RS, et al. The Juvenile Salmon Acoustic Telemetry System: A New Tool. Fisheries. 2010;35(1):9-22.

76. Rechisky ER, Welch DW, Porter AD, Jacobs MC, Ladouceur A. Experimental measurement of hydrosystem-induced mortality in juvenile Snake River spring Chinook salmon using a large-scale acoustic array. Can J Fish Aquat Sci. 2009;66:1019–24. doi: 10.1139/F09-078.

77. McMichael GA, Hanson AC, Harnish RA, Trott DM. Juvenile salmonid migratory behavior at the mouth of the Columbia River and within the plume. Animal Biotelemetry. 2013;1(1). doi: 10.1186/2050-3385-1-14.

78. McMichael GA, Harnish RA, Skalski JR, Deters KA, Ham KD, Townsend RL, et al. Migratory behavior and survival of juvenile salmonids in the lower Columbia River, estuary, and plume in 2010. Richland, Washington.: Pacific Northwest National Laboratory; 2011.

79. Rechisky EL, Welch DW, Porter AD, Jacobs-Scott M, Winchell PM, McKern JL. Estuarine and early-marine survival of transported and in-river migrant Snake River spring Chinook salmon smolts. Nature Scientific Reports. 2012;2(448). doi: 10.1038/srep00448.

80. Brosnan IG, Welch DW, Rechisky EL, Porter AD. Evaluating the influence of environmental factors on yearling Chinook salmon survival in the Columbia River plume (USA). Mar Ecol Prog Ser. 2014;496:181–96. doi: 10.3354/meps10550.

81. Brosnan IG, Welch DW, Scott MJ. Survival Rates of Out-Migrating Yearling Chinook Salmon in the Lower Columbia River and Plume after Exposure to Gas-Supersaturated Water. Journal of Aquatic Animal Health. 2016;28(4):240-51. doi: 10.1080/08997659.2016.1227398.

82. Johnson GE, Ploskey GR, Sather NK, Teel DJ. Residence times of juvenile salmon and steelhead in off-channel tidal freshwater habitats, Columbia River, USA. Can J Fish Aquat Sci. 2015:1-13. doi: 10.1139/cjfas-2014-0085.

83. Harnish RA, Johnson GE, McMichael GA, Hughes MS, Ebberts BD. Effect of Migration Pathway on Travel Time and Survival of Acoustic-Tagged Juvenile Salmonids in the Columbia River Estuary. Transactions of the American Fisheries Society. 2012;141(2):507-19. doi: 10.1080/00028487.2012.670576.

84. Roegner GC, McNatt R, Teel DJ, Bottom DL. Distribution, Size, and Origin of Juvenile Chinook Salmon in Shallow-Water Habitats of the Lower Columbia River and Estuary, 2002–2007. Marine and Coastal Fisheries. 2012;4(1):450-72. doi: 10.1080/19425120.2012.675982.

85. McNatt RA, Bottom DL, Hinton SA. Residency and Movement of Juvenile Chinook Salmon at Multiple Spatial Scales in a Tidal Marsh of the Columbia River Estuary. Transactions of the American Fisheries Society. 2016;145(4):774-85. doi: 10.1080/00028487.2016.1172509.

86. Sather NK, Johnson GE, Teel DJ, Storch AJ, Skalski JR, Cullinan VI. Shallow Tidal Freshwater Habitats of the Columbia River: Spatial and Temporal Variability of Fish Communities and Density, Size, and Genetic Stock Composition of Juvenile Chinook Salmon. Transactions of the American Fisheries Society. 2016;145(4):734-53. doi: 10.1080/00028487.2016.1150878.

87. Price MHH, English KK, Rosenberger AG, MacDuffee M, Reynolds JD. Canada’s Wild Salmon Policy: an assessment of conservation progress in British Columbia. Can J Fish Aquat Sci. 2017;74(10):1507-18. doi: 10.1139/cjfas-2017-0127.

88. Lackey RT. Science, scientists, and policy advocacy. Conserv Biol. 2007;21(1):12-7.

89. Lackey RT. Is Science Biased Toward Natural? Northwest Science. 2009;83(3):277-9.

90. Kareiva P, Marvier M. What Is Conservation Science? BioScience. 2012;62(11):962-9. doi: 10.1525/bio.2012.62.11.5.

91. Lichatowich J. Salmon without Rivers. A History of the Pacific Salmon Crisis. Washington, D.C.: Island Press; 1999. 317 p.

92. Lewis B, Grant WS, Brenner RE, Hamazaki T. Changes in Size and Age of Chinook Salmon Oncorhynchus tshawytscha Returning to Alaska. PLoS ONE. 2015;10(6):e0130184. doi: 10.1371/journal.pone.0130184.

93. Orsi J. The Alaska Chinook Salmon Production Enigma… What’s Going On? ONCORHYNCHUS [Internet]. 2013; XXXIII(2):[1-5 pp.]. Available from: http://www.afs-alaska.org/wp-content/uploads/Onco332.pdf.

94. Shelton AO, Satterthwaite WH, Ward EJ, Feist BE, Burke B. Using hierarchical models to estimate stock-specific and seasonal variation in ocean distribution, survivorship, and aggregate abundance of fall run Chinook. Can J Fish Aquat Sci. 2018 (In Press). doi: 10.1139/cjfas-2017-0204.

95. Weitkamp LA. Marine Distributions of Chinook Salmon from the West Coast of North America Determined by Coded Wire Tag Recoveries. Transactions of the American Fisheries Society. 2009;139(1):147-70. doi: doi:10.1577/T08-225.1.

96. Chamberlin JW, Quinn TP. Effects of natal origin on localized distributions of Chinook Salmon, Oncorhynchus tshawytscha, in the marine waters of Puget Sound, Washington. Fish Res. 2014;153:113-22.

97. O'Neill SM, West JE. Marine Distribution, Life History Traits, and the Accumulation of Polychlorinated Biphenyls in Chinook Salmon from Puget Sound, Washington. Transactions of the American Fisheries Society. 2009;138(3):616-32. doi: doi:10.1577/T08-003.1.

98. Kagley AN, Smith JM, Fresh KL, Frick KE, Quinn TP. Residency, partial migration, and late egress of subadult Chinook salmon (Oncorhynchus tshawytscha) and coho salmon (O. kisutch) in Puget Sound, Washington. Fishery Bulletin. 2017;115(4):544-56.

99. Quinn TP, Chamberlain J, Banks E. Experimental evidence of population-specific marine spatial distributions of Chinook salmon, Oncorhynchus tshawytscha.. Environmental Biology of Fishes. 2011;92(3):313-22. doi: 10.1007/s10641-011-9841-z.

100. Arostegui MC, Smith JM, Kagley AN, Spilsbury-Pucci D, Fresh KL, Quinn TP. Spatially Clustered Movement Patterns and Segregation of Subadult Chinook Salmon within the Salish Sea. Marine and Coastal Fisheries. 2017;9(1):1-12. doi: 10.1080/19425120.2016.1249580.

101. Anonymous. Insights from Genetic Analyses into Stock Specific Early Ocean Migration Behavior of Juvenile Columbia River Steelhead. Fishery Bulletin. (In Review).

102. Hartt AC, Dell MB. Early Oceanic Migrations and Growth of Juvenile Pacific Salmon and Steelhead Trout. Int North Pacific Fish Comm. 1986;46:1-105.

103. Drenner SM, Hinch SG, Furey NB, Clark TD, Li S, Ming T, et al. Transcriptome patterns and blood physiology associated with homing success of sockeye salmon during their final stage of marine migration.. Can J Fish Aquat Sci. 2017. doi: 10.1139/cjfas-2017-0391.

104. Furey NB, Vincent SP, Hinch SG, Welch DW. Variability in Migration Routes Influences Early Marine Survival of Juvenile Salmon Smolts. PLoS ONE. 2015;10(10):e0139269. doi: 10.1371/journal.pone.0139269.

105. Ruff CP, Anderson JH, Kemp IM, Kendall NW, McHugh PA, Velez-Espino A, et al. Salish Sea Chinook salmon exhibit weaker coherence in early marine survival trends than coastal populations. Fish Oceanogr. 2017;26(6):625-37. doi: 10.1111/fog.12222.

106. Welch DW, Melnychuk MC, Payne JC, Rechisky EL, Porter AD, Jackson G, et al. In situ Measurement of Coastal Ocean Movements and Survival of Juvenile Pacific Salmon. Proc Nat Acad Sci USA. 2011;108(21):8708-13 Epub May 10, 2011. doi: 10.1073/pnas.1014044108.

107. Neville CM, Beamish RJ, Chittenden CM. Poor Survival of Acoustically-Tagged Juvenile Chinook Salmon in the Strait of Georgia, British Columbia, Canada. Transactions of the American Fisheries Society. 2015;144(1):25-33. doi: 10.1080/00028487.2014.954053.

108. Fisher JP, Weitkamp LA, Teel DJ, Hinton SA, Orsi JA, Farley EV, et al. Early Ocean Dispersal Patterns of Columbia River Chinook and Coho Salmon. Transactions of the American Fisheries Society. 2014;143(1):252-72. doi: 10.1080/00028487.2013.847862.

109. Larson WA, Utter FM, Myers KW, Templin WD, Seeb JE, Guthrie Iii CM, et al. Single-nucleotide polymorphisms reveal distribution and migration of Chinook salmon (Oncorhynchus tshawytscha) in the Bering Sea and North Pacific Ocean. Can J Fish Aquat Sci. 2013;70(1):128-41. doi: 10.1139/cjfas-2012-0233.

110. Chasco BE, Kaplan IC, Thomas AC, Acevedo-Gutiérrez A, Noren DP, Ford MJ, et al. Competing tradeoffs between increasing marine mammal predation and fisheries harvest of Chinook salmon. Scientific Reports. 2017;7(1):15439. doi: 10.1038/s41598-017-14984-8.

111. Chasco B, Kaplan IC, Thomas A, Acevedo-Gutiérrez A, Noren D, Ford MJ, et al. Estimates of Chinook salmon consumption in Washington State inland waters by four marine mammal predators from 1970 to 2015. Can J Fish Aquat Sci. 2017;74(8):1173-94. doi: 10.1139/cjfas-2016-0203.

112. Thomas AC, Nelson BW, Lance MM, Deagle BE, Trites AW. Harbour seals target juvenile salmon of conservation concern. Can J Fish Aquat Sci. 2016. doi: 10.1139/cjfas-2015-0558.

113. Nelson BW, Walters CJ, Trites AW, McAllister MK. Wild Chinook salmon productivity is negatively related to seal density, and not related to hatchery releases in the Pacific Northwest. Can J Fish Aquat Sci. 2018 (In Press). doi: 10.1139/cjfas-2017-0481.

114. Keefer ML, Stansell RJ, Tackley SC, Nagy WT, Gibbons KM, Peery CA, et al. Use of Radiotelemetry and Direct Observations to Evaluate Sea Lion Predation on Adult Pacific Salmonids at Bonneville Dam. Transactions of the American Fisheries Society. 2012;141(5):1236-51. doi: 10.1080/00028487.2012.688918.

115. Wright BE, Riemer SD, Brown RF, Ougzin AM, Bucklin KA. Assessment of harbor seal predation on adult salmonids in a Pacific Northwest estuary. Ecol Applic. 2007;17(2):338-51. doi: 10.1890/05-1941.

116. Ohlberger J, Ward EJ, Schindler DE, Lewis B. Demographic changes in Chinook salmon across the Northeast Pacific Ocean. Fish and Fisheries. 2018:n/a-n/a. doi: 10.1111/faf.12272.

117. Barneche DR, Robertson DR, White CR, Marshall DJ. Fish reproductive-energy output increases disproportionately with body size. Science. 2018;360(6389):642-5. doi: 10.1126/science.aao6868.

118. Bigler BS, Welch DW, Helle JH. A review of size trends among north Pacific salmon (Oncorhynchus spp.). Can J Fish Aquat Sci. 1996;53:455-65. doi: 10.1139/f95-181.

119. Batten S, Ruggerone G, Ortiz I. Pink salmon induce a trophic cascade in plankton populations in the southern Bering Sea and around the Aleutian Islands. Fish Oceanogr. 2018;28(1):1-12. doi: 10.1111/fog.12276.

120. Springer AM, van Vliet GB, Bool N, Crowley M, Fullagar P, Lea M-A, et al. Transhemispheric ecosystem disservices of pink salmon in a Pacific Ocean macrosystem. Proceedings of the National Academy of Sciences. 2018. doi: 10.1073/pnas.1720577115.

121. Springer AM, van Vliet GB. Climate change, pink salmon, and the nexus between bottom-up and top-down forcing in the subarctic Pacific Ocean and Bering Sea. Proceedings of the National Academy of Sciences. 2014;111(18):E1880-E8. doi: 10.1073/pnas.1319089111.

122. Ruggerone GT, Connors BM. Productivity and life history of sockeye salmon in relation to competition with pink and sockeye salmon in the North Pacific Ocean. Can J Fish Aquat Sci. 2015;72(6):818-33. doi: 10.1139/cjfas-2014-0134.

123. Riddell BE, Brodeur RD, Bugaev AV, Moran P, Murphy JM, Orsi JA, et al. Chapter 5: Ocean Ecology of Chinook Salmon. In: Beamish RJ, editor. The Ocean Ecology Of Pacific Salmon And Trout. Bethesda. MD.: American Fisheries Society; 2018. p. 555-696.

124. Nicholas JW, Hankin DG. Chinook salmon populations in Oregon coastal river basins: descriptions of life histories and assessment of recent trends in run strengths. In: Oregon Dept. Fish Wildlife RDS, Information Report No. 88-1, Portland., editor. 1988.

125. Welch DW, Boehlert GW, Ward BR. POST–the Pacific Ocean Salmon Tracking Project. Oceanologica Acta. 2002;25(5):243-53.

126. Batten S, Hyrenbach D, Sydeman W, Morgan K, Henry M, Yen P, et al. Characterising Meso-Marine Ecosystems of the North Pacific. Deep Sea Research. 2006;53 (II):270–90.

127. Piatt JF, Arimitsu ML, Sydeman WJ, Thompson SA, Renner H, Zador S, et al. Biogeography of pelagic food webs in the North Pacific. Fish Oceanogr. 2018:1-18. doi: 10.1111/fog.12258.

128. Brodeur RD, Ware DM. Long-term variability in zooplankton biomass in the subarctic Pacific Ocean. Fish Oceanogr. 1992;1(1):32-8.

129. Brodeur RD, Ware DM. Interdecadal variability in distribution and catch rates of epipelagic nekton in the Northeast Pacific Ocean. Can Spec Publ Fish Aquat Sci. 1995;121:329-56.

130. Bisbal G, McConnaha W. Consideration of ocean conditions in the management of salmon. Can J Fish Aquat Sci. 1998;55(9):2178-86. doi: 10.1139/f98-108.

131. Lackey RT. Saving Wild Salmon: A 165 Year Policy Conundrum. Dubach Workshop: Science and Scientists in the Contemporary Policy Process. Portland, Oregon.: Oregon State University; 2014. p. 1-25.

132. Higgins JA, Kurbatov AV, Spaulding NE, Brook E, Introne DS, Chimiak LM, et al. Atmospheric composition 1 million years ago from blue ice in the Allan Hills, Antarctica. Proceedings of the National Academy of Sciences. 2015;112(22):6887-91. doi: 10.1073/pnas.1420232112.

133. Lewis S, Maslin M. The Human Planet: How We Created The Anthropocene. New Haven & London: Yale University Press; 2018. p. 465 pp.

134. Bond NA, Cronin MF, Freeland H, Mantua N. Causes and impacts of the 2014 warm anomaly in the NE Pacific. Geophysical Research Letters. 2015;42(9):3414-20.

135. Richerson K, Leonard J, Holland DS. Predicting the economic impacts of the 2017 West Coast salmon troll ocean fishery closure. Marine Policy. 2018 (In Press).

136. Frölicher TL, Fischer EM, Gruber N. Marine heatwaves under global warming. Nature. 2018;560(7718):360-4. doi: 10.1038/s41586-018-0383-9.

137. Sigmond M, Fyfe JC, Swart NC. Ice-free Arctic projections under the Paris Agreement. Nature Climate Change. 2018:1.

138. Pearcy W, McKinnell S. The Ocean Ecology of Salmon in the Northeast Pacific Ocean— An Abridged History. American Fisheries Society Symposium 572007. p. 7-30.

139. Brodeur RD, Myers KW, Helle JH. Research Conducted by the United States on the Early Ocean Life History of Pacific Salmon. N Pac Anadr Fish Comm Bull. 2003;3:89–131.

140. Beamish RJ, Pearsall IA, Healey MC. A history of the research on the early marine life of Pacific salmon off Canada’s Pacific coast. N Pac Anadr Fish Comm Bull. 2003;3:1-40.

141. Jacobson K, Peterson B, Trudel M, Ferguson J, Welch D, Baptista A, et al. The Marine Ecology of Columbia River Basin Salmonids: A Synthesis of Research 1998-2011. Portland, Or: Northwest Power and Conservation Council; 2012. p. 86 p + appendices.

142. Helle JH. Birth of Bering-Aleutian Salmon International Survey (BASIS). N Pac Anadr Fish Comm Bull. 2009;5:vii-viii http://www.npafc.org/new/pub_bulletin5.html.

143. SSMSP Workgroup. Research Work Plan: Marine Survival of Puget Sound Steelhead. 2014. p. 117.

144. Riddell BE, Pearsall I. Salish Sea Marine Survival Project, Canadian Backgrounder. Vancouver, B.C.: Pacific Salmon Foundation; 2015. p. 15 pp.

145. Riddell B, Pearsall I, Beamish RJ, Devlin B, Farrell AP, McFarlane S, et al. Strait of Georgia Chinook and Coho Proposal 2009. Available from: https://marinesurvivalproject.com/wp-content/uploads/PSF-PROPOSAL-Final-Full-Document.pdf.

146. Budy P, Schaller H. Evaluating tributary restoration potential for Pacific salmon recovery. ecological applications. 2007;17(4):1068–86.

147. Bernhardt ES, Palmer MA, Allan J, Alexander G, Barnas K, Brooks S, et al. Synthesizing US river restoration efforts. Science. 2005;308:636–7. doi: 10.1126/science.1109769.

148. NWPCC. 2016 Columbia River Basin Fish and Wildlife Program Costs Report. Northwest Power and Conservation Council; 2017. p. 23 p.

149. Godwin SC, Dill LM, Krkošek M, Price MHH, Reynolds JD. Reduced growth in wild juvenile sockeye salmon Oncorhynchus nerka infected with sea lice. J Fish Biol. 2017:1-17. doi: 10.1111/jfb.13325.

150. Peacock SJ, Bateman AW, Krkošek M, Connors B, Rogers S, Portner L, et al. Sea -louse parasites on juvenile wild salmon in the Broughton Archipelago, British Columbia, Canada. Ecology. 2016.

151. Morton A, Routledge R. Risk and precaution: Salmon farming. Marine Policy. 2016;74:205-12. doi: http://dx.doi.org/10.1016/j.marpol.2016.09.022.

152. Hemingway E. The Sun Also Rises: Charles Scribner & Sons; 1926. 286 pp p.

153. Wells BK, Schroeder ID, Bograd SJ, Hazen EL, Jacox MG, Leising A, et al. State of the California Current 2016–17: Still anything but normal in the north. CalCOFI Rep. 2017;58:1-55.

154. Brodeur RD, Hunsicker ME, Hann A, Miller TW. Effects of warming ocean conditions on feeding ecology of small pelagic fishes in a coastal upwelling ecosystem: a shift to gelatinous food sources. Mar Ecol Prog Ser. 2018.

155. ISAB. Review of the Comparative Survival Study (CSS) Draft 2018 Annual Report. Portland, Oregon: Northwest Power and Conservation Council; 2018. p. 27 pp.

156. Brown RS, Harnish RA, Carter KM, Boyd JW, Deters KA, Eppard MB. An Evaluation of the Maximum Tag Burden for Implantation of Acoustic Transmitters in Juvenile Chinook Salmon. North American Journal of Fisheries Management. 2010;30:499–505. doi: 10.1577/M09-038.1.

157. Collins AL, Hinch SG, Welch DW, Cooke SJ, Clark TD. Intracoelomic tagging of juvenile sockeye salmon: swimming performance, growth, survival, and post-surgical wound healing in freshwater and during a transition to seawater. Trans Amer Fish Soc. 2013;142(2):515-23. doi: 10.1080/00028487.2012.743928.

158. Smircich M, Kelly J. Extending the 2% rule: the effects of heavy internal tags on stress physiology, swimming performance, and growth in brook trout. Animal Biotelemetry. 2014;2(1):16. doi: 10.1186/2050-3385-2-16. PubMed PMID: doi:10.1186/2050-3385-2-16.

159. Morrison PR, Groot EP, Welch DW. The Effect of Short-Duration Seawater Exposure and Acoustic Tag Implantation on the Swimming Performance and Physiology of Presmolt Juvenile Coho Salmon. Transactions of the American Fisheries Society. 2013;142(3):783-92. doi: 10.1080/00028487.2013.772536.

160. Walker RW, Ashton NK, Brown RS, Liss SA, Colotelo AH, Beirão BV, et al. Effects of a novel acoustic transmitter on swimming performance and predator avoidance of juvenile Chinook Salmon: Determination of a size threshold. Fish Res. 2016;176:48-54. doi: http://dx.doi.org/10.1016/j.fishres.2015.12.007.

161. Chittenden CM, Butterworth KG, Cubitt KF, Jacobs MC, Ladouceur A, Welch DW, et al. Maximum tag to body size ratios for an endangered coho salmon (O. kisutch) stock based on physiology and performance.. Environmental Biology of Fishes:. 2008;84(1):129-40.

162. Prentice EF, Flagg TA, McCutcheon. CS. Feasibility of using implantable passive integrated transponder (PIT) tags in salmonids. American Fisheries Society Symposium. 1990a; 7:317–22.

163. Prentice EF, Flagg TA, McCutcheon CS, Brastow DF, Cross DC. Equipment, methods, and an automated data-entry station for PIT tagging. American Fisheries Society Symposium. 1990c;7:335–40.

164. Prentice EF, Flagg TA, McCutcheon CS, Brastow DF. PIT-tag monitoring systems for hydroelectric dams and fish hatcheries.. American Fisheries Society Symposium. 1990b;7:323–34.

165. Gibbons JW, Andrews KM. PIT tagging: simple technology at its best. Bioscience. 2004;54(5):447-54.

166. Jefferts KB, Bergman PK, Fiscus HF. A coded wire identification system for macro-organisms. Nature. 1963;198:160-2.

167. Johnson JK, editor Regional overview of coded wire tagging of anadromous salmon and steelhead in northwest America. American Fisheries Society Symposium; 1990.

168. Weitkamp L, Neely K. Coho salmon (Oncorhynchus kisutch) ocean migration patterns: insight from marine coded-wire tag recoveries. Can J Fish Aquat Sci. 2002;59:1100-115.

169. Bernard DR, Clark JE. Estimating salmon harvest with coded-wire tags. Can J Fish Aquat Sci. 1996;53(10):2323-32.

170. Ricker WE. Ocean growth and mortality of pink and chum salmon. J Fish Res Bd Canada. 1964;21(5):905-6. doi: 10.1139/f64-087.

171. Sharma R, Quinn TP. Linkages between life history type and migration pathways in freshwater and marine environments for Chinook salmon, Oncorhynchus tshawytscha. Acta Oecologica. 2012;41:1-13. doi: http://dx.doi.org/10.1016/j.actao.2012.03.002.

172. Healey MC. Life history of Chinook salmon (Oncorhynchus tshawytscha). In: Groot C, Margolis L, editors. Pacific Salmon Life Histories. Vancouver: University of British Columbia Press; 1991. p. 311-94.

173. Healey MC. Coastwide Distribution and Ocean Migration Patterns of Stream-and Ocean-Type Chinook Salmon, Oncorhynchus tshawytscha. Canadian Field Naturalist. 1983;97(4):427-33.

174. Waples RS, Teel DJ, Myers JM, Marshall AR. Life-history divergence in Chinook salmon: historic contingency and parallel evolution. Evolution. 2004;58(2):386-403. doi: 10.1554/03-323.

175. Myers KW, Rogers DE. Stock origins of chinook slamon in incidental catches by groundfish fisheries in the eastern Bering Sea. North American Journal of Fisheries Management. 1988;8:162-71.

176. Myers KW, Haris CK, Knudsen CM, Walker RV, Davis ND, Rogers DE. Stock origins of chinook salmon in the area of the Japanese mothership salmon fishery. North American Journal of Fisheries Management. 1987;7 No. 4 (Fall 1987):459-74.

177. Clarke WC, Withler RE, Shelbourn JE. Inheritance of smolting phenotypes in backcrosses of hybrid stream-type x ocean-type chinook salmon (Oncorhynchus tshawytscha). Estuaries. 1994;17:13-25.

178. Prince DJ, O’Rourke SM, Thompson TQ, Ali OA, Lyman HS, Saglam IK, et al. The evolutionary basis of premature migration in Pacific salmon highlights the utility of genomics for informing conservation. Science Advances. 2017;3(8). doi: 10.1126/sciadv.1603198.

179. Anonymous. Insights from Genetic Analyses into Stock Specific Early Ocean Migration Behavior of Juvenile Columbia River Steelhead. Fisheries Bulletin. Under Review.

180. Welch DW, Ishida Y, Nagasawa K, Eveson JP. Thermal limits on the ocean distribution of steelhead trout (Oncorhynchus mykiss). N Pac Anadr Fish Comm Bull. 1998;1:396-404.

181. Ward BR, Slaney PA. Life history and smolt-to-adult survival of Keogh River steelhead trout (Salmo gairdneri) and the relationship to smolt size. Can J Fish Aquat Sci. 1988;45(7):1110-22.

182. Welch DW, Ward BR, Smith BD, Evenson JP. Influence of the 1990 Ocean Climate Shift on British Columbia Steelhead (Oncorhynchus mykiss) and Coho (O.kisutch) Populations. NPAFC. 1998;310:17p.

183. Smith BD, Ward BR, Welch DW. Trends in British Columbia steelhead (Oncorhynchus mykiss) abundance indexed by angler success. Can J Fish Aquat Sci. 2000;57:1-16. doi: 10.1139/f99-254.

184. Welch DW, Ward BR, Smith BD, Whitney F. Changes associated with the 1989-90 ocean climate shift, and effects on British Columbia steelhead (O. mykiss) populations. In: Fisheries and Oceans Canada, editor.: Pacific Stock Assessment Review Committee (PSARC) Paper S97-7; 1997.

185. Neilson J, Taylor E. Emergency assessments of the Steelhead Trout (Oncorhynchus mykiss): Thompson River and Chilcotin River populations (2018). (Available upon request by contacting the COSEWIC secretariat at). In: COSEWIC, editor. Ottawa: Government of Canada, Ministry of Environment and Climate Change; 2018. p. 26 pp.

186. Bradford MJ. Comparative review of Pacific salmon survival rates. Can J Fish Aquat Sci. 1995;52(6):1327-38. doi: 10.1139/f95-129.

## References

2. McCann J, Chockley B, Cooper E, Hsu B, Schaller H, Haeseker S, et al. Comparative Survival Study of PIT-tagged Spring/Summer/Fall Chinook, Summer Steelhead, and Sockeye. 2017 Annual Report. Portland, Oregon2017.

