## Supplementary material for "The coast-wide collapse in marine survival of west coast Chinook and steelhead: slow-moving catastrophe or deeper failure?"

**Table S2. Source populations for SAR estimates used in the study; map code is used in Figure 1, Race refers to adult run timing, Rear is either H (Hatchery) or W (Wild) , Jacks indicates whether precocious male returns are included in survival estimates, N is the sample size (years of data). Reach refers to the migration segment over which SARs were estimated (survival from smolt enumeration site to adult enumeration site). See Methods for discussion.**

### A. Chinook

| Region | Stock | Short Name | Map Code | Race | Rear | Smolt Age | Jacks | Reach | N | From | To | Source |
|---|---|---|---|---|---|---|---|---|---|---|---|---|
| AK | Alaska Spring | Alaska | 5 | Spring | H | 1 | Y | | 35 | 1978 | 2012 | PSC CWT |
| | Chilkat Spring | Chilkat | 1 | Spring | W | 1 | Y | | 12 | 2001 | 2012 | PSC CWT |
| | Unuk Spring | Unuk | 4 | Spring | W | 1 | Y | | 24 | 1984 | 2012 | PSC CWT |
| | Stikine River Spring | Stikine | 3 | Spring | W | 1 | Y | | 13 | 2000 | 2012 | PSC CWT |
| | Taku Spring | Taku | 2 | Spring | W | 1 | Y | | 27 | 1977 | 2012 | PSC CWT |
| NCBC | Atnarko River Summer | Atnarko | 8 | Summer | H | 0 | Y | | 23 | 1987 | 2011 | PSC CWT |
| | Kitsumkalum Yearlings | Kits | 6 | Summer | H | 1 | Y | | 11 | 2001 | 2012 | PSC CWT |
| WCVI | Robertson Creek | Robertson | 17 | Fall | H | 0 | Y | | 39 | 1974 | 2012 | PSC CWT |
| SOG | Big Qualicum | BigQual | 16 | Fall | H | 0 | Y | | 39 | 1974 | 2012 | PSC CWT |
| | Chilliwack Fall | Chilliwack | 19 | Fall | H | 0 | Y | | 31 | 1982 | 2012 | PSC CWT |
| | Cowichan | Cowichan | 22 | Fall | H | 0 | Y | | 25 | 1986 | 2012 | PSC CWT |
| | Dome Creek Spring | Dome | 7 | Spring | H | 1 | Y | | 16 | 1988 | 2004 | PSC CWT |
| | Harrison River Fall | Harrison | 18 | Fall | H | 0 | Y | | 30 | 1982 | 2012 | PSC CWT |
| | Lower Shuswap River Summers | LowShuswap | 11 | Summer | H | 0 | Y | | 28 | 1985 | 2012 | PSC CWT |
| | Middle Shuswap Summers | MidShuswap | 12 | Summer | H | 0 | Y | | 4 | 2009 | 2012 | PSC CWT |
| | Nanaimo River Fall | Nanaimo | 20 | Fall | H | 0 | Y | | 19 | 1980 | 2005 | PSC CWT |
| | Nicola River Spring | Nicola | 13 | Spring | H | 1 | Y | | 27 | 1987 | 2013 | PSC CWT |
| | Phillips River Fall | Phillips | 10 | Fall | H | 0 | Y | | 3 | 2010 | 2012 | PSC CWT |
| | Puntledge Summer | Puntledge | 15 | Summer | H | 0 | Y | | 36 | 1976 | 2012 | PSC CWT |
| | Quinsam Fall | Quinsam | 14 | Fall | H | 0 | Y | | 38 | 1975 | 2012 | PSC CWT |
| PS | George Adams Fall Fingerling | GeorgeAdam | 44 | Summer/Fall | H | 0 | Y | | 33 | 1973 | 2011 | PSC CWT |

| | | | | | | | | | | | |
|---|---|---|---|---|---|---|---|---|---|---|---|
| | Nisqually Fall Fingerling | Nisqually | 51 | Summer/Fall | H | 0 | Y | 13 | 1999 | 2011 | PSC CWT |
| | Nooksack Spring Fingerling | NookFng | 21 | Spring | H | 0 | Y | 21 | 1990 | 2012 | PSC CWT |
| | Nooksack Spring Yearling | NookYrl | 21 | Spring | H | 1 | Y | 13 | 1983 | 1998 | PSC CWT |
| | Samish Fall Fingerling | Samish | 23 | Summer/Fall | H | 0 | Y | 29 | 1975 | 2011 | PSC CWT |
| | Skagit Spring Fingerling | SkagSpFng | 24 | Spring | H | 0 | Y | 19 | 1987 | 2012 | PSC CWT |
| | Skagit Spring Yearling | SkagSpYrl | 24 | Spring | H | 1 | Y | 26 | 1983 | 2012 | PSC CWT |
| | Skagit Summer Fingerling | SkagitSu | 24 | Summer | H | 0 | Y | 17 | 1995 | 2011 | PSC CWT |
| | Skykomish Fall Fingerling | Skykomish | 34 | Summer/Fall | H | 0 | Y | 11 | 2001 | 2011 | PSC CWT |
| | South Puget Sound Fall Fingerling | SPugetFng | 52 | Summer/Fall | H | 0 | Y | 38 | 1972 | 2011 | PSC CWT |
| | Squaxin Pens Fall Yearling | Squaxin | 46 | Fall | H | 1 | Y | 10 | 1987 | 1998 | PSC CWT |
| | Stillaguamish Fall Fingerling | Stillag | 27 | Summer/Fall | H | 0 | Y | 26 | 1981 | 2011 | PSC CWT |
| | University of Washington Accelerated | UW | 37 | Fall | H | 0 | Y | 10 | 1976 | 1985 | PSC CWT |
| | White River Spring Yearling | White | 48 | Spring | H | 1 | Y | 11 | 1976 | 2012 | PSC CWT |
| WAC | Elwha Fall Fingerling | Elwha | 29 | Summer/Fall | H | 0 | Y | 12 | 1983 | 1995 | PSC CWT |
| | Hoko Fall Fingerling | Hoko | 28 | Fall | H | 0 | Y | 25 | 1986 | 2011 | PSC CWT |
| | Queets Fall Fingerling | Queets | 40 | Fall | H | 0 | Y | 33 | 1978 | 2011 | PSC CWT |
| | Sooes Fall Fingerling | Sooes | 26 | Fall | H | 0 | Y | 25 | 1986 | 2011 | PSC CWT |
| LCOL | Columbia Lower River Hatchery | LowCol | 64 | Fall Tule | H | 0 | Y | 35 | 1977 | 2011 | PSC CWT |
| | Cowlitz Tule | Cowlitz | 65 | Fall Tule | H | 0 | Y | 34 | 1978 | 2011 | PSC CWT |
| | Lewis River Wild | Lewis | 71 | Fall Bright | W | 0 | Y | 31 | 1978 | 2011 | PSC CWT |
| | Willamette Spring | Willamette | 92 | Spring | H | 1 | Y | 35 | 1977 | 2011 | PSC CWT |

| | | | | | | | | | | | | |
|---|---|---|---|---|---|---|---|---|---|---|---|---|
| MCOL | Carson Hatchery Spring Chinook | Carson | 70 | Spr | H | 1 | N | Rel to BOA | 14 | 2000 | 2013 | FPC PIT |
| | Carson Hatchery Spring Chinook | Carson | 70 | Spr | H | 1 | Y | Rel to BOA | 1 | 2014 | 2014 | FPC PIT |
| | Cle Elum Hatchery Spring Chinook | CleElum | 47 | Spr | H | 1 | Y | MCN to MCA | 13 | 2002 | 2014 | FPC PIT |
| | Deschutes River Wild Fall Chinook | Deschutes | 80 | Fall | W | 0 | Y | Rel to BOA | 2 | 2011 | 2012 | FPC PIT |
| | Hanford Reach Wild Fall Chinook | Hanford | 57 | Fall | W | 0 | N | Rel to BOA | 9 | 2000 | 2010 | FPC PIT |
| | Hanford Reach Wild Fall Chinook | Hanford | 57 | Fall | W | 0 | Y | Rel to BOA | 2 | 2011 | 2012 | FPC PIT |
| | Hanford Wild | Hanford2 | 57 | Fall Bright | W | 0 | Y | | 25 | 1987 | 2011 | PSC CWT |
| | John Day River Wild Spring Chinook | JohnDay | 79 | Spr | W | 1 | Y | JDA to BOA | 15 | 2000 | 2014 | FPC PIT |
| | Little White Salmon Hatchery Fall Chinook | LtlWhite | 75 | Fall | H | 0 | N | Rel to BOA | 3 | 2008 | 2010 | FPC PIT |
| | Little White Salmon Hatchery Fall Chinook | LtlWhite | 75 | Fall | H | 0 | Y | Rel to BOA | 2 | 2011 | 2012 | FPC PIT |
| | Spring Creek Hatchery Fall Chinook (March Release) | SprMarch | 74 | Fall | H | 0 | N | Rel to BOA | 5 | 2008 | 2012 | FPC PIT |
| | Spring Creek Hatchery Fall Chinook (May Release) | SprMay | 74 | Fall | H | 0 | N | Rel to BOA | 5 | 2008 | 2012 | FPC PIT |
| | Spring Creek Tule | SprTule | 74 | Fall Tule | H | 0 | Y | | 39 | 1973 | 2011 | PSC CWT |
| | Upriver Bright | Upriver | 55 | Fall Bright | H | 0 | Y | | 36 | 1976 | 2011 | PSC CWT |
| | Warm Springs Hatchery Spring Chinook | Warm | 87 | Spr | H | 1 | N | Rel to BOA | 7 | 2007 | 2013 | FPC PIT |
| | Warm Springs Hatchery Spring Chinook | Warm | 87 | Spr | H | 1 | Y | Rel to BOA | 1 | 2014 | 2014 | FPC PIT |
| | Yakima River Wild Spring Chinook | Yakima | 63 | Spr | W | 1 | Y | MCN to MCA | 11 | 2002 | 2013 | FPC PIT |

| | | | | | | | | | | | | |
|---|---|---|---|---|---|---|---|---|---|---|---|---|
| UCOL | Columbia Summers | Columbia | 30 | Summer | H | 0 | Y | | 31 | 1976 | 2011 | PSC CWT |
| | Combined Hatch Wild Spring Chinook tagged at Rock Island Dam | RockIsSp | 42 | Spr | HW | 1 | Y | Rel to BOA | 14 | 2000 | 2014 | FPC PIT |
| | Combined Hatch Wild Summer Chinook tagged at Rock Island Dam | RockIsSu | 42 | Sum | HW | 0 | Y | Rel to BOA | 15 | 2000 | 2014 | FPC PIT |
| | Entiat and Methow River Wild Spring Chinook | EntMeth | 35 | Spr | W | 1 | Y | RRE to BOA | 7 | 2008 | 2014 | FPC PIT |
| | Entiat Hatchery Summer Chinook | Entiat | 36 | Sum | H | 0 | Y | RRE to BOA | 4 | 2011 | 2014 | FPC PIT |
| | Leavenworth Hatchery Spring Chinook | Leaven | 39 | Spr | H | 1 | Y | MCN to BOA | 15 | 2000 | 2014 | FPC PIT |
| | Mid-Columbia River Hatchery Spring Chinook | MColHSp | 30 | Spr | H | 1 | Y | first to PRA | 13 | 1972 | 1984 | Raymond |
| | Mid-Columbia River Wild Hatchery Combined Summer Chinook | MColWHSu | 30 | Sum | HW | 0 | Y | first to PRA | 16 | 1968 | 1983 | Raymond |
| | Mid-Columbia River Wild Spring Chinook | MColWSp | 30 | Spr | W | 1 | Y | first to PRA | 23 | 1962 | 1984 | Raymond |
| | Mid-Columbia River Wild Summer Chinook | MColWSu | 30 | Sum | W | 0 | Y | first to PRA | 7 | 1962 | 1968 | Raymond |
| | Upper Columbia River (above Wells Dam) Wild Summer Chinook | AboveWells | 30 | Sum | W | 0 | Y | RRE to BOA | 3 | 2011 | 2013 | FPC PIT |
| | Wenatchee River Wild Spring Chinook | Wenatchee | 41 | Spr | W | 1 | Y | MCN to BOA | 8 | 2007 | 2014 | FPC PIT |

| | | | | | | | | | | | | |
|---|---|---|---|---|---|---|---|---|---|---|---|---|
| | Winthrop Hatchery Spring Chinook | Winthrop | 25 | Spr | H | 1 | Y | RRE to BOA | 6 | 2009 | 2014 | FPC PIT |
| SNAK | Catherine Creek Hatchery Spring Chinook | Catherine | 83 | Spr | H | 1 | Y | LGR to GRA | 14 | 2001 | 2014 | FPC PIT |
| | Clearwater Hatchery Spring Chinook | ClearHSp | 59 | Spr | H | 1 | Y | LGR to GRA | 9 | 2006 | 2014 | FPC PIT |
| | Clearwater Hatchery Summer Chinook | ClearHSu | 59 | Sum | H | 1 | Y | LGR to GRA | 4 | 2011 | 2014 | FPC PIT |
| | Clearwater River Wild Spring Chinook | ClearWSp | 61 | Spr | W | 1 | Y | LGR to GRA | 9 | 2006 | 2014 | FPC PIT |
| | Dworshak Hatchery Fall Chinook at Snake River (Surrogates) | Dworshak | 59 | Fall | H | 0 | Y | LGR to GRA | 5 | 2006 | 2011 | FPC PIT |
| | Dworshak Hatchery Spring Chinook | Dworshak | 59 | Spr | H | 1 | Y | LGR to GRA | 18 | 1997 | 2014 | FPC PIT |
| | Grande Ronde River Hatchery Fall Chinook | GrndRonde | 73 | Fall | H | 0 | Y | LGR to GRA | 6 | 2006 | 2012 | FPC PIT |
| | Grande Ronde River Wild Spring Chinook | GrndRonde | 73 | Spr | W | 1 | Y | LGR to GRA | 9 | 2006 | 2014 | FPC PIT |
| | Imnaha Hatchery Summer Chinook | ImnahaH | 72 | Sum | H | 1 | Y | LGR to GRA | 18 | 1997 | 2014 | FPC PIT |
| | Imnaha River Wild Summer Chinook | ImnahaW | 72 | Sum | W | 1 | Y | LGR to GRA | 9 | 2006 | 2014 | FPC PIT |
| | Lyons Ferry | Lyons | 56 | Fall Bright | H | 0 | Y | | 18 | 1985 | 2011 | PSC CWT |
| | Lyons Ferry Hatchery Fall Chinook at Big Canyon Creek AP | LyonsBgCan | 60 | Fall | H | 0 | Y | LGR to GRA | 6 | 2006 | 2012 | FPC PIT |
| | Lyons Ferry Hatchery Fall Chinook at Captain | LyonsCptJ | 66 | Fall | H | 0 | Y | LGR to GRA | 5 | 2008 | 2012 | FPC PIT |

| | | | | | | | | | | | |
|---|---|---|---|---|---|---|---|---|---|---|---|
| John Rapids AP | | | | | | | | | | | |
| Lyons Ferry Hatchery Fall Chinook at Pittsburg Landing AP | LyonsPitts | 77 | Fall | H | 0 | Y | LGR to GRA | 6 | 2006 | 2012 | FPC PIT |
| Lyons Ferry Hatchery Fall Chinook at Snake River | LyonsSnR | 56 | Fall | H | 0 | Y | LGR to GRA | 6 | 2006 | 2012 | FPC PIT |
| McCall Hatchery Summer Chinook | McCall | 86 | Sum | H | 1 | Y | LGR to GRA | 18 | 1997 | 2014 | FPC PIT |
| Middle Fork Salmon River Wild Spring Summer Chinook | MidSalmon | 90 | SpSu | W | 1 | Y | LGR to GRA | 9 | 2006 | 2014 | FPC PIT |
| Nez Perce Hatchery Fall Chinook at Cedar Flats AP | NezPerCedr | 67 | Fall | H | 0 | Y | LGR to GRA | 3 | 2010 | 2012 | FPC PIT |
| Nez Perce Hatchery Fall Chinook at Lukes Gulch AP | NezPerLuke | 68 | Fall | H | 0 | Y | LGR to GRA | 3 | 2010 | 2012 | FPC PIT |
| Oxbow Hatchery Fall Chinook below Hells Canyon Dam | Oxbow | 85 | Fall | H | 0 | Y | LGR to GRA | 4 | 2008 | 2012 | FPC PIT |
| Pahsimeroi Hatchery Summer Chinook | Pahsimeroi | 88 | Sum | H | 1 | Y | LGR to GRA | 7 | 2008 | 2014 | FPC PIT |
| Rapid River Hatchery Spring Chinook | Rapid | 78 | Spr | H | 1 | Y | LGR to GRA | 18 | 1997 | 2014 | FPC PIT |
| Sawtooth Hatchery Spring Chinook | Sawtooth | 91 | Spr | H | 1 | Y | LGR to GRA | 8 | 2007 | 2014 | FPC PIT |
| Snake River Hatchery Spring Chinook | SnakeHSp | 54 | Spr | H | 1 | Y | GOJ to IHA | 5 | 1970 | 1974 | Raymond |

| | | | | | | | | | | | |
|---|---|---|---|---|---|---|---|---|---|---|---|
| Snake River Hatchery Spring Chinook | SnakeHSp | 54 | Spr | H | 1 | Y | ICH to IHA | 3 | 1966 | 1968 | Raymond |
| Snake River Hatchery Spring Chinook | SnakeHSp | 54 | Spr | H | 1 | Y | LGR to IHA | 10 | 1975 | 1984 | Raymond |
| Snake River Hatchery Spring Chinook | SnakeHSp | 54 | Spr | H | 1 | Y | LMJ to IHA | 1 | 1969 | 1969 | Raymond |
| Snake River Wild Fall Chinook | SnakeWFa | 54 | Fall | W | 0 | Y | LGR to GRA | 4 | 2006 | 2011 | FPC PIT |
| Snake River Wild Spring Chinook | SnakeWSp | 54 | Spr | W | 1 | Y | GOJ to IHA | 5 | 1970 | 1974 | Raymond |
| Snake River Wild Spring Chinook | SnakeWSp | 54 | Spr | W | 1 | Y | ICH to IHA | 5 | 1964 | 1968 | Raymond |
| Snake River Wild Spring Chinook | SnakeWSp | 54 | Spr | W | 1 | Y | LGR to IHA | 10 | 1975 | 1984 | Raymond |
| Snake River Wild Spring Chinook | SnakeWSp | 54 | Spr | W | 1 | Y | LMJ to IHA | 1 | 1969 | 1969 | Raymond |
| Snake River Wild Spring Summer Chinook | SnakeWSpSu | 54 | SpSu | W | 1 | Y | LGR to GRA | 21 | 1994 | 2014 | FPC PIT |
| Snake River Wild Summer Chinook | SnakeWSu | 54 | Sum | W | 1 | Y | GOJ to IHA | 5 | 1970 | 1974 | Raymond |
| Snake River Wild Summer Chinook | SnakeWSu | 54 | Sum | W | 1 | Y | ICH to IHA | 5 | 1964 | 1968 | Raymond |
| Snake River Wild Summer Chinook | SnakeWSu | 54 | Sum | W | 1 | Y | LGR to IHA | 10 | 1975 | 1984 | Raymond |
| Snake River Wild Summer Chinook | SnakeWSu | 54 | Sum | W | 1 | Y | LMJ to IHA | 1 | 1969 | 1969 | Raymond |
| South Fork Salmon River Wild Spring Summer Chinook | SthSalmon | 89 | SpSu | W | 1 | Y | LGR to GRA | 9 | 2006 | 2014 | FPC PIT |
| Umatilla Irrigon Hatchery Fall Chinook below Hells Canyon Dam | UmaIrri | 69 | Fall | H | 0 | Y | LGR to GRA | 6 | 2006 | 2012 | FPC PIT |
| Upper Salmon River Wild Spring Summer Chinook | UpSalmon | 82 | SpSu | W | 1 | Y | LGR to GRA | 9 | 2006 | 2014 | FPC PIT |

| | | | | | | | | | | | | |
|---|---|---|---|---|---|---|---|---|---|---|---|---|
| ORC | Elk River | Elk | 93 | Fall | H | 0 | Y | | 34 | 1978 | 2011 | PSC CWT |
| | Salmon River | Salmon | 84 | Fall | H | 0 | Y | | 34 | 1977 | 2011 | PSC CWT |

### B. Steelhead

| Region | Stock | Short Name | Map Code | Race | Rear | Smolt Age | Jacks | Reach | N | From | To | Source |
|---|---|---|---|---|---|---|---|---|---|---|---|---|
| KEOG | Keogh | Keogh | 9 | Win | W | NA | Y | | 37 | 1977 | 2013 | Davies |
| PS | Big Beef Creek | BigBeef | 38 | Win | W | NA | Y | | 6 | 2005 | 2010 | WDFW |
| | Green R. summer | GreenSu | 43 | Sum | H | NA | Y | | 19 | 1993 | 2011 | WDFW |
| | Green R. winter | GreenWn | 43 | Win | H | NA | Y | | 29 | 1982 | 2010 | WDFW |
| | Nisqually River | Nisqually | 51 | Win | W | NA | Y | | 3 | 2009 | 2011 | WDFW |
| | Nooksack R. winter | Nooksack | 21 | Win | H | NA | Y | | 13 | 1999 | 2011 | WDFW |
| | Puyallup R. winter | Puyallup | 49 | Win | H | NA | Y | | 23 | 1984 | 2006 | WDFW |
| | Samish R. winter | Samish | 23 | Win | H | NA | Y | | 3 | 1977 | 1979 | WDFW |
| | Skagit R. winter | Skagit | 24 | Win | H | NA | Y | | 31 | 1982 | 2012 | WDFW |
| | Snohomish R. summer | SnohoSu | 33 | Sum | H | NA | Y | | 18 | 1994 | 2011 | WDFW |
| | Snohomish R. winter | SnohoWn | 33 | Win | H | NA | Y | | 25 | 1986 | 2010 | WDFW |
| | Snow Creek | Snow | 31 | Win | W | NA | Y | | 36 | 1978 | 2013 | WDFW |
| | Stillaguamish R. summer | StillagSu | 27 | Sum | H | NA | Y | | 16 | 1996 | 2011 | WDFW |
| | Stillaguamish R. winter | StillagWn | 27 | Win | H | NA | Y | | 17 | 1994 | 2010 | WDFW |
| WAC | Chehalis winter | Chehalis | 50 | Win | H | NA | Y | | 32 | 1981 | 2012 | WDFW |
| | Elwha R. winter | Elwha | 29 | Win | H | NA | Y | | 15 | 1985 | 2001 | WDFW |
| | Humptulips R. summer | HumptulSu | 45 | Sum | H | NA | Y | | 10 | 1995 | 2008 | WDFW |
| | Humptulips R. winter | HumptulWn | 45 | Win | H | NA | Y | | 36 | 1977 | 2012 | WDFW |
| | Quillayute R. summer | QuilSu | 32 | Sum | H | NA | Y | | 13 | 1999 | 2011 | WDFW |
| | Quillayute R. winter | QuilWn | 32 | Win | H | NA | Y | | 30 | 1982 | 2011 | WDFW |
| | Willapa R. winter | Willapa | 58 | Win | H | NA | Y | | 17 | 1994 | 2010 | WDFW |
| | Wynoochee R. summer | Wynoochee | 53 | Sum | H | NA | Y | | 16 | 1994 | 2009 | WDFW |
| MCOL | Deschutes River Wild Steelhead | Deschutes | 80 | Sum | W | NA | Y | BON to BOA | 8 | 2006 | 2014 | FPC PIT |

| | | | | | | | | | | | | |
|---|---|---|---|---|---|---|---|---|---|---|---|---|
| | John Day River Wild Steelhead | JohnDay | 79 | Sum | W | NA | Y | JDA to BOA | 11 | 2004 | 2014 | FPC PIT |
| | Yakima River Wild Steelhead | Yakima | 63 | Sum | W | NA | Y | MCN to MCA | 13 | 2002 | 2014 | FPC PIT |
| UCOL | Combined Hatch Wild Steelhead tagged at Rock Island Dam | RockIs | 42 | Sum | HW | NA | Y | Rel to BOA | 14 | 2000 | 2014 | FPC PIT |
| | Eastbank and Chelan Hatchery Steelhead at Wenatchee River | EastbkChel | 41 | Sum | H | NA | Y | MCN to BOA | 12 | 2003 | 2014 | FPC PIT |
| | Entiat and Methow River Wild Steelhead | EntMeth | 35 | Sum | W | NA | Y | RRE to BOA | 7 | 2008 | 2014 | FPC PIT |
| | Mid-Columbia River Wild Hatchery Combined Steelhead | MCol | 30 | NA | HW | NA | Y | first to PRA | 23 | 1962 | 1984 | Raymond |
| | Wenatchee Entiat and Methow River Wild Steelhead | WenEntMeth | 35 | Sum | W | NA | Y | MCN to BOA | 9 | 2006 | 2014 | FPC PIT |
| SNAK | Clearwater River Hatchery Steelhead B-Run | ClearH | 59 | Sum | H | NA | Y | LGR to GRA | 7 | 2008 | 2014 | FPC PIT |
| | Clearwater River Wild Steelhead A-Run | ClearW | 59 | Sum | W | NA | Y | LGR to GRA | 9 | 2006 | 2014 | FPC PIT |
| | Grande Ronde River Hatchery Steelhead A-Run | GrndRondeH | 73 | Sum | H | NA | Y | LGR to GRA | 7 | 2008 | 2014 | FPC PIT |
| | Grande Ronde River Wild Steelhead A-Run | GrndRondeW | 73 | Sum | W | NA | Y | LGR to GRA | 9 | 2006 | 2014 | FPC PIT |
| | Hells Canyon Hatchery Steelhead A-Run | HellsC | 81 | Sum | H | NA | Y | LGR to GRA | 6 | 2009 | 2014 | FPC PIT |
| | Imnaha River Hatchery Steelhead A-Run | ImnahaH | 72 | Sum | H | NA | Y | LGR to GRA | 7 | 2008 | 2014 | FPC PIT |
| | Imnaha River Wild Steelhead A-Run | ImnahaW | 72 | Sum | W | NA | Y | LGR to GRA | 9 | 2006 | 2014 | FPC PIT |
| | Salmon River Hatchery Steelhead A-Run | SalmonHA | 76 | Sum | H | NA | Y | LGR to GRA | 7 | 2008 | 2014 | FPC PIT |
| | Salmon River Hatchery Steelhead B- | SalmonHB | 76 | Sum | H | NA | Y | LGR to GRA | 7 | 2008 | 2014 | FPC PIT |

| | | | | | | | | | | | |
|---|---|---|---|---|---|---|---|---|---|---|---|
| Run | | | | | | | | | | | |
| Salmon River Wild Steelhead A-Run | SalmonWA | 76 | Sum | W | NA | Y | LGR to GRA | 9 | 2006 | 2014 | FPC PIT |
| Snake River Hatchery Steelhead | SnakeH | 54 | NA | H | NA | Y | GOJ to IHA | 5 | 1970 | 1974 | Raymond |
| Snake River Hatchery Steelhead | SnakeH | 54 | NA | H | NA | Y | ICH to IHA | 2 | 1967 | 1968 | Raymond |
| Snake River Hatchery Steelhead | SnakeH | 54 | NA | H | NA | Y | LGR to IHA | 10 | 1975 | 1984 | Raymond |
| Snake River Hatchery Steelhead | SnakeH | 54 | NA | H | NA | Y | LMJ to IHA | 1 | 1969 | 1969 | Raymond |
| Snake River Hatchery Steelhead (all B-Run combined) | SnakeHB | 54 | Sum | H | NA | Y | LGR to GRA | 2 | 2013 | 2014 | FPC PIT |
| Snake River Hatchery Steelhead (all groups combined) | SnakeHC | 54 | Sum | H | NA | Y | LGR to GRA | 17 | 1997 | 2013 | FPC PIT |
| Snake River Wild Steelhead | SnakeW | 54 | NA | W | NA | Y | GOJ to IHA | 5 | 1970 | 1974 | Raymond |
| Snake River Wild Steelhead | SnakeW | 54 | NA | W | NA | Y | ICH to IHA | 5 | 1964 | 1968 | Raymond |
| Snake River Wild Steelhead | SnakeW | 54 | NA | W | NA | Y | LGR to IHA | 10 | 1975 | 1984 | Raymond |
| Snake River Wild Steelhead | SnakeW | 54 | NA | W | NA | Y | LMJ to IHA | 1 | 1969 | 1969 | Raymond |
| Snake River Wild Steelhead Aggregate | SnakeWAg | 54 | Sum | W | NA | Y | LGR to GRA | 18 | 1997 | 2014 | FPC PIT |
| Snake River Wild Steelhead A-Run | SnakeWA | 54 | Sum | W | NA | Y | LGR to GRA | 9 | 2006 | 2014 | FPC PIT |
| Snake River Wild Steelhead B-Run | SnakeWB | 54 | Sum | W | NA | Y | LGR to GRA | 9 | 2006 | 2014 | FPC PIT |

**Sources**

PSC CWT: Pacific Salmon Commission SAR database provided by G. Brown, Personal Communication. Department of Fisheries and Oceans, Government of Canada.

FPC PIT: McCann J, Chockley B, Cooper E, Hsu B, Schaller H, Haeseker S, Lessard R, Petrosky C, Copeland T, Tinus E, Van Dyke E, Storch A. Comparative Survival Study of PIT-tagged Spring/Summer/Fall Chinook, Summer Steelhead, and Sockeye. 2017 Annual Report. Comparative Survival Study Oversight Committee and Fish Passage Center, Portland, Oregon. 2017. Project No.: 19960200. Contract No.:74406. Sponsored by the Bonneville Power Administration. URL: http://www.fpc.org/documents/CSS/CSS_2017_Final_ver1-1.pdf

Davies: Investigators interested in accessing the Keogh SAR data should request these data from Dr Trevor Davies, Province of British Columbia,

WDFW: Updated dataset provided by Dr Neala Kendall (Pers. Comm.;) that is primarily reported in Kendall, N. W., G. W. Marston and M. M. Klungle (2017). "Declining patterns of Pacific Northwest steelhead trout (*Oncorhynchus mykiss*) adult abundance and smolt survival in the ocean." Canadian Journal of Fisheries and Aquatic Sciences **74**: 1275–1290. DOI: 10.1139/cjfas. Dr Kendall kindly provided an updated steelhead SAR dataset with data for more recent years than was available for her own publication.

Raymond: Raymond HL. Effects of Hydroelectric Development and Fisheries Enhancement on Spring and Summer Chinook Salmon and Steelhead in the Columbia River Basin. N Am J Fish Manag. 1988; 8(1): 1-24.

Vélez-Espino et al. 2011: Vélez-Espino LA, Willis J, Parken CK, Brown G. Cohort Analyses and New Developments for Coded Wire Tag Data of Atnarko River Chinook Salmon. Can. Manuscr. Rep. Fish. Aquat. Sci. 2011; 2958.

David Willis. Section Head, Coastal Operations, Fisheries and Oceans Canada.

Larry LaVoy. National Marine Fisheries Service, National Oceanic and Atmospheric Administration.

Tommy Garrison. Biometrician. Fisheries Management Department. Columbia River Inter-Tribal Fish Commission.

PSC (2015): Pacific Salmon Commission Joint Chinook Technical Committee. 2014 Exploitation Rate Analysis and Model Calibration Volume One. 2015. TCCHINOOK (15)-1 V.1.

PSC (2005): Pacific Salmon Commission Joint Chinook Technical Committee Report. Annual Exploitation Rate Analysis and Model Calibration. 2005. TCCHINOOK (05)-3.

**Notes on PSC Data**

- Atnarko River Summer Chinook: SARS estimates were available for subyearling and yearling stocks which are abbreviated as ATN and ATY by the PSC respectively. We retained ATN but excluded ATY because Atnarko is primarily a subyearling stock and the yearling releases are a hatchery management practise (Velez-Espino et al. 2011).
- Kitsumkalum River Chinook: SARS estimates were available for subyearling and yearling stocks which are abbreviated as KLM and KLY by the PSC respectively. We excluded KLM but retained KLY because Kitsumkalum is primarily a yearling stock. The subyearlings are released by the hatchery as fry and remain in the river an extra year until they migrate to sea at the same time as their sibling KLM fish (David Willis personnal communication May 2018).
- Lyons Ferry Chinook: SARS estimates were available for subyearling and yearling stocks which are abbreviated as LYF and LYY by the PSC respectively. We retained LYF but excluded LYY because Lyons Ferry is primarily a subyearling stock and the yearling releases are a hatchery management practise (Tommy Garrison personnal communication Jan 2018).

- Nooksack Spring Chinook: SARS estimates were available for subyearling and yearling stocks which are abbreviated as NKF and NKS by the PSC respectively. We retained both because the Nooksack stock naturally a mix of both life-history strategies (Larrie LaVoy personnal communication Jan 2018).
- Skagit Spring Chinook: SARS estimates were available for subyearling and yearling stocks which are abbreviated as SSF and SKS by the PSC respectively. We retained both because the Skagit stock naturally a mix of both life-history strategies (Larrie LaVoy personnal communication Jan 2018).
- South Puget Sound Fall Chinook: SARS estimates were available for subyearling and yearling stocks which are abbreviated as SPS and SPY by the PSC respectively. We retained SPS but excluded SPY because South Puget is primarily a subyearling stock and the yearling releases are a hatchery management practise (Larrie LaVoy personnal communication Jan 2018).
- Yearling/subyearling designations were taken from PSC (2015; Table 2.1) with the following exceptions: 1) Squaxin Pens Fall Chinook and University of Washington Accelerated Chinook were designated using PSC (2005; Table 2.1); and 2) we assumed Stikine Spring Chinook outmigrate as yearlings, and Phillips Fall Chinook outmigrate as subyearlings based on the typical behaviour for their adult run timing (neither stock was listed in PSC (2015)).
- SARS were available in several formats. We used survival data calculated as the sum of adults returning at all ages, uninflated for losses to natural mortality for Chinook remaining at sea for longer than two years because these values are most similar to the CSS PIT-tag based survival estimates.
- We excluded SARS estimates for ocean entry years with incomplete adult returns.

**Notes on CSS Data**

- For most stocks, SARS are provided with and without jack returns, and with differing start and end points to fish enumeration. When available, we used the estimates that included jacks and that covered the largest portion of the migration. For some MCOL populations, the estimates that included jack returns were available only for the shorter migration segment. In these cases, we used the estimates for the longer migration segment excluding jacks. Includes Spring Creek Hatchery Fall Chinook (5 of 5 years), Little White Salmon Hatchery Fall Chinook (3 of 5 years), Carson Hatchery Spring Chinook (14 of 15 years), Warm Springs Hatchery Spring Chinook (7 of 8 years), and Hanford Reach Wild Fall Chinook (9 of 11 years).
- We excluded SARS estimates for ocean entry years with incomplete adult returns.
- SARS data are referenced to McCann et al. (2017) Appendix B, but were actually downloaded from the Fish Passage Center: http://www.fpc.org/survival/smolttoadult_queries.php. Where there were discrepancies between these data sources, we retained the estimates from the online source.
