## Supplementary material for "The coast-wide collapse in marine survival of west coast Chinook and steelhead: slow-moving catastrophe or deeper failure?"

**Table S1. Source populations for freshwater survival estimates used in the study. Rear is either H (Hatchery), W (Wild) or U (Unknown). Reach refers to the migration segment spanned by the survival estimates. N is sample size (years of data). Rel: release. The full dataset is available from the authors upon request without restriction.**

### A. Chinook

| Region | Stock | Rear | Smolt Age | Tag Type | Reach | N | From | To | Source |
|---|---|---|---|---|---|---|---|---|---|
| NIMP | Nimpkish | U | 1 | V7 | Rel to mouth: Rkm 8.5 to mouth | 1 | 2006 | 2006 | Welch et al. 2011, Figure 3 |
| SOG | Chilko | H | 1 | V5 | Rel to mouth | 1 | 2016 | 2016 | Rechisky et al. in prep |
| | Coldwater | H | 1 | V7 | Rel to mouth: Rkm 395 to mouth | 1 | 2006 | 2006 | Welch et al. 2011, Figure 3 |
| | Coldwater | U | 1 | V7, V9 | Rel to mouth: Rkm 331 to mouth | 1 | 2005 | 2005 | Welch et al. 2011, Figure 3 |
| | Nicola | H | 1 | V7 | Rel to mouth: Rkm 331 to mouth | 1 | 2005 | 2005 | Welch et al. 2011, Figure 3 |
| | Nicola | H | 1 | V9 | Rel to mouth: Rkm 368 to mouth | 1 | 2004 | 2004 | Welch et al. 2011, Figure 3 |
| | Spius | H | 1 | V7 | Rel to mouth: Rkm 355 to mouth | 1 | 2006 | 2006 | Welch et al. 2011, Figure 3 |
| COL | Columbia | HW | 0 | JSATS | LRE: BON to Rkm 8.3 | 1 | 2007 | 2007 | McComas et al. 2009, Table 5 and Table 4 |
| | Columbia | HW | 0 | JSATS | LRE: BON to Rkm 8.3 | 1 | 2005 | 2005 | McMichael et al. 2007, Table 4 |
| | Columbia | HW | 0 | JSATS | LRE: BON to Rkm 8.3 | 1 | 2006 | 2006 | McMichael et al. 2007, Table 5 |
| | Columbia | HW | 0 | JSATS | LRE: BON to Rkm 8.3 | 1 | 2010 | 2010 | McMichael et al. 2010, Table ES3 |
| | Columbia | HW | 0 | JSATS | LRE: Rkm 153 to Rkm 8.3 | 1 | 2010 | 2010 | McMichael et al. 2011, Table 3.10 |
| | Columbia | HW | 1 | JSATS | LRE: BON to Rkm 8.3 | 1 | 2007 | 2007 | McComas et al. 2009, Table 5 and Table 3 |
| | Columbia | HW | 1 | JSATS | LRE: BON to Rkm 8.3 | 1 | 2005 | 2005 | McMichael et al. 2007, Table 4 |
| | Columbia | HW | 1 | JSATS | LRE: BON to Rkm | 1 | 2006 | 2006 | McMichael et al. 2007, Table 5 |

| | | | | | | | | | |
|---|---|---|---|---|---|---|---|---|---|
| | | | | | 8.3 | | | | |
| | Columbia | HW | 1 | JSATS | LRE: BON to Rkm 8.3 | 1 | 2009 | 2009 | McMichael et al. 2010, Table ES1 |
| | Columbia | HW | 1 | JSATS | LRE: Rkm 153 to Rkm 8.3 | 1 | 2009 | 2009 | McMichael et al. 2011, Table 3.4 |
| | Columbia | HW | 1 | V7 | LRE: JDA to Rkm 8 | 1 | 2010 | 2010 | Rechisky et al. 2014, Table 3 |
| MCOL | Mid-Columbia | H | 1 | V7 | full river: Rel to Rkm 23 | 2 | 2008 | 2009 | Rechisky et al. 2013, Table 1, Figure 3 |
| | Mid-Columbia | H | 1 | V9 | full river: Rel to Rkm 40 | 1 | 2006 | 2006 | Rechisky et al. 2009, Table 1 |
| | Mid-Columbia | HW | 1 | V7 | LRE: BON to Rkm 8 | 1 | 2011 | 2011 | Rechisky et al. 2014, Table 3 |
| UCOL | Upper Columbia | HW | 1 | V7 | LRE: BON to Rkm 8 | 1 | 2011 | 2011 | Rechisky et al. 2014, Table 3 |
| SNAK | Catherine Creek Hatchery Spring Chinook | H | 1 | PIT | hydrosystem: LGR to BON | 16 | 2001 | 2016 | McCann et al. 2017, Appendix A |
| | Clearwater Hatchery Spring Chinook | H | 1 | PIT | hydrosystem: LGR to BON | 11 | 2006 | 2016 | McCann et al. 2017, Appendix A |
| | Clearwater Hatchery Summer Chinook | H | 1 | PIT | hydrosystem: LGR to BON | 6 | 2011 | 2016 | McCann et al. 2017, Appendix A |
| | Clearwater River Wild Spring Chinook | W | 1 | PIT | hydrosystem: LGR to BON | 3 | 2014 | 2016 | McCann et al. 2017, Appendix A |
| | Dworshak Hatchery Fall Chinook at Snake River (Surrogates) | H | 0 | PIT | hydrosystem: LGR to BON | 5 | 2006 | 2011 | McCann et al. 2017, Appendix A |
| | Dworshak Hatchery Spring Chinook | H | 1 | PIT | hydrosystem: LGR to BON | 20 | 1997 | 2016 | McCann et al. 2017, Appendix A |
| | Grande Ronde River Hatchery Fall Chinook | H | 0 | PIT | hydrosystem: LGR to BON | 6 | 2006 | 2012 | McCann et al. 2017, Appendix A |
| | Grande Ronde River Wild Spring Chinook | W | 1 | PIT | hydrosystem: LGR to BON | 3 | 2014 | 2016 | McCann et al. 2017, Appendix A |
| | Imnaha Hatchery Summer Chinook | H | 1 | PIT | hydrosystem: LGR to BON | 20 | 1997 | 2016 | McCann et al. 2017, Appendix A |
| | Imnaha River Wild Summer Chinook | W | 1 | PIT | hydrosystem: LGR to BON | 4 | 2013 | 2016 | McCann et al. 2017, Appendix A |
| | Irrigon | H | 0 | PIT | Rel to BON: Cougar Ck to BON | 4 | 2008 | 2011 | Smith, Steve. Pers. comm. July 20, 2017 |
| | Kooskia Hatchery Spring Chinook | H | 1 | PIT | hydrosystem: LGR to BON | 3 | 2014 | 2016 | McCann et al. 2017, Appendix A |

| | | | | | | | | |
|---|---|---|---|---|---|---|---|---|
| Lyons Ferry | H | 0 | PIT | Rel to BON: Big Canyon Ck to BON | 6 | 2006 | 2012 | Smith, Steve. Pers. comm. July 20, 2017 |
| Lyons Ferry | H | 0 | PIT | Rel to BON: Captain John Rapids to BON | 5 | 2006 | 2012 | Smith, Steve. Pers. comm. July 20, 2017 |
| Lyons Ferry | H | 0 | PIT | Rel to BON: Cougar Ck to BON | 1 | 2006 | 2006 | Smith, Steve. Pers. comm. July 20, 2017 |
| Lyons Ferry | H | 0 | PIT | Rel to BON: Pittsburg Landing to BON | 6 | 2006 | 2012 | Smith, Steve. Pers. comm. July 20, 2017 |
| Lyons Ferry | H | 0 | PIT | Rel to BON: Snake R. at Couse Ck to BON | 4 | 2006 | 2012 | Smith, Steve. Pers. comm. July 20, 2017 |
| Lyons Ferry Hatchery Fall Chinook at Big Canyon Creek AP | H | 0 | PIT | hydrosystem: LGR to BON | 6 | 2006 | 2012 | McCann et al. 2017, Appendix A |
| Lyons Ferry Hatchery Fall Chinook at Captain John Rapids AP | H | 0 | PIT | hydrosystem: LGR to BON | 7 | 2008 | 2016 | McCann et al. 2017, Appendix A |
| Lyons Ferry Hatchery Fall Chinook at Pittsburg Landing AP | H | 0 | PIT | hydrosystem: LGR to BON | 8 | 2006 | 2016 | McCann et al. 2017, Appendix A |
| Lyons Ferry Hatchery Fall Chinook at Snake River | H | 0 | PIT | hydrosystem: LGR to BON | 6 | 2006 | 2012 | McCann et al. 2017, Appendix A |
| McCall Hatchery Summer Chinook | H | 1 | PIT | hydrosystem: LGR to BON | 21 | 1997 | 2016 | McCann et al. 2017, Appendix A |
| Middle Fork Salmon River Wild Spring_Summer Chinook | W | 1 | PIT | hydrosystem: LGR to BON | 3 | 2014 | 2016 | McCann et al. 2017, Appendix A |
| Nez Perce | H | 0 | PIT | Rel to BON: Cedar Flats to BON | 5 | 2006 | 2012 | Smith, Steve. Pers. comm. July 20, 2017 |
| Nez Perce | H | 0 | PIT | Rel to BON: Lukes Gulch to BON | 6 | 2006 | 2012 | Smith, Steve. Pers. comm. July 20, 2017 |
| Nez Perce | H | 0 | PIT | Rel to BON: Nez Perce Hatchery to BON | 5 | 2006 | 2012 | Smith, Steve. Pers. comm. July 20, 2017 |
| Nez Perce | H | 0 | PIT | Rel to BON: North Lapwai Valley to BON | 3 | 2010 | 2012 | Smith, Steve. Pers. comm. July 20, 2017 |
| Nez Perce Hatchery Fall Chinook | H | 0 | PIT | hydrosystem: LGR | 3 | 2010 | 2012 | McCann et al. 2017, Appendix A |

| | | | | | | | | |
|---|---|---|---|---|---|---|---|---|
| at Cedar Flats AP | | | | to BON | | | | |
| Nez Perce Hatchery Fall Chinook at Lukes Gulch AP | H | 0 | PIT | hydrosystem: LGR to BON | 3 | 2010 | 2012 | McCann et al. 2017, Appendix A |
| Oxbow | H | 0 | PIT | Rel to BON: Hells Canyon to BON | 5 | 2006 | 2012 | Smith, Steve. Pers. comm. July 20, 2017 |
| Oxbow Hatchery Fall Chinook below Hells Canyon Dam | H | 0 | PIT | hydrosystem: LGR to BON | 4 | 2008 | 2012 | McCann et al. 2017, Appendix A |
| Pahsimeroi Hatchery Summer Chinook | H | 1 | PIT | hydrosystem: LGR to BON | 9 | 2008 | 2016 | McCann et al. 2017, Appendix A |
| Rapid River Hatchery Spring Chinook | H | 1 | PIT | hydrosystem: LGR to BON | 20 | 1997 | 2016 | McCann et al. 2017, Appendix A |
| Sawtooth Hatchery Spring Chinook | H | 1 | PIT | hydrosystem: LGR to BON | 10 | 2007 | 2016 | McCann et al. 2017, Appendix A |
| Snake | H | 1 | JSATS | hydrosystem+LRE: LGR to Rkm 8 | 1 | 2008 | 2008 | Deitrich et al. 2016, Table 3 |
| Snake | H | 1 | V7 | full river: Rel to Rkm 23 | 2 | 2008 | 2009 | Rechisky et al. 2013, Table 1, Figure 3 |
| Snake | H | 1 | V9 | full river: Rel to Rkm 40 | 1 | 2006 | 2006 | Rechisky et al. 2009, Table 1 |
| Snake | HW | 1 | JSATS | hydrosystem+LRE: LGR to Rkm 8 | 1 | 2008 | 2008 | McMichael et al. 2010 |
| Snake | HW | 1 | JSATS | hydrosystem+LRE: LGR to Rkm 8.3 | 1 | 2006 | 2006 | McMichael et al. 2007, Table 5 |
| Snake | HW | 1 | PIT | hydrosystem: Snake trap to BON | 18 | 1999 | 2016 | Faulkner et al. 2017, Table 26 |
| Snake | HW | 1 | V7 | hydrosystem+LRE: LGR to Rkm 8 | 1 | 2010 | 2010 | Rechisky et al. 2014, Table 3 |
| Snake | U | 1 | V9 | LRE: BON to Rkm 8 | 1 | 2004 | 2004 | Welch et al. 2008; Clemens et al. 2009 |
| Snake | HW | 1 | V7 | LRE: BON to Rkm 8 | 1 | 2011 | 2011 | Rechisky et al. 2014, Table 3 |
| Snake River Wild Fall Chinook | W | 0 | PIT | hydrosystem: LGR to BON | 6 | 2006 | 2012 | McCann et al. 2017, Appendix A |
| Snake River Wild Spring_Summer Chinook | W | 1 | PIT | hydrosystem: LGR to BON | 23 | 1994 | 2016 | McCann et al. 2017, Appendix A |
| South Fork Salmon River Wild Spring_Summer Chinook | W | 1 | PIT | hydrosystem: LGR to BON | 3 | 2014 | 2016 | McCann et al. 2017, Appendix A |
| Umatilla | H | 0 | PIT | Rel to BON: Hells | 6 | 2006 | 2012 | Smith, Steve. Pers. comm. July 20, 2017 |

| | | | | | | | | | |
|---|---|---|---|---|---|---|---|---|---|
| | | | | | Canyon to BON | | | | |
| | Umatilla_Irrigon Hatchery Fall Chinook below Hells Canyon Dam | H | 0 | PIT | hydrosystem: LGR to BON | 6 | 2006 | 2012 | McCann et al. 2017, Appendix A |
| | Upper Salmon River Wild Spring_Summer Chinook | W | 1 | PIT | hydrosystem: LGR to BON | 3 | 2014 | 2016 | McCann et al. 2017, Appendix A |

### B. Steelhead

| | | | | | | | | | |
|---|---|---|---|---|---|---|---|---|---|
| KEOG | Keogh | H | | V9 | Rel to mouth: Rkm 0.3 to mouth | 2 | 2004 | 2005 | Welch et al. 2011, Figure 3 |
| | Keogh | W | | V9 | Rel to mouth: Rkm 0.3 to mouth | 2 | 2004 | 2006 | Welch et al. 2011, Figure 3 |
| SOG | Coldwater | W | | V7, V9 | Rel to mouth: Rkm 57 to mouth | 1 | 2006 | 2006 | Welch et al. 2011, Figure 3 |
| | Coldwater | W | | V9 | Rel to mouth: Rkm 31-51 to mouth | 2 | 2004 | 2005 | Welch et al. 2011, Figure 3 |
| | Cowichan | H | | V9 | Rel to mouth | 1 | 2006 | 2006 | Welch et al. 2011, Figure 3 |
| | Deadman | W | | V7, V9 | Rel to mouth: Rkm 363 to mouth | 1 | 2006 | 2006 | Welch et al. 2011, Figure 3 |
| | Deadman | W | | V9 | Rel to mouth: Rkm 342 to mouth | 1 | 2005 | 2005 | Welch et al. 2011, Figure 3 |
| | Englishman | W | | V9 | Rel to mouth: Rkm 2.5 to mouth | 3 | 2004 | 2006 | Welch et al. 2011, Figure 3 |
| | Seymour | H | | V7 | Rel to mouth | 1 | 2015 | 2015 | Healy et al. 2017, Page 7 |
| | Seymour | H | | V9 | Rel to mouth | 3 | 2006 | 2009 | Balfry et al. 2011, Table 3 |
| | Seymour | H | | V9 | Rel to mouth | 1 | 2007 | 2007 | Balfry et al. 2011, Table 3; Welch et al. 2011, Figure 3 |
| | Squamish | H | | V9 | Rel to mouth: Rkm 15 to mouth | 1 | 2007 | 2007 | Welch et al. 2011, Figure 3 |
| | Squamish | W | | V9 | Rel to mouth: Rkm 16 to mouth | 2 | 2004 | 2005 | Welch et al. 2011, Figure 3 |
| PS | Big Beef Creek | W | | V7, V9 | Rel to mouth: Rkm 0.1 to mouth | 1 | 2006 | 2009 | Moore et al. 2015, Figure 3 |
| | Dewatto | W | | V7 | Rel to mouth: Rkm 0.3 to mouth | 1 | 2007 | 2007 | Moore et al. 2015, Figure 3 |
| | Duckabush | H | | V7 | Rel to mouth: Rkm | 1 | 2009 | 2009 | Moore et al. 2015, Figure 3 |

| | | | | | | | | |
|---|---|---|---|---|---|---|---|---|
| | | | | 1.9 to mouth | | | | |
| | Green | H | V7 | Rel to mouth: Rkm 55 to mouth | 1 | 2006 | 2008 | Goetz et al. 2015, Table 3; Moore et al. 2015, Figure 3 |
| | Green | W | V7 | Rel to mouth: Rkm 54.5 to mouth | 1 | 2006 | 2009 | Goetz et al. 2015, Table 3; Moore et al. 2015, Figure 3 |
| | Hamma Hamma | H | V7, V9 | Rel to mouth: Rkm 2 to mouth | 1 | 2006 | 2007 | Moore et al. 2015, Figure 3 |
| | Nisqually | W | V7, V9 | Rel to mouth: Rkm 0-21 to mouth | 1 | 2006 | 2009 | Moore et al. 2015, Figure 3 |
| | Puyallup | H | V7 | Rel to mouth: Rkm 55.8 to mouth | 1 | 2006 | 2009 | Moore et al. 2015, Figure 3 |
| | Puyallup | W | V7 | Rel to mouth: Rkm 17 to mouth | 1 | 2006 | 2006 | Moore et al. 2015, Figure 3 |
| | Skagit | H | V7, V9 | Rel to mouth: Rkm 102 to mouth | 1 | 2008 | 2009 | Moore et al. 2015, Figure 3 |
| | Skagit | W | V7 | Rel to mouth: Rkm 10 to mouth | 1 | 2006 | 2009 | Moore et al. 2015, Figure 3 |
| | Skokomish | H | V7 | Rel to mouth: Rkm 13.5 to mouth | 1 | 2008 | 2009 | Moore et al. 2015, Figure 3 |
| | Skokomish | W | V7, V9 | Rel to mouth: Rkm 13.5 to mouth | 1 | 2006 | 2009 | Moore et al. 2015, Figure 3 |
| UCOL | Upper Columbia | U | PIT | hydrosystem+LRE: RIS to Rkm 75 | 7 | 2008 | 2014 | Hostetter et al. 2018, Figure 4 |
| | Upper Columbia | HW | JSATS | LRE: BON to Rkm 8.3 | 1 | 2009 | 2009 | McMichael et al 2010, Table ES2 |
| | Upper Columbia | HW | JSATS | LRE: BON to Rkm 8.3 | 1 | 2010 | 2010 | McMichael et al 2011, Table 3.7 |
| SNAK | Asotin River Wild Steelhead | W | PIT | hydrosystem: LGR to BON | 1 | 2014 | 2014 | McCann et al. 2017, Appendix A |
| | Clearwater River Hatchery Steelhead B-Run | H | PIT | hydrosystem: LGR to BON | 9 | 2008 | 2016 | McCann et al. 2017, Appendix A |
| | Clearwater River Wild Steelhead A-Run | W | PIT | hydrosystem: LGR to BON | 2 | 2013 | 2014 | McCann et al. 2017, Appendix A |
| | Grande Ronde River Hatchery Steelhead A-Run | H | PIT | hydrosystem: LGR to BON | 9 | 2008 | 2016 | McCann et al. 2017, Appendix A |
| | Grande Ronde River Wild Steelhead A-Run | W | PIT | hydrosystem: LGR to BON | 1 | 2014 | 2014 | McCann et al. 2017, Appendix A |

| | | | | | | | | |
|---|---|---|---|---|---|---|---|---|
| | Hells Canyon Hatchery Steelhead A-Run | H | PIT | hydrosystem: LGR to BON | 8 | 2009 | 2016 | McCann et al. 2017, Appendix A |
| | Imnaha River Hatchery Steelhead A-Run | H | PIT | hydrosystem: LGR to BON | 9 | 2008 | 2016 | McCann et al. 2017, Appendix A |
| | Imnaha River Wild Steelhead A-Run | W | PIT | hydrosystem: LGR to BON | 2 | 2013 | 2014 | McCann et al. 2017, Appendix A |
| | Salmon River Hatchery Steelhead A-Run | H | PIT | hydrosystem: LGR to BON | 9 | 2008 | 2016 | McCann et al. 2017, Appendix A |
| | Salmon River Hatchery Steelhead B-Run | H | PIT | hydrosystem: LGR to BON | 9 | 2008 | 2016 | McCann et al. 2017, Appendix A |
| | Salmon River Wild Steelhead A-Run | W | PIT | hydrosystem: LGR to BON | 1 | 2014 | 2014 | McCann et al. 2017, Appendix A |
| | Snake | HW | PIT | hydrosystem: Snake Trap to BON | 20 | 1997 | 2016 | Faulkner et al. 2017, Table 29 |
| | Snake | (blank ) | V9 | LRE: BON to Rkm 23 | 2 | 2002 | 2003 | Clemens et al. 2009; Welch et al. 2008, Table 1 |
| | Snake River Hatchery Steelhead (all A-Run combined) | H | PIT | hydrosystem: LGR to BON | 9 | 2008 | 2016 | McCann et al. 2017, Appendix A |
| | Snake River Hatchery Steelhead (all B-Run combined) | H | PIT | hydrosystem: LGR to BON | 9 | 2008 | 2016 | McCann et al. 2017, Appendix A |
| | Snake River Hatchery Steelhead (all groups combined) | H | PIT | hydrosystem: LGR to BON | 20 | 1997 | 2016 | McCann et al. 2017, Appendix A |
| | Snake River Wild Steelhead Aggregate | W | PIT | hydrosystem: LGR to BON | 20 | 1997 | 2016 | McCann et al. 2017, Appendix A |
| | Snake River Wild Steelhead A-Run | W | PIT | hydrosystem: LGR to BON | 2 | 2013 | 2014 | McCann et al. 2017, Appendix A |
| | Snake River Wild Steelhead B-Run | W | PIT | hydrosystem: LGR to BON | 2 | 2013 | 2014 | McCann et al. 2017, Appendix A |
| ORC | Alsea | W | V7 | Rel to mouth: Rkm 55 to mouth | 1 | 2009 | 2009 | Romer et al. 2013, Table 2 |
| | Alsea | W | V7, V9 | Rel to mouth: Rkm 55 to mouth | 1 | 2007 | 2007 | Romer et al. 2013, Table 2; Johnson et al. 2010 |
| | Nehalam | W | V7 | Rel to mouth: Rkm 33 to mouth | 1 | 2009 | 2009 | Romer et al. 2013, Table 2 |
| | Nehalam | W | V9 | Rel to mouth: Rkm 33 to mouth | 2 | 2001 | 2002 | Romer et al. 2013, Table 2; Clements et al. 2012, Table 2 |

**Notes**

Estimates from Moore et al. 2015 were averaged across years.
Nicola and Spius yearling Chinook: omitted from analysis because tag burden exceeded current best practises (>75% of smolts had fork lengths under 130 mm for V7-tagged and under 140 mm for V9-tagged).
Cowichan steelhead: omitted from analysis because this was the only estimate outside the Columbia River area where the migration segment did not terminate in the river mouth.

**Sources**

1. Balfry S, Welch DW, Atkinson J, Lill A, Vincent S. The Effect of Hatchery Release Strategy on Marine Migratory Behaviour and Apparent Survival of Seymour River Steelhead Smolts (Oncorhynchus mykiss). Bograd SJ, editor. PLoS ONE. 2011 Mar;6(3):e14779.
2. Clemens BJ, Clements SP, Karnowski MD, Jepsen DB, Gitelman AI, Schreck CB. Effects of Transportation and Other Factors on Survival Estimates of Juvenile Salmonids in the Unimpounded Lower Columbia River. Trans Am Fish Soc. 2009 Jan;138(1):169–88.
3. Clements S, Stahl T, Schreck CB. A comparison of the behavior and survival of juvenile coho salmon (*Oncorhynchus kisutch*) and steelhead trout (*O. mykiss*) in a small estuary system. Aquaculture. 2012 Sep;362–363:148–57.
4. Dietrich J, Eder K, Thompson D, Buchanan R, Skalski J, McMichael G, Fryer D, Loge F. Survival and transit of in-river and transported yearling Chinook salmon in the lower Columbia River and estuary. Fisheries Research. 2016 183:435–446.
5. Faulkner JR, Widener DL, Smith SG, Marsh DM, Zabel RW. Survival estimates for the passage of spring-migrating juvenile salmonids through Snake and Columbia River dams and reservoirs, 2016. Prep US Dep Energy Bonneville Power Adm Div Fish Wildl Portland Or. 2017;
6. Goetz FA, Jeanes E, Moore ME, Quinn TP. Comparative migratory behavior and survival of wild and hatchery steelhead (*Oncorhynchus mykiss*) smolts in riverine, estuarine, and marine habitats of Puget Sound, Washington. Environ Biol Fishes. 2015 Jan;98(1):357–75.
7. Harnish RA, Johnson GE, McMichael GA, Hughes MS, Ebberts BD. Effect of Migration Pathway on Travel Time and Survival of Acoustic-Tagged Juvenile Salmonids in the Columbia River Estuary. Trans Am Fish Soc. 2012 Mar;141(2):507–19.
8. Healy S, Hinch S, Porter A, Rechisky E, Welch D, Eliason E, et al. Route-specific movements and survival during early marine migration of hatchery steelhead Oncorhynchus mykiss smolts in coastal British Columbia. Mar Ecol Prog Ser. 2017 Aug 18;577:131–47.
9. Hostetter NJ, Gardner B, Evans AF, Cramer BM, Payton Q, Collis K, et al. Wanted dead or alive: a state-space mark–recapture–recovery model incorporating multiple recovery types and state uncertainty. Can J Fish Aquat Sci. 2018 Jul;75(7):1117–27.
10. Johnson SL, Power JH, Wilson DR, Ray J. A Comparison of the Survival and Migratory Behavior of Hatchery-Reared and Naturally Reared Steelhead Smolts in the Alsea River and Estuary, Oregon, using Acoustic Telemetry. North Am J Fish Manag. 2010 Feb;30(1):55–71.
11. McCann J, Chockley B, Cooper E, Hsu B, Schaller H, Haeseker S, Lessard R, Petrosky C, Copeland T, Tinus E, Van Dyke E, Storch A. Comparative Survival Study of PIT-tagged Spring/Summer/Fall Chinook, Summer Steelhead, and Sockeye. 2017 Annual Report. Comparative Survival Study Oversight Committee and Fish Passage Center, Portland, Oregon. 2017. Project No.: 19960200. Contract No.:74406. Sponsored by the Bonneville Power Administration. URL: http://www.fpc.org/documents/CSS/CSS_2017_Final_ver1-1.pdf
12. McComas RL, McMichael GA, Carter JA, Johnson GE, Gilbreath L, Everett JP, Smith SG, Carlson TJ, Matthews GM, Ferguson JW. A study of salmonid survival and behaviour through the Columbia River estuary using acoustic tags, 2007. 2009.
13. McMichael G, Harnish, RA, Bellgraph, BA, Carter JA, Ham KA, Titzler, PS, et al. Migratory Behavior and Survival of Juvenile Salmonids in the Lower Columbia River and Estuary in 2009. Pacific Northwest National Laboratory (PNNL), Richland, WA (US); 2010.

14. McMichael GA, Kim J, Skalski JR, Townsend RL, Deters KA, Ham KD, et al. Migratory behavior and survival of juvenile salmonids in the lower Columbia River, estuary, and plume in 2010. Pacific Northwest National Laboratory. 2011. PNNL-20443. Available from: http://137.161.203.100/environment/docs/afep/estuary/2010_Post-FCRPS_final.pdf
15. McMichael GA, Vucelick JA, Bellgraph BJ, Carlson TJ, McComas RL, Gilbreath L, Smith SG, Sandford B, Matthews G, Ferguson JW. Technical Memo: A study to estimate salmonid survival through the Columbia River estuary using acoustic tags, 2005 and 2006 synthesis report. Pacific Northwest National Laboratory. 2007. PNNL-SA-54927.
16. Moore ME, Berejikian BA, Goetz FA, Berger AG, Hodgson SS, Connor EJ, et al. Multi-population analysis of Puget Sound steelhead survival and migration behavior. Mar Ecol Prog Ser. 2015;537:217.
17. Rechisky EL, Welch DW, Porter AD, Jacobs MC, Ladouceur A. Experimental measurement of hydrosystem-induced delayed mortality in juvenile Snake River spring Chinook salmon (*Oncorhynchus tshawytscha*) using a large-scale acoustic array. Can J Fish Aquat Sci. 2009;66(7):1019–1024.
18. Rechisky EL, Welch DW, Porter AD, Jacobs-Scott MC, Winchell PM. Influence of multiple dam passage on survival of juvenile Chinook salmon in the Columbia River estuary and coastal ocean. Proc Natl Acad Sci [Internet]. 2013 Apr 1 [cited 2013 Apr 5]; Available from: http://www.pnas.org/cgi/doi/10.1073/pnas.1219910110
19. Rechisky E, Welch D, Porter A, Hess J, Narum S. Testing for delayed mortality effects in the early marine life history of Columbia River Basin yearling Chinook salmon. Mar Ecol Prog Ser. 2014 Jan 27;496:159–80.
20. Romer JD, Leblanc CA, Clements S, Ferguson JA, Kent ML, Noakes D, et al. Survival and behavior of juvenile steelhead trout (*Oncorhynchus mykiss*) in two estuaries in Oregon, USA. Environ Biol Fishes. 2013 Jul;96(7):849–63.
21. Welch DW, Rechisky EL, Melnychuk MC, Porter AD, Walters CJ, Clements S, et al. Survival of Migrating Salmon Smolts in Large Rivers With and Without Dams. PLoS Biol. 2008;6(10):e265.
22. Welch DW, Melnychuk MC, Payne JC, Rechisky EL, Porter AD, Jackson GD, et al. In situ measurement of coastal ocean movements and survival of juvenile Pacific salmon. Proc Natl Acad Sci. 2011 May; Available from: http://www.pnas.org/cgi/doi/10.1073/pnas.1014044108
