## Supplementary material for "The coast-wide collapse in marine survival of west coast Chinook and steelhead: slow-moving catastrophe or deeper failure?"

**SI-3: Direct Comparison of CWT and PIT tag-based SAR Estimates.**

We can identify two major systematic differences between the PIT- and CWT-based survival estimates: (1) PIT-based SAR estimate adult survival using those smolts that first survived to reach the topmost dam in the Columbia River hydrosystem and then return as adults to a dam where they are enumerated, while CWT-based SAR estimate adult survival from smolt release from the hatchery or (for wild fish) from an enumeration site in the river after smolt migration starts until adult return to the hatchery or spawning ground. (2) CWT-based survival estimates from the Pacific Salmon Commission (PSC) add the estimated sport and commercial harvest to the adult return to the river, while PIT tag-based survival estimates do not. The first difference (upstream losses not being included in the PIT tag-based analysis) will reduce CWT survival estimates relative to PIT tags, while the latter (harvest) will increase CWT survival.

Overall, it is difficult to predict exactly how the two estimates should combine to influence relative SAR for any given population. In this section we report our attempts to investigate this issue more thoroughly.

To assess the magnitude of the disparity between these two tagging methodologies, we searched the datasets for populations that had both PIT and CWT-based survival estimates for the same smolt outmigration year. We found three populations of subyearling (Fall) Chinook that had both CWT and PIT tag-based survival estimates (Table S3-1), but no matching populations for yearling (Spring) Chinook.

**Sub-Yearling Comparison**

We calculated a conversion factor between the CWT and PIT-based SAR estimates using linear regression on the matched pairs of survival estimates, with the PIT-based estimates as the independent and CWT-based estimates as the dependent variables (Figure S3-1; Table S3-1). The intercept from this relationship was not significantly different from zero (Table S3-2), so we ran the regression again with the intercept set to zero. The resulting slope (Table S3-2) was our conversion factor; the CWT-based SAR estimates were ca. 1.5X larger than the PIT-based estimates for the same stock and year of outmigration. That is, the combined effects of (a) migratory survival by downstream migrating smolts between the hatchery and the top-most dam in the hydrosystem, (b) upstream survival of migrating adults between the top-most dam and the spawning grounds (or enumeration site), and (c) sport and commercial harvest result in CWT-based SAR estimates averaging ca. 150% of the PIT tag-based estimates because the PIT tag-based SAR estimates do not take into account these processes.

**Table S3-1.** Stocks with SAR estimates for common outmigration years in both the CWT and PIT datasets. The “Stock” fields give the names as accessed from the source. For the PIT-based SARs, the “Stock” field was called “GroupDescription” on download. For PIT-based SAR where more than one release group is listed (i.e., for Spring Creek and Lyons Ferry) we used the mean SAR weighted by the sample size in the regression.

| Stock PIT-based | Stock CWT-based | H/W | Years |
|---|---|---|---|
| Hanford Reach Wild Fall Chinook | Hanford Wild | W | 2000-2001, 2003-2005, 2007-2011 |
| Spring Creek Hatchery Fall Chinook (March Release)<br>Spring Creek Hatchery Fall Chinook (April Release)<br>Spring Creek Hatchery Fall Chinook (May Release) | Spring Creek Tule | H | 2008-2011 |
| Lyons Ferry Hatchery Fall Chinook at Big Canyon Creek<br>Lyons Ferry Hatchery Fall Chinook at Captain John Rapids<br>Lyons Ferry Hatchery Fall Chinook at Pittsburg Landing<br>Lyons Ferry Hatchery Fall Chinook at Snake River | Lyons Ferry | H | 2006, 2008-2011 |

**Table S3-2**. Results from the linear regression between CWT (dependent) and PIT-tag (independent)-based SAR estimates for subyearling Chinook salmon with matched stocks and years of outmigration.

| Model | Variable | Estimate | SE | t value | P | $R^2$ |
|---|---|---|---|---|---|---|
| Model 1 | Intercept | 0.186 | 0.151 | 1.232 | 0.235 | 0.697 |
| | Slope | 1.341 | 0.206 | 6.507 | <0.001 | |
| Model 2 | Slope | 1.535 | 0.135 | 11.360 | <0.001 | 0.871 |

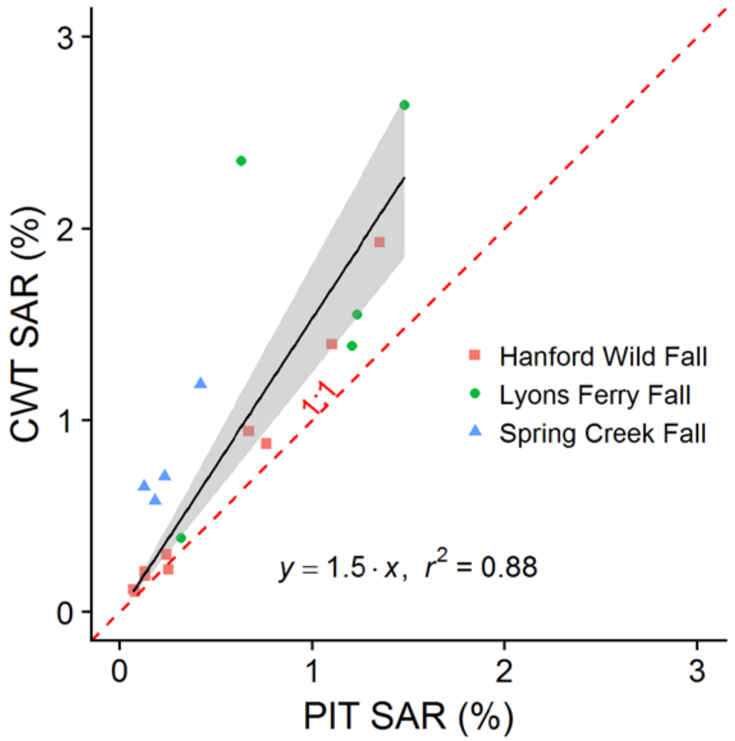


**Figure S3-1**. Comparative SARs of CWT and PIT tag based SARS estimates for subyearling Chinook salmon with matched stocks and years of outmigration. The red dashed line shows the expected 1:1 relationship if both PIT and CWT based survival estimates were exactly equivalent.

### Yearling Comparison

Because we found no matching populations of yearling Chinook in the PIT and CWT-based SAR datasets included in the main paper (see Methods), we approximated a conversion factor using SAR estimates from the University of Washington's DART database as an intermediate step (http://www.cbr.washington.edu/trends/index.php, data provided by Chris Van Holmes and Rich Townsend of U. Washington on Aug 17, 2017). The DART SAR estimates are also based on CWT recoveries, but for clarity we refer to them as "DART" and use "CWT" for the estimates from the PSC reported in the main body of the paper. The DART estimates are not inflated for harvest; methods are documented in (Skalski and Townsend 2005).

We found one population of yearling Chinook that had both DART and PIT tag-based SAR estimates, and a second population that had both DART and CWT SAR estimates (Table S3-3). For each population separately, we used linear regression on the matched pairs of survival estimates with the DART estimates as the dependent variable and the 1) PIT and 2) CWT-based estimates as the independent variable (Figure S3-2; Figure S3-3; Table S3-4). The intercept from both these relationships was not significantly different from zero, so in both cases we repeated the regressions with the intercept set to zero (Table S3-4).

The ratio of the resulting slopes allows us to calculate an overall yearling conversion factor. The DART vs PIT regression had a slope of 0.447, while the CWT vs DART regression yielded a slope of 1.478 (Table S3-4). Their product yields an overall CWT vs PIT tag relationship of 0.447x1.478=0.66. The CWT-based SAR estimates were thus only 2/3rds the PIT-based SAR estimates for yearling Chinook. This should be considered only a rough estimate of the "typical" difference between CWT and PIT tag-based SAR estimates because it is based on only two populations that were from different areas of the Columbia River system; however, the direction of the difference is roughly as expected because the PIT tag survival estimates (McCann et al. 2017) exclude smolt and adult losses above the dams while these are included in the PSC (& DART) SAR estimates.  Also, as expected, the DART CWT-based SAR estimates are lower than the PIT tag-based SAR estimates.  Neither the DART CWT-based SAR or the PIT tag-based SAR incorporate losses to commercial and sport harvest; the Pacific Salmon Commission's CWT-based SAR estimates used in the main paper include harvest in calculating SARs, which should bring them closer to the PIT tag-based SAR estimates.  However, as noted in the main paper, harvest rates of yearling (Spring) Chinook tend to be lower than for subyearling (Fall) Chinook because populations of the latter group remain exposed to fisheries on the continental shelf for several years.

**Table S3-3**. Stocks with SAR estimates for the same years of outmigration in the 1) PIT and DART, and 2) CWT and DART datasets. The "Stock" fields give the names as accessed from the source. For the PIT-based SARs, the "Stock" field was called "GroupDescription" on download. For the DART-based SARs, the "Stock" field was called "hatchery_location_name". Also for the DART-based SARs, when more than one release location is listed we used the mean SAR weighted by the sample size. Dworshak spring Chinook were released from Dworshak National Fish Hatchery for the PIT-based estimates; release locations were not provided for the CWT-based estimates.

| | | **DART** | | | |
|---|---|---|---|---|---|
| **Stock** | **PIT or CWT-based** | **Stock** | **Release location** | **H/W** | **Years** |
| PIT | Dworshak Hatchery Spring Chinook | Dworshak Nat. Hatchery | Dworshak Nat. Hatchery | H | 1997-2013 |
| CWT | Willamette Spring | Willamette Hatchery | Willamette R M FK-1<br>Santiam R S FK<br>Molalla R | H | 1989-1989, 1996-2011 |

**Table S3-4**. Results from the linear regression between DART (dependent) and CWT/PIT tag (independent)-based SAR estimates for yearling Chinook salmon with matched stocks and years of outmigration.

| Model | Variable | Estimate | SE | t value | p | $R^2$ |
|---|---|---|---|---|---|---|
| | | | **DART vs PIT** | | | |
| Model 1 | Intercept | -0.023 | 0.079 | -0.291 | 0.775 | 0.502 |
| | Slope | 0.477 | 0.115 | 4.141 | <0.001 | |
| Model 2 | Slope | 0.447 | 0.050 | 8.903 | <0.001 | 0.822 |
| | | | **CWT vs DART** | | | |
| Model 1 | Intercept | 0.268 | 0.211 | 1.270 | 0.222 | 0.522 |
| | Slope | 1.192 | 0.270 | 4.421 | <0.001 | |
| Model 2 | Slope | 1.478 | 0.150 | 9.842 | <0.001 | 0.842 |

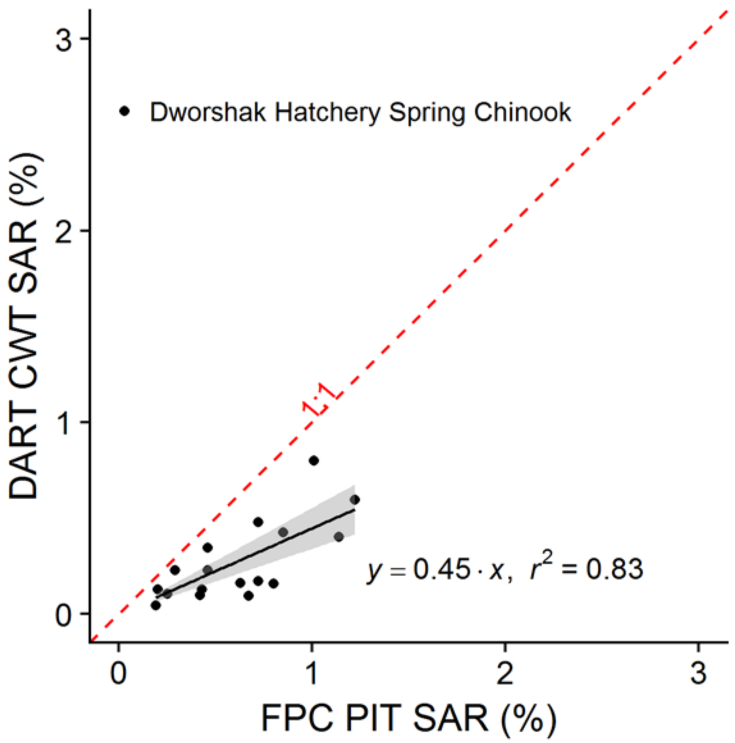


**Figure S3-2**. Comparative SAR of DART and PIT tag-based SAR estimates for yearling Chinook salmon with matched stocks and years of outmigration. The red dashed line shows the expected 1:1 relationship if both DART and PIT-based survival estimates were equivalent. The derived relationship indicates that the product of migratory survival by downstream migrating smolts after release from the hatchery until arrival at the top-most dam in the hydrosystem *and* multiplied by the survival of the upstream migrating adults above the top-most dam is 45%, as this component of the SAR is excluded from the published PIT tag SAR estimates (McCann et al. 2017).

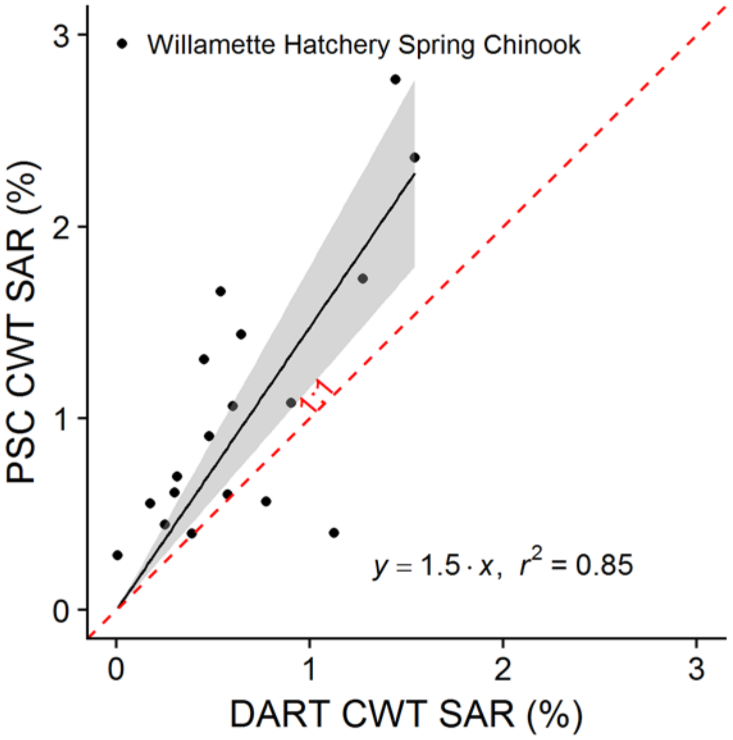


**Figure S3-3**. Comparative SAR of CWT and DART tag-based SAR estimates for yearling Chinook salmon with matched stocks and years of outmigration. The red dashed line shows the expected 1:1 relationship if both DART and CWT-based survival estimates were equivalent. The higher survival for the PSC's CWT-based SARs is in accord with expectation, as harvest is incorporated into the PSC's SAR estimates but not the DART-based estimates.

**References**

McCann, J., B. Chockley, E. Cooper, B. Hsu, H. Schaller, S. Haeseker, R. Lessard, C. Petrosky, T. Copeland, E. Tinus, E. V. Dyke, A. Storch and D. Rawding (2017). Comparative Survival Study of PIT-tagged Spring/Summer/Fall Chinook, Summer Steelhead, and Sockeye. 2017 Annual Report. Portland, Oregon.

Skalski, J. R. and R. L. Townsend (2005). Pacific Northwest Hatcheries Smolt-To-Adult Ratio (SAR) Estimation Using Coded Wire Tags (CWT) Data. Portland, OR, U.S. Department of Energy, Bonneville Power Administration. **Prepared by: Columbia Basin Research, School of Aquatic and Fishery Sciences, University of Washington, Seattle, WA. Project No. 1991-051-00; Contract No. 00013690. :** 13 pp.
